# E484K as an innovative phylogenetic event for viral evolution: Genomic analysis of the E484K spike mutation in SARS-CoV-2 lineages from Brazil

**DOI:** 10.1101/2021.01.27.426895

**Authors:** Patrícia Aline Gröhs Ferrareze, Vinícius Bonetti Franceschi, Amanda de Menezes Mayer, Gabriel Dickin Caldana, Ricardo Ariel Zimerman, Claudia Elizabeth Thompson

## Abstract

The COVID-19 pandemic caused by SARS-CoV-2 has affected millions of people since its beginning in 2019. The propagation of new lineages and the discovery of key mechanisms adopted by the virus to overlap the immune system are central topics for the entire public health policies, research and disease management. Since the second semester of 2020, the mutation E484K has been progressively found in the Brazilian territory, composing different lineages over time. It brought multiple concerns related to the risk of reinfection and the effectiveness of new preventive and treatment strategies due to the possibility of escaping from neutralizing antibodies. To better characterize the current scenario we performed genomic and phylogenetic analyses of the E484K mutated genomes sequenced from Brazilian samples in 2020. From October, 2020, more than 40% of the sequenced genomes present the E484K mutation, which was identified in three different lineages (P1, P2 and B.1.1.33) in four Brazilian regions. We also evaluated the presence of E484K associated mutations and identified selective pressures acting on the spike protein, leading us to some insights about adaptive and purifying selection driving the virus evolution.

## Introduction

The etiological agent of COVID-19, named SARS-CoV-2, belongs to the *Coronaviridae* family composed of enveloped positively oriented single-stranded RNA viruses (Zhu et al., 2020). The first cases of the disease were reported in Wuhan province, China, in December 2019. Since then the number of infected people has increased exponentially worldwide and the World Health Organization (WHO) declared it a pandemic on 11 March 2020. As of 24 January, 2021, the number of confirmed cases globally has scaled up to 99 million, with over 2 million deaths. Brazil is the third country in total number of confirmed cases, already exceeding 8.8 million infected people and 216,445 deaths (Johns Hopkins Coronavirus Resource Center, 2021).

Nearly 410,000 genomes sequenced by researchers from several countries are available on the Global Initiative on Sharing All Influenza Data (GISAID) (Shu and McCauley, 2017), which is crucial to investigate the SARS-CoV-2 genomic epidemiology. A standardized nomenclature was proposed to reflect genetic characteristics and the viral geographical spread patterns. Two ancestral lineages, A and B, have been described containing sublineages that are distinguished by different recurrent mutations and phylogenetic profiles (Rambaut et al., 2020a). More recently special attention has been directed to the understanding of aspects that might impact the virulence and transmissibility of SARS-CoV-2 (Korber et al., 2020; Li, 2020; Toyoshima et al., 2020; Volz et al., 2020) and the influence of different mutations in the effectiveness of COVID-19 vaccines and immunotherapies (Andreano et al., 2020; Pinto et al., 2020; Yu et al., 2020).

Some viral lineages have been studied in more detail, mainly those carrying mutations in the spike (S) glycoprotein, since it is the binding site to the human ACE2 (hACE2) receptor, an essential step to invade the host cell. Currently, there are three lineages of major worldwide concern: B.1.1.7, B.1.351, and P.1. The former emerged in England in mid-September 2020 and it is characterized by 14 lineage-specific amino acid substitutions and has been rapidly spread across the UK and Europe ever since its first appearance (Rambaut et al., 2020b). Two substitutions present in this lineage deserve special attention: N501Y in the Receptor Binding Domain (RBD) of S1 and P681H near the polybasic RRAR sequence in the furin-like cleavage region. N501Y is one of the key contact residues interacting with hACE2 and P681H is one of four residues comprising the insertion that creates a furin-like cleavage site between S1 and S2, which is not found in closely-related coronaviruses (Xia et al., 2020). The second lineage probably emerged in South Africa in August 2020 and harbors three mutations in RBD: K417N, E484K, and N501Y (Tegally et al., 2020). The most recent lineage is P.1, derived from B.1.1.28, which was recently identified in returning travelers from Manaus (Amazonas state, Brazil) who arrived in Japan. It has almost the same three mutations present in RBD as the South African lineage, except for the substitution in amino acid site 417, where the original lysine (K) is substituted for a threonine (T), instead of an asparagine (N). Its appearance is likely to have arisen independently (Faria et al., 2021). In Brazil, the E484K mutation (glutamic acid to lysine substitution at amino acid 484) also arose independently and was identified in Rio de Janeiro state (Southeast Brazil) in early-October carried by the P.2 lineage (Voloch et al., 2020).

Currently, the E484K mutation is found in several viral genomes from Brazil. Due to its rapid spread, the independent origins and the potential implications in vaccination and passive immune therapies, E484K has received particular attention ever since. Importantly, B.1.351, P.1, and P.2 carry this substitution associated with escape from neutralizing antibodies (Baum et al., 2020; Greaney et al., 2020; Weisblum et al., 2020). Recently, this mutation was also identified in a sample from a reinfected patient (Nonaka et al., 2021). These two events are evidence of the virus’ potential for immune escape.

Moreover, B.1.1.7, B.1.351, and P.1 also harbor N501Y mutation, which is associated with enhanced receptor affinity (Starr et al., 2020) and increased infectivity and virulence in a mouse model (Gu et al., 2020). The presence of E484K and N501Y substitutions in the same SARS-CoV-2 genome may be particularly relevant for viral evolution. This combination was shown to induce more conformational changes than the N501Y mutant alone, potentially altering antibodies’ complementarity to this region resulting in the above mentioned immune escape phenomena (Nelson et al., 2021).

Structural analysis points to E484K as potentially the most crucial mutation so far. It creates a new site for the amino acid 75 hACE-2 binding. This interaction seems even stronger than the binding between hACE-2 and the original main site located at position 501 (at RBD and hACE-2 interface) (Nelson et al., 2021). We speculate that the consequent neutralization escape due to E484K alone or as a part of a larger array of distinct mutations might act as a common evolutionary solution for several different viral lineages. Due to the independent origin of two current known lineages harboring E484K in Brazil, we aimed to describe the mutation patterns of these lineages, investigate phylogenetic relationships and perform positive selection tests to identify if adaptive evolution has acted as a major evolutionary force leading to the increase of amino acid variability in the RBD sites.

## Results

### Comparative Genomic Analyses

Firstly detected in the South African B.1.351 lineage, the S1 protein mutation E484K is now present in new emerging variants from Brazil. The analysis of genomes containing E484K, downloaded from the GISAID database, showed a distribution of 169 amino acid residues corresponding to nonsynonymous mutations and four amino acid residue deletions in 134 Brazilian samples (Table S1) collected between October and December 2020 (Figure S1). Regarding synonymous mutations, 217 genome positions had transitions (Figure 1 and Table S2) and 113 genome positions had associated transversions (Figure 1 and Table S2). The arising of E484K mutated genomes with at least four different associated mutation patterns for lineages P.1, P.2, and B.1.1.33 can be seen over time in the aligned genomes (Figure S1 and Table S1).

**Figure 1.**
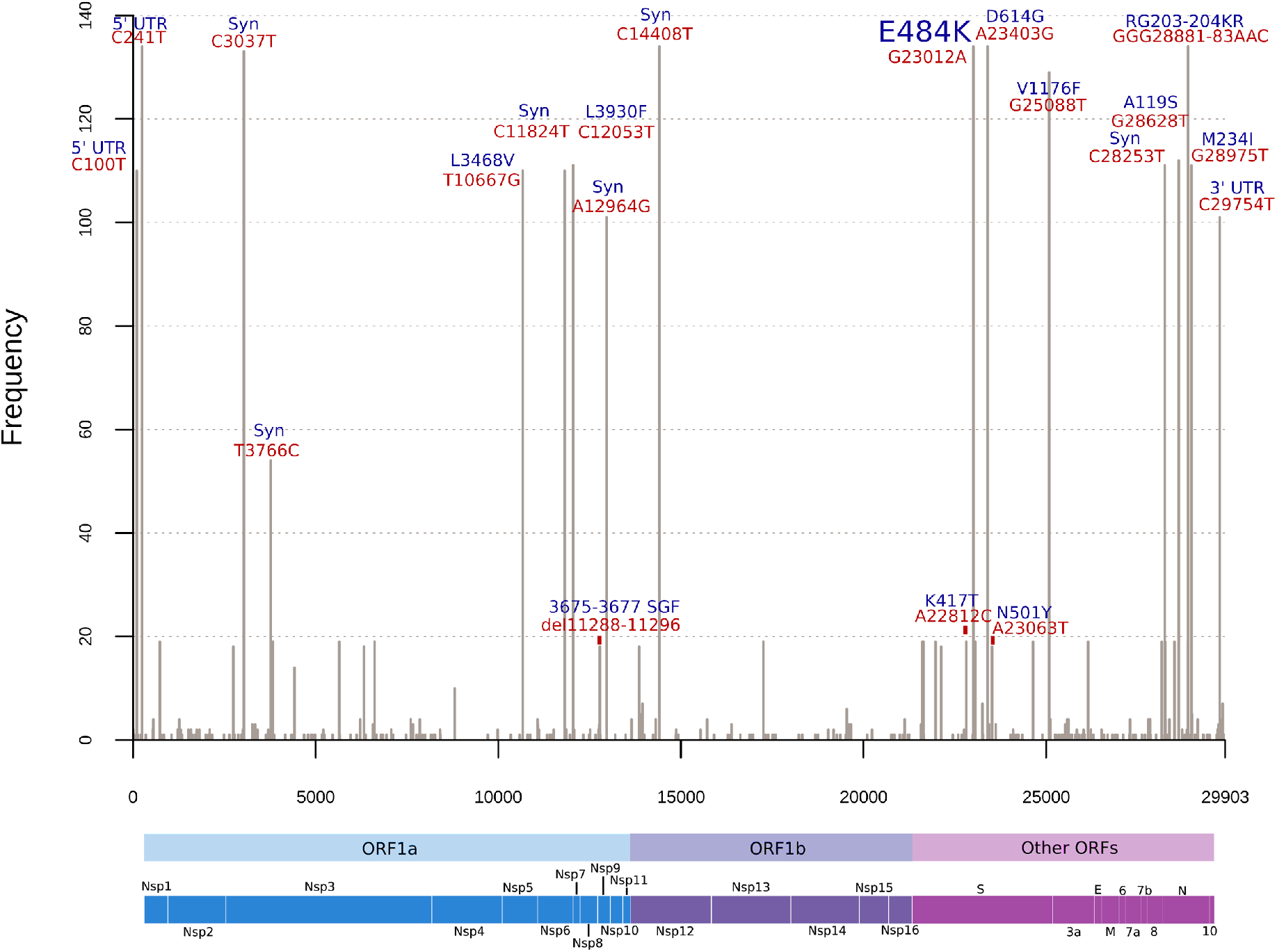
Histogram of frequent mutations observed in the Brazilian SARS-CoV-2 genomes harboring E484K mutation. Red labels above the bars indicate absolute nucleotide position and the blue labels indicate effects of these mutations in the corresponding proteins. As P.1 has only 19 genomes represented and multiple mutations, only main mutations of concern were highlighted. UTR: Untranslated region; Syn: Synonymous substitution; del: deletion; ORF: Open Reading Frame; Nsp: Non-structural protein; S: Spike; E: Envelope; M: Membrane; N: Nucleocapsid.

B.1.1 defining mutations were widespread in almost all sequences as expected (*e. g.* S:D614G, N:R203K, N:G204R, 5’UTR: C241T, and synonymous substitutions in nucleotide positions C3037T and C14408T) (Figure 1). As B.1.1.28 is the ancestral lineage of P.1 and P.2, sequences from these three lineages harbor C12053T (Nsp7:L71F) and G25088T (S:V1176F) mutations. Four sequences classified as B.1.1.33 carry its lineage-defining mutations (T27299C (ORF6: I33T) and T29148C (N: I292T)), but have also five missense mutations in ORF1ab and one in ORF7b (E33A). This pattern indicates a new lineage derived from B.1.1.33, which also possesses the E484K replacement, as P.1 and P.2 sequences. A hallmark of these Brazilian lineages is the presence of multiple lineage-defining mutations per lineage (Table 1).

**Table 1.** Lineage-defining mutations of each of the three Brazilian lineages carrying the E484K mutation. These mutations do not necessarily reflect pangolin lineage assignment defining-mutations, but were extracted based on their representativeness in the majority of sequences of each lineage (https://github.com/cov-lineages/pangolin).

| Lineage / Mutation | B.1.1.33 | P.1 | P.2 |
| --- | --- | --- | --- |
| **5' UTR** | - | - | C100T |
| **ORF1ab** | G1264T (Nsp2)<br>C6573T (Nsp3: S2103F)<br>C7600T (Nsp3)<br>C7851T (Nsp3: A2529V)<br>T11078C (Nsp6: F3605L)<br>C19602T (Nsp14: T6446I)<br>G19656T (Nsp15: R6464M) | T733C (Nsp1)<br>C2749T (Nsp3)<br>C3828T (Nsp3: S1188L)<br>A5648C (Nsp3: K1795Q)<br>A6319G (Nsp3)<br>A6613G (Nsp3)<br>del11288-11296 (Nsp6: 3675-3677 SGF)<br>C12778T (Nsp9)<br>C13860T (Nsp12: T4532I)<br>G17259T (Nsp13: S5665I) | T10667G (Nsp5: L3468V)<br>C11824T (Nsp6)<br>A12964G (Nsp9) |
| **S** | **G23012A (E484K)** | C21614T (L18F)<br>C21621A (T20N)<br>C21638T (P26S)<br>G21974T (D138Y)<br>G22132T (R190S)<br>A22812C (K417T)<br>**G23012A (E484K)**<br>A23063T (N501Y)<br>C23525T (H655Y)<br>C24642T (T1027I) | **G23012A (E484K)** |
| **ORF3a** | - | T26149C (S253P) | - |
| **E** | - | - | - |
| **M** | - | - | - |
| **ORF6** | - | - | - |
| **ORF7a** | - | - | - |
| **ORF7b** | A27853C (E33A) | - | - |
| **ORF8** | - | G28167A (E92K)<br>insG28262GAACA (ORF8) | C28253T (F120F) |
| **N** | - | C28512G (P80R)<br>AGTAGGG28877-83TCTA<br>AAC (RG203KR) | G28628T (A119S)<br>G28975T (M234I) |
| **ORF10** | - | - | - |
| **3' UTR** | C29722T | - | C29754T |

ORF1ab mutations are represented by its amino acid positions relative to ORF1a (Nsp1-Nsp11) and ORF1b (Nsp12-Nsp16). ins: insertion.

The number of genetic changes associated with each E484K Brazilian lineage is highly diverse. B.1.1.33 (E484K) carries a mean of 19.2 (range: 17-22) mutations (considering single nucleotide polymorphisms, insertion and deletion as a single event) in its genome, while P.1 possesses on average 30.1 changes (range: 24-33). Despite harboring a lower number of mutations (mean: 18.5), P.2 genomes have the highest standard deviation (SD=2.2) and range (15-25) of these lineages. These data suggest that both P.1 and P.2 have been circulating in Brazil for a longer period and might be fastly evolving.

The analysis of the E484K mutated sequence EPI_ISL_832010, early detected in the municipality of Esteio, Rio Grande do Sul, shows a simpler set of nonsynonymous mutations (n=10) when compared to others. This genome combines all the prevalent substitutions D614G from spike protein, N:R203K, N:G204R, and Nsp12:P323L, allied to other very frequent mutations (S:V1176F, N:A119S, N:M234I, Nnp5:L205V, and Nsp7:L71F). The presence of spike S:D614G, N:R203K, N:G204R, and Nsp12:P323L in all sequenced E484-containing genomes was observed.

The last samples associated with the recent public health crisis in the state of Amazonas showed an expressive increase in the number of divergences (26 nonsynonymous substitutions and 3 amino acid deletions - Table S1) when compared with the SARS-CoV-2 reference genome (NC_045512.2). Regarding the lineage-defining mutations from P.1 lineages, 14 of a total of 19 genomes from the Amazonas monophyletic group present the spike mutations L18F, T20N, P26S, D138Y, and R190S. In this way, the spike protein of P.1 lineages in these Brazilian E484K mutated genomes is characterized by the presence of a variable number of modified sites, without the fixation of all known mutations. In fact, the P.1 and P2. clades are part of a larger monophyletic group, separated from the B.1.1.33 and the reference genome. These results corroborate the clusterization of these genomes in different lineages (Figure 2 and Figure S2).

**Figure 2.**
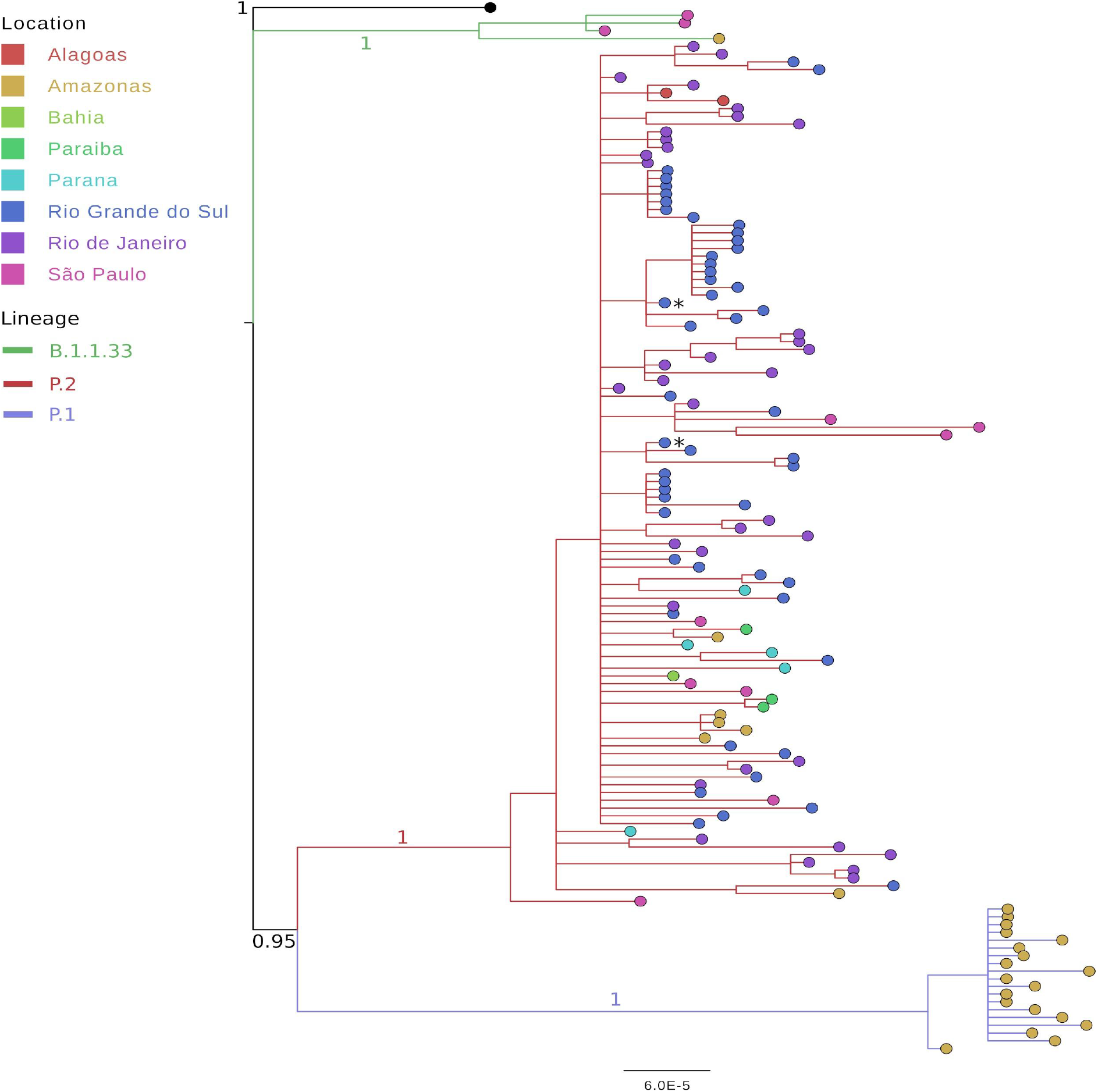
Bayesian phylogenetic inference of the 134 Brazilian E484K mutated genomes. Tips were colored by Brazilian state and the reference genome NC_045512.2 is represented in black. Branches were colored by lineage. The branch values in key branches indicate posterior probabilities.

Lineage-defining mutations as S:K471T, S:N501Y, S:T1027I, N:P80R, Nsp6:S106-107del, Nsp6:F108del, and NS8:E92K were found only in P.1 group and reported for all 19 P.1 sequences. Other mutations such as S:V1176F were not found only in B.1.1.33 lineage. There are also those mutations that are not known as lineage markers but were found in all lineages (n=19) as Nsp3:K977Q, Nsp13:E341D, and NS3:S253P. New specific single substitutions (S:A27V, N:T16M, N:P151L, N:A267V, Nsp13:T216N, and Nsp14:P443S) were also evaluated. Specifically, S:A27V is located in the N-terminal domain.

The B.1.1.33 lineage carrying the E484K mutation were found in São Paulo (n=3) and Amazonas (n=1), while P.1 sequences were found only in the Amazonas state (n=19). All other sequences (n=111) belong to P.2 lineage, the most widespread lineage considering sequenced data available so far. The Northeast region was represented by sequences from Alagoas (n=2), Paraiba (n=3), and Bahia (n=1). The North region is represented by Amazonas sequences (n=6). Southeast sequenced 44 P.2 genomes, 36 from Rio de Janeiro and 8 from São Paulo. Southern Brazil classified 55 genomes as P.2, 5 from Parana and 50 from Rio Grande do Sul, reinforcing its probable dissemination to all regions of Brazil (Figure 3).

**Figure 3.**
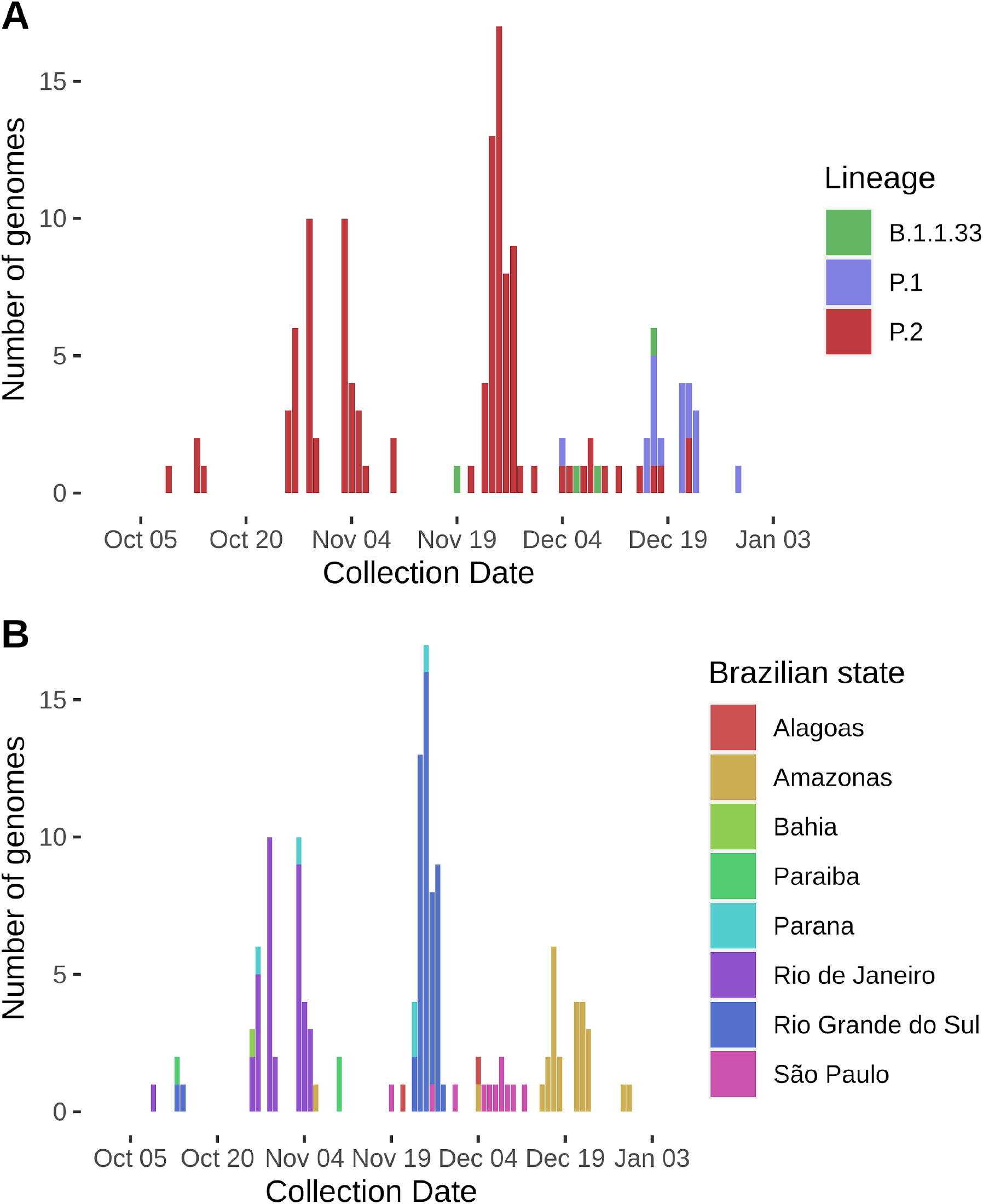
Distribution of genomes harboring E484K mutation across different lineages (A) and Brazilian states (B) from October to December 2020.

The worldwide emergence of E484K began in March 2020, with three sequences represented first. A significant increase was observed in October (n=86), followed by successive increases in November (n=366) and December (n=374) (Figure 4A). A similar trend was observed in Brazil, as E484K was observed in sequences obtained from October (n=31), November (n=87), and December (n=40). Most importantly, the proportion of genomes carrying this replacement was 39.7%, 43.9%, and 43.5% from October to December in comparison to all genomes sequenced from Brazil (Figure 4B). We believe this apparent slow-growing pattern in Brazil is due to heterogeneous and limited initiatives of sequencing in the country, and probably this substitution is already widespread through Brazilian states, as its harbored by three distinct and apparently independent evolving lineages.

**Figure 4.**
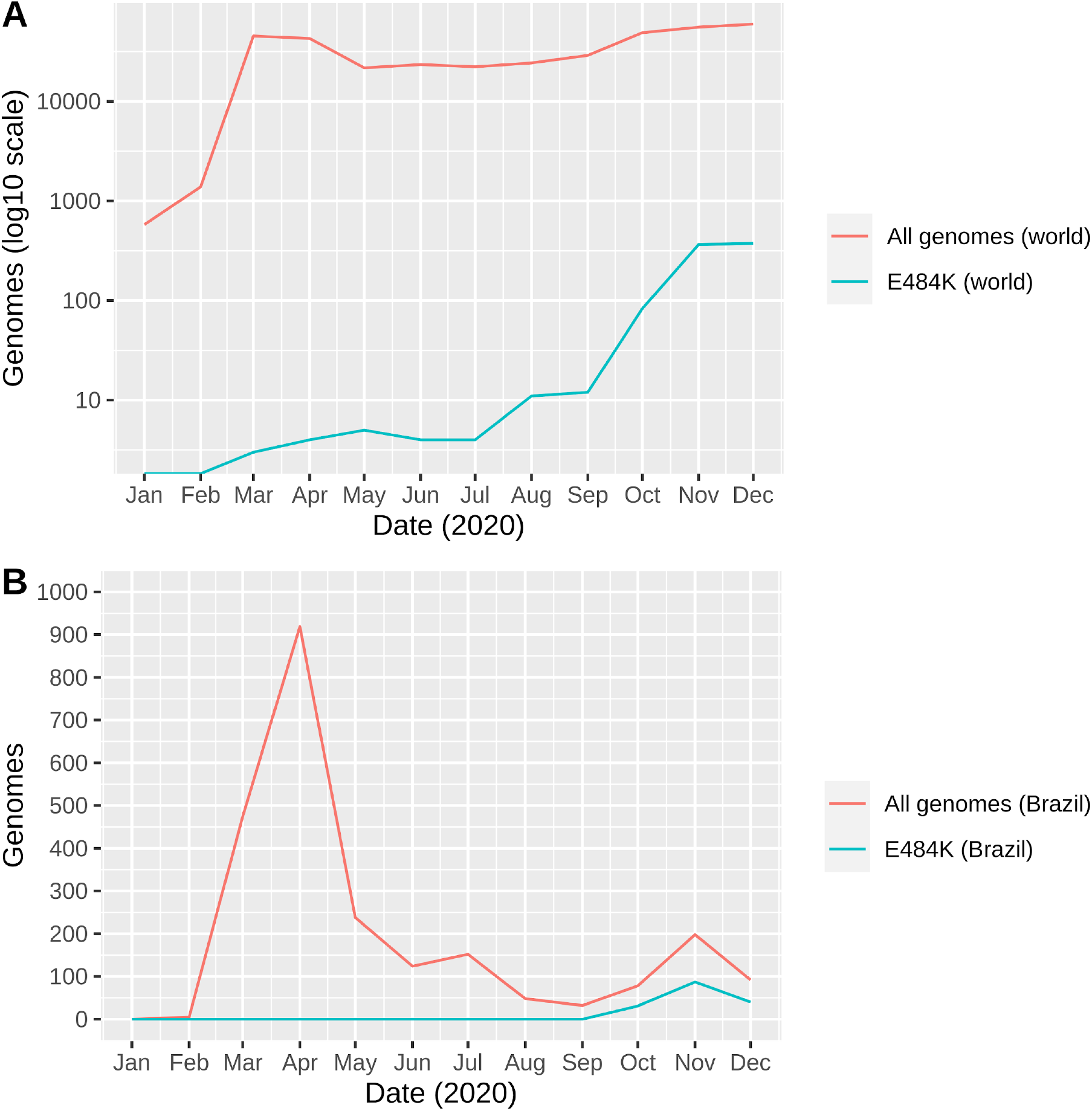
Monthly presence of the E484K mutation considering worldwide available data (A) and Brazilian genomes (B). For clarity, the number of genomes in (A) are represented in log10 scale.

### Selection Analysis

In order to obtain a reliable detection of sites submitted to positive/adaptive or negative/purifying selection, a random set of Brazilian genomes was tested with different approaches. Regarding individual site models, the Bayesian inference FUBAR (Fast, Unconstrained Bayesian AppRoximation) identified eight sites evolving under adaptive selection (positive selection) in the spike sequence (Table 2), with calculated synonymous and nonsynonymous average rates of 1.227 and 0.857, respectively. For these residues under adaptive pressure, six are included in known mutation sites of spike protein, including E484K (L5F, S12F, P26S, D138Y, A688V).

**Table 2.** FUBAR site table for positively selected sites.

| Site | $\alpha$ | $\beta$ | Prob[$\alpha < \beta$] |
| --- | --- | --- | --- |
| 5 | 1.880 | 20.275 | 0.9559 |
| 12 | 1.896 | 21.520 | 0.9584 |
| 26 | 1.879 | 20.481 | 0.9564 |
| 138 | 2.300 | 19.109 | 0.9214 |
| 155 | 2.266 | 17.643 | 0.9158 |
| 484 | 3.850 | 25.932 | 0.9109 |
| 677 | 2.931 | 19.933 | 0.9083 |
| 688 | 1.917 | 23.771 | 0.9628 |

FUBAR inferred the sites submitted to diversifying positive selection with a posterior probability $\geq 0.9$ .

The analysis with the codeml program from the PAML package confirmed all the positively selected sites predicted by FUBAR, also indicating a highly variable selective pressure among sites by the M3 vs. M0 (p < 0.001) comparison and positive selection by the M2 *vs.* M1 and M8 *vs.* M8a model comparison (p < 0.05) (Table 3, 4 and 5). The substitution calculations of M3 model reached a kappa value (ts/tv) of 2.82630 and a site proportion of 93.156% with an estimated omega (ω) value of 0.25424 (negative selection), while 3.536% of the sites were estimated to ω = 4.90450 (positive selection) (Table 3). The ω estimative (ω *= d_N_/d_S_*) is a common method to detect positive selection, since it assumes that synonymous substitutions are neutral and the nonsynonymous are subject to selection. Consequently, a ω statistically higher than 1 would indicate the action of positive selection or a relaxed selective constraint, whereas low *d_N_/d_S_* values would mean conservation of the gene product due to purifying selection (Tennessen, 2008). Additionally to the FUBAR results, three sites were also considered with ω > 1 (Table 3 and 5): 222, 626 and 1263. Of these, only site 222 is not related to the E484K-presenting lineages (A626S and P1263L/S are known). The list of positively selected sites identified by M8 model is available on table S4.

**Table 3.**
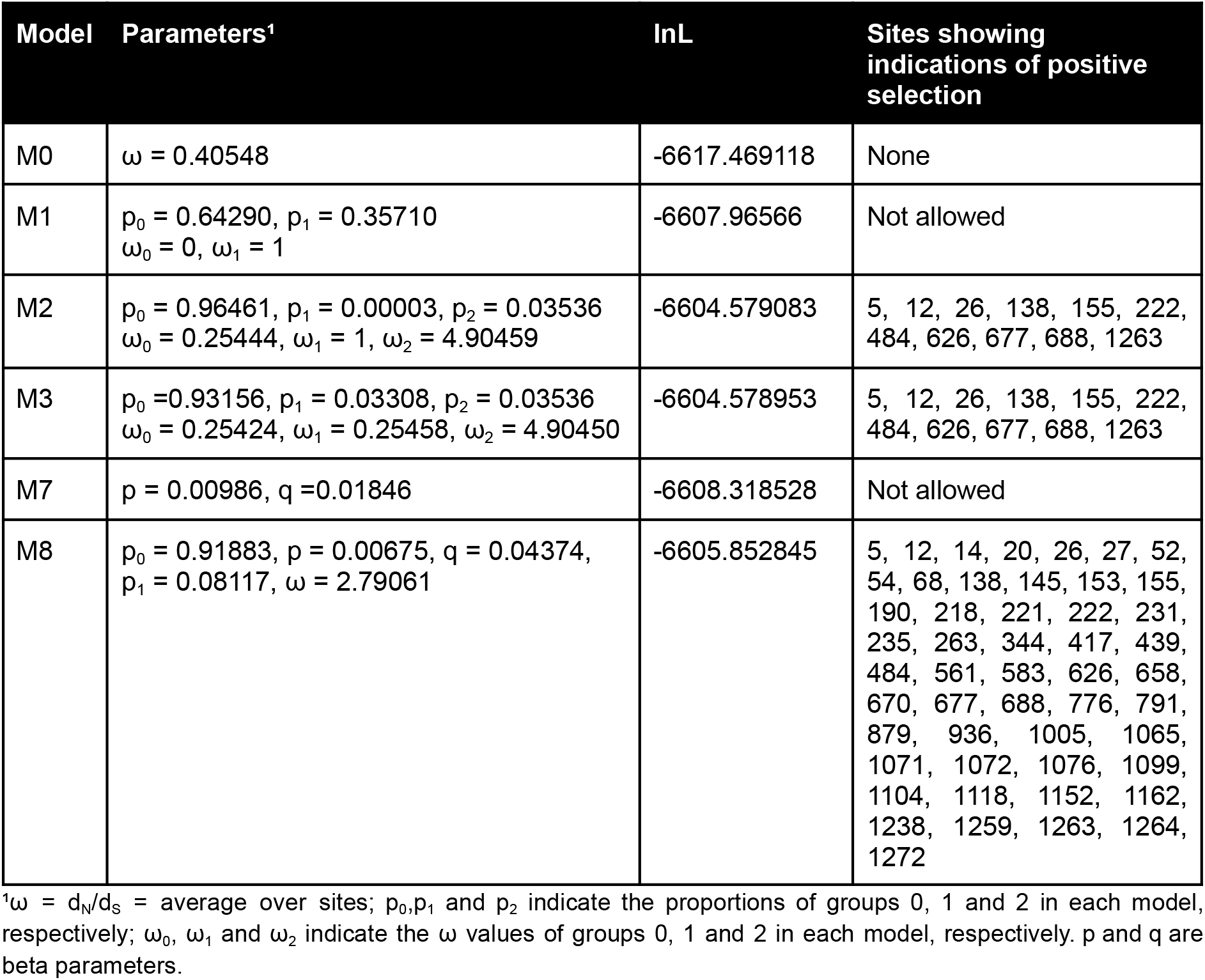
PAML (codeml): Parameters estimates and log-likelihood values under models of variable ω ratios among sites

**Table 4.**
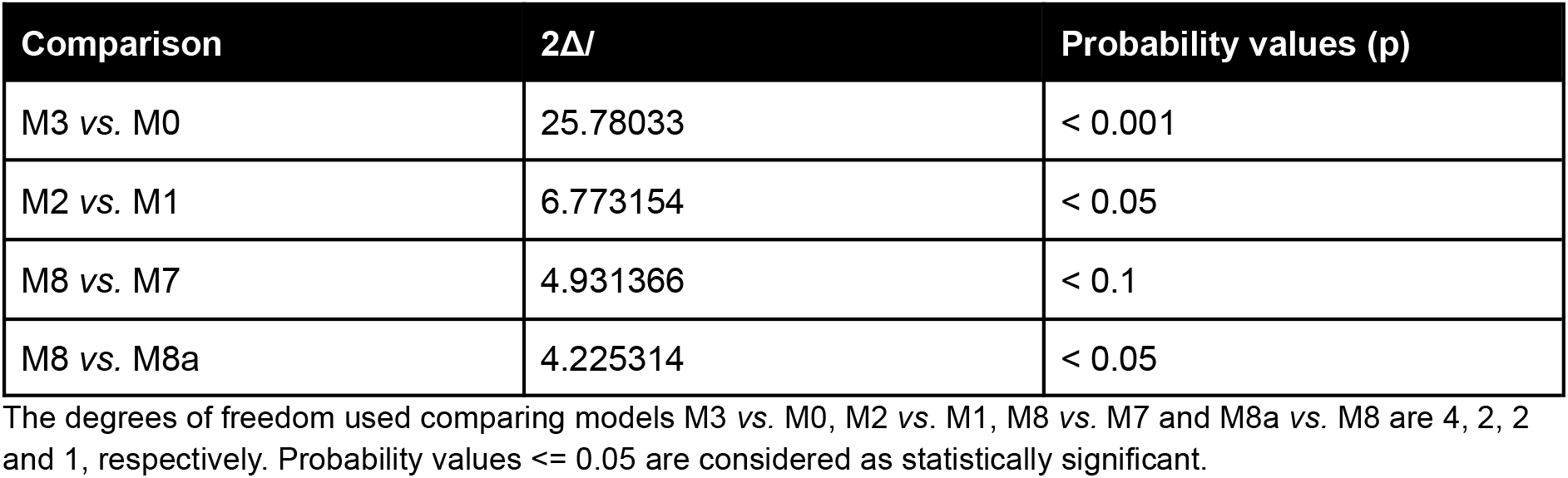
PAML (codeml): Likelihood ratio statistics (2Δ/) for some comparisons between selection models

| Comparison | $2\Delta$ | Probability values (p) |
| --- | --- | --- |
| M3 vs. M0 | 25.78033 | < 0.001 |
| M2 vs. M1 | 6.773154 | < 0.05 |
| M8 vs. M7 | 4.931366 | < 0.1 |
| M8 vs. M8a | 4.225314 | < 0.05 |

The degrees of freedom used comparing models M3 vs. M0, M2 vs. M1, M8 vs. M7 and M8a vs. M8 are 4, 2, 2 and 1, respectively. Probability values $\leq 0.05$ are considered as statistically significant.

**Table 5.** PAML (codeml) site table for positively selected sites

| Site | NEB probabilities |  | BEB probabilities |  |
| --- | --- | --- | --- | --- |
| | Prob ($\omega > 1$) | post mean for $\omega$ | Prob ($\omega > 1$) | post mean ± SE for $\omega$ |
| 5 L | 0.988* | 4.850 | 0.892 | 2.153 ± 0.516 |
| 12 S | 0.988* | 4.849 | 0.892 | 2.153 ± 0.517 |
| 26 P | 0.989* | 4.854 | 0.894 | 2.156 ± 0.514 |
| 138 D | 0.758 | 3.779 | 0.741 | 1.922 ± 0.719 |
| 155 S | 0.782 | 3.893 | 0.749 | 1.935 ± 0.712 |
| 222 A | 0.775 | 3.857 | 0.746 | 1.930 ± 0.714 |
| 484 E | 0.841 | 4.163 | 0.770 | 1.968 ± 0.692 |
| 626 A | 0.860 | 4.252 | 0.778 | 1.980 ± 0.685 |
| 677 Q | 0.909 | 4.483 | 0.802 | 2.017 ± 0.659 |
| 688 A | 0.985* | 4.835 | 0.886 | 2.145 ± 0.525 |
| 1263 P | 0.880 | 4.348 | 0.787 | 1.994 ± 0.675 |

M2 selection model: Naive Empirical Bayes (NEB) analysis. Positively selected sites (\*: $P > 95\%$ ; \*\*: $P > 99\%$ ). Bayes Empirical Bayes (BEB).

To avoid the potential missense effects of the function lost by the sequence mutation, the purifying selection acts as a protective model. The FEL (Fixed Effects Likelihood) approach detected 19 sites under negative selection (Table 6). Three of them (189, 191, and 564) are near residues presenting known nonsynonymous mutations in the E484K mutated genomes, as S:R190S and S:F565L. The SLAC (Single-Likelihood Ancestor Counting) method identified four sites (55, 856, 943, and 1215), two of which were already predicted by FEL. The prevalence of negative selection across the spike protein is consistent with the low genome-wide mutation rate inferred for SARS-CoV-2 (van Dorp et al., 2020).

**Table 6.** FEL site table for negatively selected sites.

| Site | $\alpha$ | $\beta$ | LRT | Prob[$\alpha > \beta$] |
| --- | --- | --- | --- | --- |
| 55 | 18.379 | 0.000 | 4.058 | 0.0440 |
| 91 | 27.071 | 0.000 | 2.792 | 0.0947 |
| 132 | 121.449 | 0.000 | 4.395 | 0.0360 |
| 180 | 121.798 | 0.000 | 4.401 | 0.0359 |
| 189 | 33.258 | 0.000 | 5.432 | 0.0198 |
| 191 | 120.557 | 0.000 | 4.393 | 0.0361 |
| 266 | 27.216 | 0.000 | 2.794 | 0.0946 |
| 324 | 118.682 | 0.000 | 4.370 | 0.0366 |
| 428 | 26.763 | 0.000 | 2.852 | 0.0913 |
| 475 | 16.429 | 0.000 | 3.268 | 0.0707 |
| 564 | 121.798 | 0.000 | 4.159 | 0.0414 |
| 821 | 21.710 | 0.000 | 3.065 | 0.0800 |
| 897 | 42.768 | 0.000 | 4.241 | 0.0395 |
| 910 | 42.680 | 0.000 | 3.305 | 0.0691 |
| 1120 | 43.397 | 0.000 | 3.598 | 0.0578 |
| 1126 | 26.763 | 0.000 | 3.495 | 0.0616 |
| 1215 | 18.327 | 0.000 | 2.861 | 0.0907 |
| 1228 | 42.759 | 0.000 | 4.136 | 0.0420 |
| 1251 | 42.680 | 0.000 | 3.305 | 0.0691 |

Sites identified under negative selection at $p \leq 0.1$ . Grey rows: significant sites also identified by SLAC.

## Discussion

The permanence of a pathogen inside a population host depends on its efficiency in key processes such as the replication, the scaping of the immune system and the binding to the cell receptor. Mutations that create an advantageous scenario towards the host response to infection enhance the pathogen fitness under the natural selection pressure. Consequently, the host-pathogen coevolution represents an important mechanism to understand the establishment and the prognostics of pathogenesis, since the infections are possibly the major selective pressure acting on humans (Sironi et al., 2015). Therefore, the circulation of low and moderate pathogens provides time for the pathogen adaptations, in such a way that the modulation of the immunity may possibly promote molecular convergence in different lineages over the time (Longdon et al., 2014).

The presence of spike S:D614G, N:R203K, N:G204R, and Nsp12:P323L in all sequenced E484 mutated samples, reaching three different lineages, might suggest that these lineages show increased viral replication. Considering that E484K enhances escape from immune system antibodies, these may potentially lead to a viral advantage. The occurrence of simultaneous mutations as N:R203K and N:G204R is already known in the SARS-CoV-2 literature. However, the fixation of these mutations, as well as Nsp12:P323L and D614G in all the E484K evaluated genomes may indicate a novel adaptive relationship among these modifications resulting in viral evolutionary success.

Even as an independent evolutionary event, the potential fixation of mutations such as E484K across lineages may indicate active mechanisms of adaptive selection and are very relevant in planning future therapeutic strategies (for example, newer vaccines and immunotherapy platforms). The deletion of three amino acids in the second helix of the transmembrane protein Nsp6 in P.1 genomes may affect the virus-induced cellular autophagy and the formation of double-membrane vesicles for the viral RNA synthesis (Benvenuto et al., 2020).

The important role of subsequent stabilization of the flexible NTD by mutations has been speculated (Laha et al., 2020). Typically, NTD can harbor a larger number of evolutionary events than RBD, including mutations, insertions and deletions that could act allosterically altering the binding affinities between RBD and hACE-2 and inducing immune evasion. It is known that distinct positions may have a linked relationship in the final protein structure and may display some advantage by acting together to achieve increased stability, adaptability, viability and/or transmission efficiency (Laha et al., 2020). The potential causality or influence of the E484K and others substitutions for the effectiveness of neutralizing antibodies that bind the N-terminal domain of the spike protein (Chi et al., 2020) remains uncertain.

Concerning the selection analysis, the FUBAR method was the first one to be tested. According to Murrell (2013), the FUBAR method may have more detection power than methods like FEL, in particular when positive selection is present but relatively weak. The analysis of the South African clade V501.V2 (https://observablehq.com/@spond/zav501v2#table1) found some similar results for FUBAR evaluation, with the detection of adaptive selection at the sites 5 (L5F), 12 (S12F), and 484 (E484K). The residues 155 and 677 are not associated with the E484K-presenting lineages, however, recent data found evidence of evolutionary convergence of this mutation in at least six distinct sub-lineages, which could improve proteolytic processing, cell tropism, and transmissibility (Hodcroft et al., 2021). The amino acid residue 677 is next to the furin-like cleavage site (678 to 688). Interestingly, some studies suggest that the emergence of the SARS-CoV-2 in the human species resulted from a five amino acid change in the critical S glycoprotein binding site (Fam et al., 2020).

To the best of our knowledge, the impact of E484K in different lineages have not been deeply explored. It was structurally demonstrated that, at least in combination with K417N and N501Y, the substitution has profound impact in shifting the main site of contact between viral RBD and hACE-2 residues (Nelson et al., 2021). However, we still do not know if this holds true for other sets of mutations associated with E484K.

In summary, we have demonstrated widespread dissemination of mutants harboring E484K replacement in geographically diverse regions in Brazil. This substitution was disseminated in our country as early as October 2020, although phylodynamics inferences placed its emergence in July (Voloch et al., 2020). The fact that E484K was found in the context of different mutations and lineages is suggestive that this particular substitution may act as a common solution for viral evolution in different genotypes. This hypothesis may be related to the profound impact of the mutation, which changes a negatively charged amino acid (glutamic acid) for a positively charged amino acid (lysine). Since this position is present in a highly flexible loop, it has been proposed that the presence of such mutation could create a strong ionic interaction between lysine (K) in RBD and amino acid 75 of the hACE-2 receptor shifting the major sites of binding in S1 from positions 497-502 to 484 (Nelson et al., 2021).

Mutations in RBD could theoretically decrease neutralizing activity of serum of patients receiving vaccines against SARS-CoV-2 with unknown clinical consequences. Arguably the effect of E484K could be particularly relevant. In a recent publication, this mutation was associated with complete abolishment of all neutralizing activity in a high proportion of convalescent serum tested (Wibmer et al., 2021). When taken together these growing body of evidence suggest that E484K should be the target of intense virologic surveillance. Studies testing the activity of serum from vaccinated patients against viruses or pseudoviruses with the aforementioned substitution should be considered a high public health priority. Second generation immune therapies and vaccines focusing on more conserved domains (for instance, in S2 fusion domain) may deserve special attention to assure continuous therapeutic efficacy.

## Methods

### Sequencing Data Retrieval

For the graphic evaluation of Brazilian lineages, all genomes available on the GISAID database until January 18^th^, 2021, and presenting the E484K mutation were included in the analysis. Genomes from Brazil and other countries submitted until January 24^th^, 2021, and with collected dates between January 1^st^ and December 31^st^, 2020, were selected. The counting of genomes containing E484K was performed by the specification of the mutation in the search fields. The monthly count was defined by the values between the first and the last day of each month.

### Comparative Genomic Analyses

The comparative genomic analysis of the E484K mutated genomes was performed with all the 134 genomes (Table S1) presenting the E484K mutation for Brazil in GISAID until January 18^th^, 2021. For these, a genomic multiple sequence alignment was performed on the MAFFT web server (Katoh and Standley, 2013) using default options and 1PAM = k/2 scoring matrix. Single nucleotide polymorphisms (SNPs) and insertions/deletions (INDELs) were assessed by using snippy variant calling pipeline v4.6.0 (https://github.com/tseemann/snippy). Mutations were concatenated with associated metadata and counted by Brazilian states using custom Python scripts. The mapping of the aligned sequences to the reference genome (NC_045512.2) features was created by the software GENEIOUS 2021.0.3 (https://www.geneious.com). Histogram of mutations was generated using a modified code from Lu et al. 2020 (https://github.com/laduplessis/SARS-CoV-2_Guangdong_genomic_epidemiology/).

### Phylogenetic Analysis

The previously aligned genomic sequences were used as input for the phylogenetic analysis. The reference genome NC_045512.2 was added as an outgroup. The inference of the best evolutionary model was performed by ModelTest-NG (Darriba et al., 2020), which identified GTR+I (Generalised time-reversible model + proportion of invariant sites). The tree was built by Bayesian inference in MrBayes v3.2.7 (Huelsenbeck and Ronquist, 2001) with the GTR+I model and 5 million generations. The data convergence was evaluated with Tracer v1.7.1 (Rambaut et al., 2018) and tree visualization was created by FigTree software (http://tree.bio.ed.ac.uk/software/figtree/).

For the phylogenetic analysis using the spike protein, the multiple sequence alignment previously obtained was edited and only the spike coding sequences were maintained. The inference of the evolutionary model and tree construction were performed according to the described model: ModelTest-NG model selection, MrBayes with GTR+I and 10 million generations.

### Selection analysis

In total, 780 SARS-CoV-2 genomes, collected from Brazil samples between February 1^st^ and December 31^st^, 2020, available on GISAID, were used for this analysis. After an initial filtering step of undefined nucleotides in the region of spike protein, with deletion of genomes with a N ratio >0.01, 589 genomes were selected. The second step of subsampling, with Augur toolkit (Huddleston et al., 2021) kept 498 genomes (approximately 119 representative genomes per lineage) for the multiple sequence alignment (Table S3). The genomic multiple sequence alignment was performed by MAFFT (Katoh and Standley, 2013) with default options and 1PAM = k/2 scoring matrix. The selection of the genome location for the spike protein was performed with the software UGENE (Okonechnikov et al., 2012), using the ORF coordinates of the NC_045512 reference genome as parameter. GTR+I was inferred by ModelTest-NG (Darriba et al., 2020) as the best evolutionary model for the spike sequences. ModelTest-NG also indicated identical sequences, which were removed resulting in 161 sequences. The Bayesian phylogenetic inference was performed by MrBayes v3.2.7 using those 161 unique spike selected sequences and 5 million generations. The data convergence was evaluated with Tracer v1.7.1.

The analysis of sites under positive and negative selection was performed by HyPhy v2.5.23 (Pond et al., 2005) according to different approaches: (i) FUBAR (Unconstrained Bayesian AppRoximation) (Murrell et al., 2013), (ii) FEL (**F**ixed **E**ffects **L**ikelihood) (Kosakovsky Pond and Frost, 2005), and (iii) SLAC (**S**ingle-**L**ikelihood **A**ncestor **C**ounting) (Kosakovsky Pond and Frost, 2005). Finally, the PAML (Phylogenetic Analysis by Maximum Likelihood) (Yang, 2007) program was used for confirmation of possibly selected sites with the codeml package. All models were run using the F3×4 option in the PAML program, where expected codon frequencies were based upon nucleotide frequencies occurring at the three codon positions. The one-ratio model (M0) assumes one ω ratio for all sites. The neutral model (M1) presupposes a proportion p_0_ of conserved sites with ω_0_ = 0 and p_1_ = 1 - p_0_ of neutral sites with ω_1_ = 1, as would occur if almost all non-synonymous substitutions were either deleterious or neutral. The positive selection model (M2) adds an additional class of sites with frequency p_2_ = 1 - p_0_ - p_1_ and ω_2_ is estimated from the data. In the discrete model (M3), the probabilities (p_0_, p_1_ and p_2_) of each site which was submitted to purifying selection, neutral selection and positive selection, respectively, and their corresponding w ratios (ω_0_, ω_1_, ω_2_) are inferred from the data. The beta model (M7) is a null test for positive selection, assuming a beta distribution with ω between 0 and 1. By last, the beta & ω (M8) model adds one extra class with the same ratio ω_1_. The LRTs (Likelihood Ratio Tests) were tested to investigate whether ω was significantly different from 1 for each pairwise comparison: M1a *vs*. M2a, M0 *vs*. M3, and M7 *vs*. M8. LRT performs the comparison both with the constraint of ω=1 and without such constraint: LRT = 2 (ln_1_ - ln_2_). These LRT statistics approximately follow a chi-square distribution and the number of degrees of freedom is equal to the number of additional parameters in the more complex model. For rejection of the null hypothesis of neutrality, we considered p ≤ 0.05. Finally, we applied the Naive Empirical Bayes (NEB) and Bayes Empirical Bayes (BEB) approaches available in the PAML package to calculate the posterior probability that each site belongs to the positively selected class.

## Competing interest statement

The authors declare no competing interests.

## Funding

Scholarships and Fellowships were supplied by the Coordenação de Aperfeiçoamento de Pessoal de Nível Superior – Brasil (CAPES) – Finance Code 001 and Universidade Federal de Ciências da Saúde de Porto Alegre. The funders had no role in the study design, data generation and analysis, decision to publish or the preparation of the manuscript.

## Acknowledgements

We thank the administrators and curators of the GISAID database and research groups across the globe for supporting the rapid and transparent sharing of genomic data during the COVID-19 pandemic. We also thank the Mayor’s Office, Health Department and São Camilo Hospital (Esteio, RS, Brazil), Leonardo Duarte Pascoal and Ana Regina Boll for their work in combating Covid-19 and for supporting the work developed by our research group. A full table acknowledging the authors and corresponding labs submitting sequencing data used in this study can be found in Supplementary Files 1 and 3.

## Author contributions

C.E.T. and R.A.Z. conceived the study. C.E.T., P.A.G.F., V.B.F. contributed bioinformatics tools. C.E.T., P.A.G.F., R.A.Z., V.B.F. performed the data analysis. A.M.M., C.E.T., G.D.C., P.A.G.F., R.A.Z., V.B.F. wrote the manuscript.

## Supplementary material

**Tables**

**Table S1.** Brazilian sequenced genomes containing the E484K mutation.

**Table S2.** Single Nucleotide Polymorphisms identified in the multiple genomes comparison.

**Table S3.** Brazilian sequenced genomes used for selection analysis (499 sequences).

**Figure S1.**
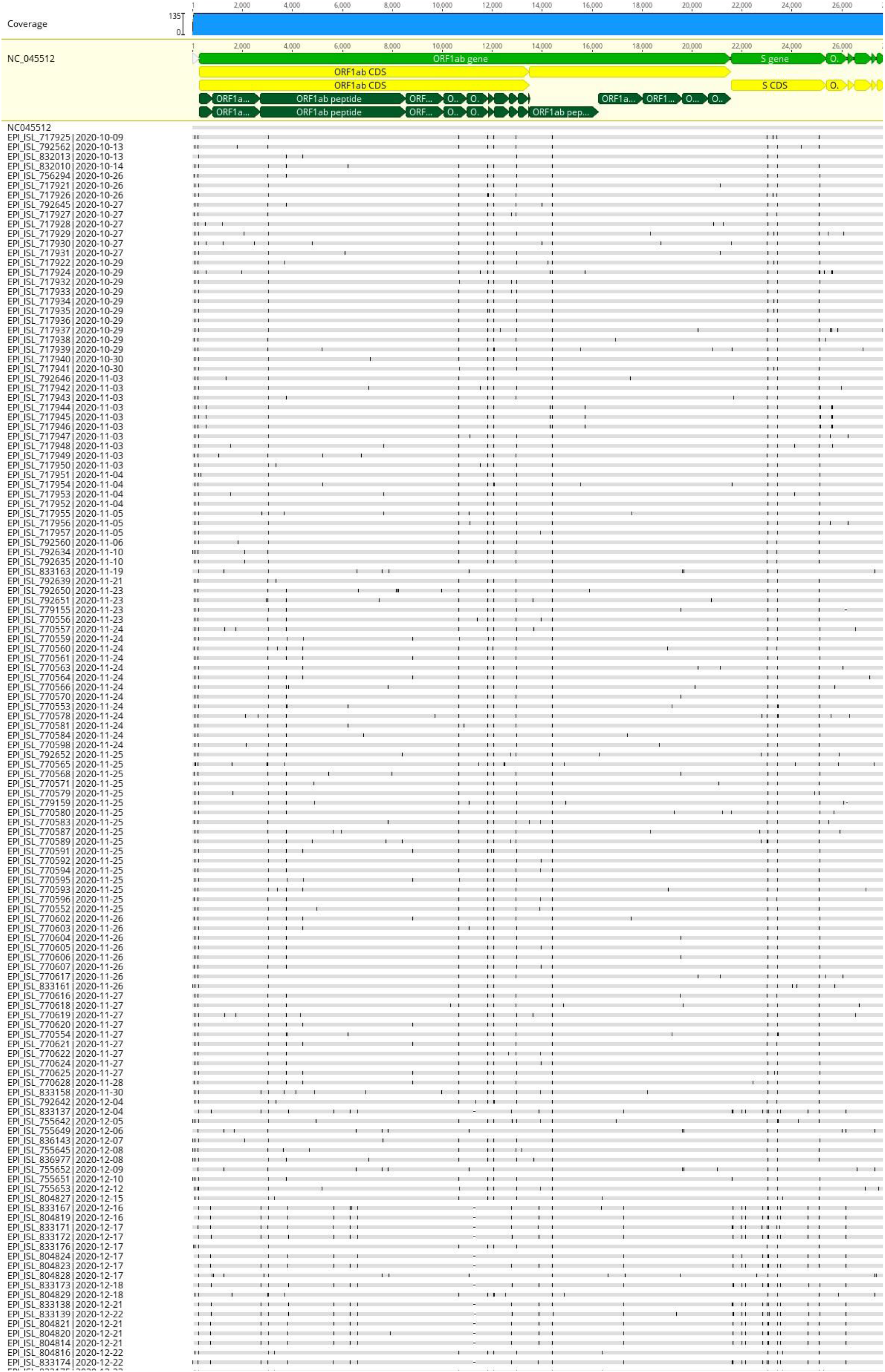
Multiple sequence alignment (time-ordered) of the 134 sequenced brazilian genomes with the E484K mutation. Genomes retrieved from GISAID until January 18^th^, 2020.

**Figure S2.**
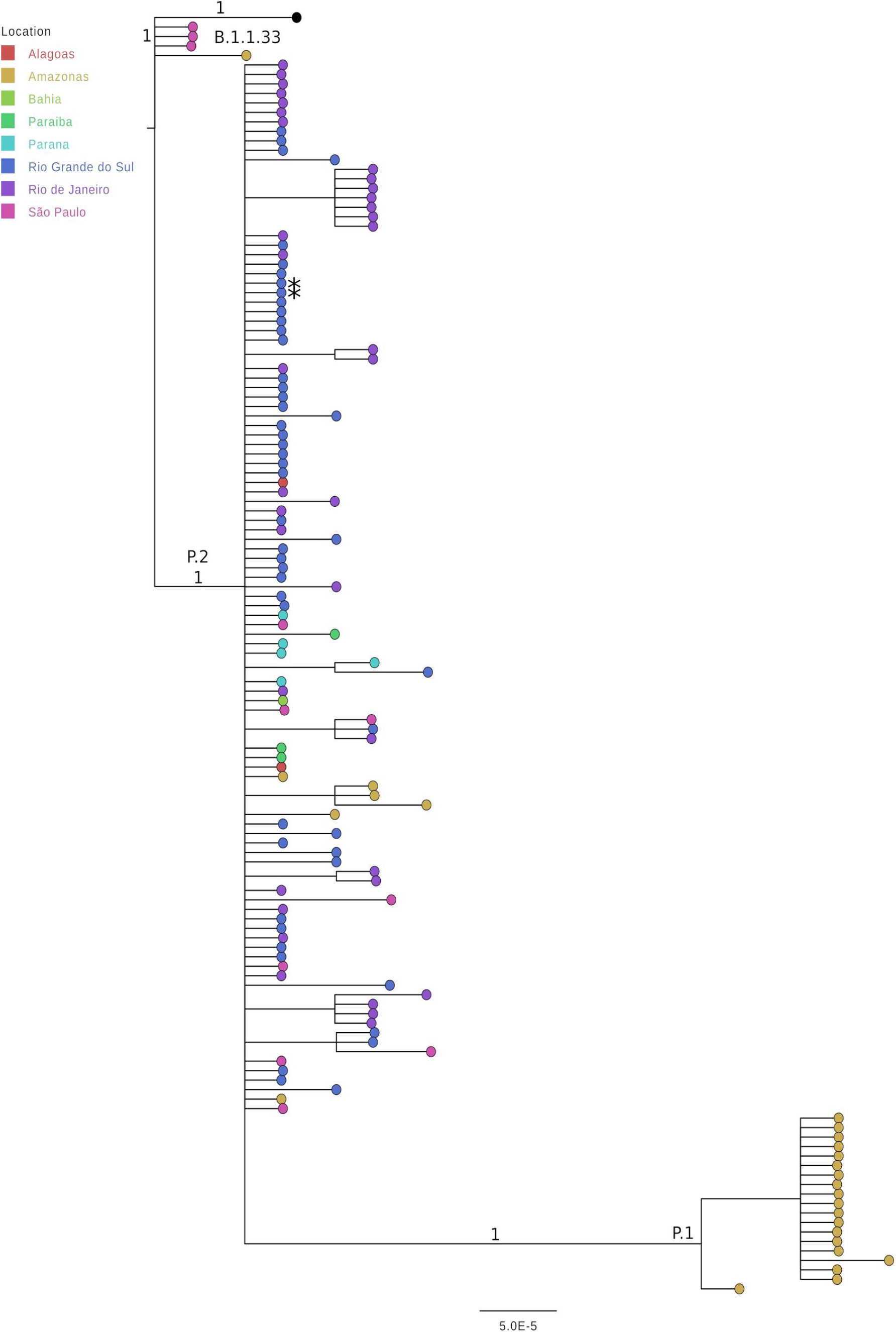
Phylogenetic analysis of the 134 Spike protein sequences with the E484K mutation from Brazilian genomes. Tree inferred by MrBayes using the GTR+IN evolutionary model. Values in key clades indicate posterior probabilities and lineages.

**Figure S3.**
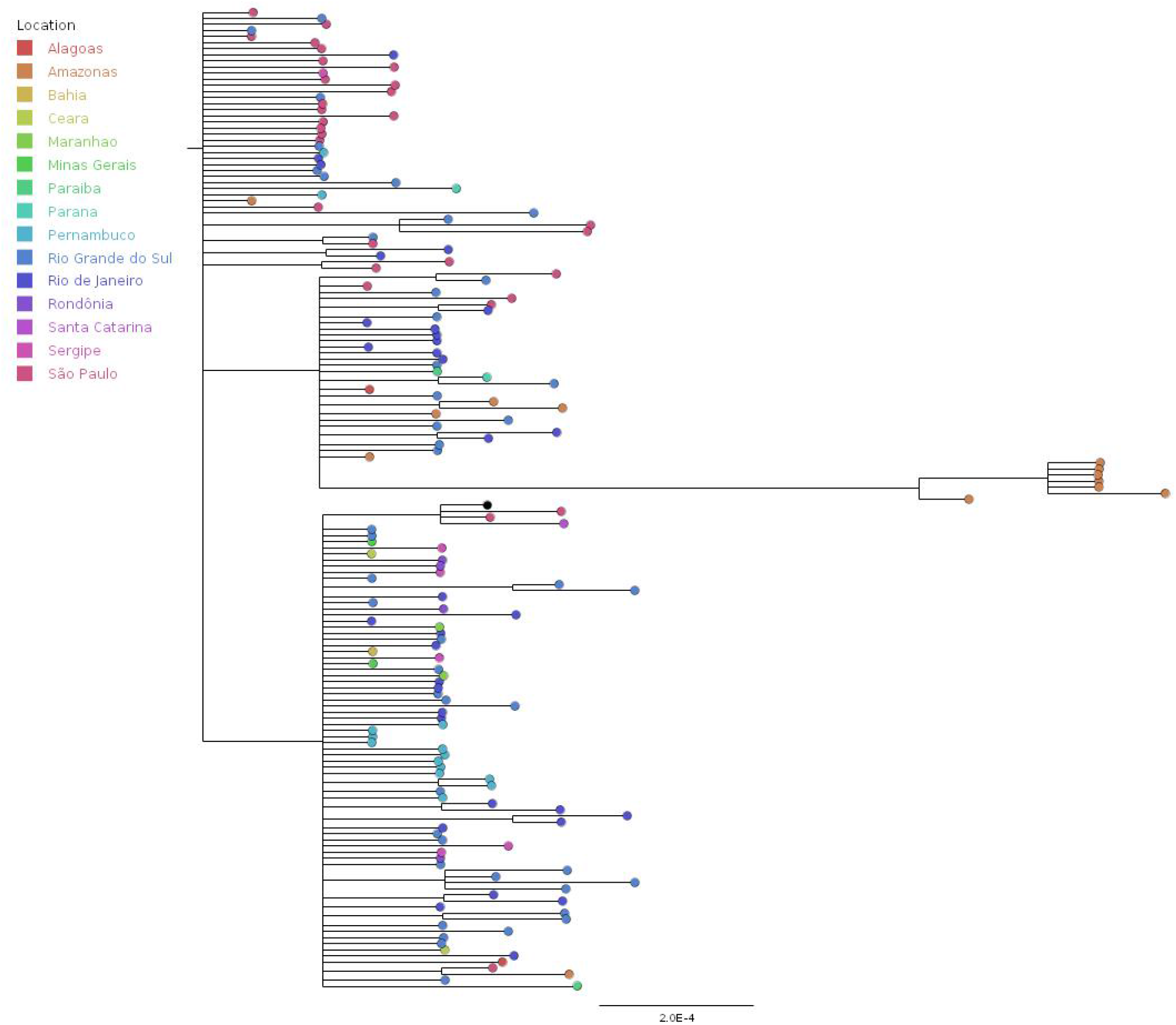
Phylogenetic analysis of the 161 unique Spike protein sequences from Brazilian genomes. Tree inferred by MrBayes using the GTR+I evolutionary model.

## Notes

### Competing Interest Statement

The authors have declared no competing interest.

### Summary of Updates

Detailed description of the positive selection results from PAML analysis.

## References

Andreano, E., Piccini, G., Licastro, D., Casalino, L., Johnson, N.V., Paciello, I., Monego, S.D., Pantano, E., Manganaro, N., Manenti, A., Manna, R., Casa, E., Hyseni, I., Benincasa, L., Montomoli, E., Amaro, R.E., McLellan, J.S., Rappuoli, R., (2020). SARS-CoV-2 escape in vitro from a highly neutralizing COVID-19 convalescent plasma. bioRxiv 2020.12.28.424451.

Baum, A., Fulton, B.O., Wloga, E., Copin, R., Pascal, K.E., Russo, V., Giordano, S., Lanza, K., Negron, N., Ni, M., Wei, Y., Atwal, G.S., Murphy, A.J., Stahl, N., Yancopoulos, G.D., Kyratsous, C.A., (2020). Antibody cocktail to SARS-CoV-2 spike protein prevents rapid mutational escape seen with individual antibodies. Science 369, 1014–1018.

Benvenuto, D., Angeletti, S., Giovanetti, M., Bianchi, M., Pascarella, S., Cauda, R., Ciccozzi, M., Cassone, A., (2020). Evolutionary analysis of SARS-CoV-2: how mutation of Non-Structural Protein 6 (NSP6) could affect viral autophagy. J. Infect. 81, e24–e27.

Chi, X., Yan, R., Zhang, Jun, Zhang, G., Zhang, Y., Hao, M., Zhang, Z., Fan, P., Dong, Y., Yang, Y., Chen, Z., Guo, Y., Zhang, Jinlong, Li, Y., Song, X., Chen, Y., Xia, L., Fu, L., Hou, L., Xu, J., Yu, C., Li, J., Zhou, Q., Chen, W., (2020). A neutralizing human antibody binds to the N-terminal domain of the Spike protein of SARS-CoV-2. Science 369, 650–655.

Darriba, D., Posada, D., Kozlov, A.M., Stamatakis, A., Morel, B., Flouri, T., (2020). ModelTest-NG: A New and Scalable Tool for the Selection of DNA and Protein Evolutionary Models. Mol. Biol. Evol. 37, 291–294.

Fam, Bibiana S.O., Vargas-Pinilla, Pedro, Amorim, Carlos Eduardo G., Sortica, Vinicius A., & Bortolini, Maria Cátira., (2020). ACE2 diversity in placental mammals reveals the evolutionary strategy of SARS-CoV-2. Genetics and Molecular Biology 43(2), e20200104. Epub June 08, 2020.

Faria, N., Claro, I.M., Candido, D., Franco, L.A.M., Andrade, P.S., Coletti, T.M., Silva, C.A.M., Fraiji, N.A., Esashika Crispim, M.A., Carvalho, M. do P.S.S., Rambaut, A., Loman, N., Pybus, O.G., Sabino, E.C., (2021). Genomic characterisation of an emergent SARS-CoV-2 lineage in Manaus: preliminary findings [WWW Document]. Virological. URL https://virological.org/t/genomic-characterisation-of-an-emergent-sars-cov-2-lineage-in-manaus-preliminary-findings/586 (accessed 1.14.21).

Greaney, A.J., Starr, T.N., Gilchuk, P., Zost, S.J., Binshtein, E., Loes, A.N., Hilton, S.K., Huddleston, J., Eguia, R., Crawford, K.H.D., Dingens, A.S., Nargi, R.S., Sutton, R.E., Suryadevara, N., Rothlauf, P.W., Liu, Z., Whelan, S.P.J., Carnahan, R.H., Crowe, J.E., Bloom, J.D., (2021). Complete mapping of mutations to the SARS-CoV-2 spike receptor-binding domain that escape antibody recognition. Cell Host Microbe 29(1), 44–57.e9.

Gu, H., Chen, Q., Yang, G., He, L., Fan, H., Deng, Y.-Q., Wang, Y., Teng, Y., Zhao, Z., Cui, Y., Li, Yuchang, Li, X.-F., Li, J., Zhang, N.-N., Yang, Xiaolan, Chen, S., Guo, Y., Zhao, G., Wang, X., Luo, D.-Y., Wang, H., Yang, Xiao, Li, Yan, Han, G., He, Y., Zhou, X., Geng, S., Sheng, X., Jiang, S., Sun, S., Qin, C.-F., Zhou, Y., (2020). Adaptation of SARS-CoV-2 in BALB/c mice for testing vaccine efficacy. Science 369, 1603–1607.

Hodcroft, E.B., Domman, D.B., Oguntuyo, K., Snyder, D.J., Van Diest, M., Densmore, K.H., Schwalm, K.C., Femling, J., Carroll, J.L., Scott, R.S., Whyte, M.M., Edwards, M.D., Hull, N.C., Kevil, C.G., Vanchiere, J.A., Lee, B., Dinwiddie, D.L., Cooper, V.S., Kamil, J.P., (2021). Emergence in late 2020 of multiple lineages of SARS-CoV-2 Spike protein variants affecting amino acid position 677. medRxiv 2021.02.12.21251658.

Huddleston, J., Hadfield, J., Sibley, T.R., Lee, J., Fay, K., Ilcisin, M., Harkins, E., Bedford, T., Neher, R.A., Hodcroft, E.B., (2021). Augur: a bioinformatics toolkit for phylogenetic analyses of human pathogens. J. Open Source Softw. 6, 2906.

Huelsenbeck, J.P., Ronquist, F., (2001). MRBAYES: Bayesian inference of phylogenetic trees. Bioinformatics 17, 754–755.

Johns Hopkins Coronavirus Resource Center, n.d. COVID-19 Map [WWW Document]. Johns Hopkins Coronavirus Resour. Cent. URL https://coronavirus.jhu.edu/map.html (accessed 11.10.20).

Katoh, K., Standley, D.M., (2013). MAFFT Multiple Sequence Alignment Software Version 7: Improvements in Performance and Usability. Mol. Biol. Evol. 30, 772–780.

Korber, B., Fischer, W.M., Gnanakaran, S., Yoon, H., Theiler, J., Abfalterer, W., Hengartner, N., Giorgi, E.E., Bhattacharya, T., Foley, B., Hastie, K.M., Parker, M.D., Partridge, D.G., Evans, C.M., Freeman, T.M., de Silva, T.I., Angyal, A., Brown, R.L., Carrilero, L., Green, L.R., Groves, D.C., Johnson, K.J., Keeley, A.J., Lindsey, B.B., Parsons, P.J., Raza, M., Rowland-Jones, S., Smith, N., Tucker, R.M., Wang, D., Wyles, M.D., McDanal, C., Perez, L.G., Tang, H., Moon-Walker, A., Whelan, S.P., LaBranche, C.C., Saphire, E.O., Montefiori, D.C., (2020). Tracking Changes in SARS-CoV-2 Spike: Evidence that D614G Increases Infectivity of the COVID-19 Virus. Cell 182, 812–827.e19.

Kosakovsky Pond, S.L., Frost, S.D.W., (2005). Not So Different After All: A Comparison of Methods for Detecting Amino Acid Sites Under Selection. Mol. Biol. Evol. 22, 1208–1222.

Laha, S., Chakraborty, J., Das, S., Manna, S.K., Biswas, S., Chatterjee, R., (2020). Characterizations of SARS-CoV-2 mutational profile, spike protein stability and viral transmission. Infect. Genet. Evol. 85, 104445.

Li, S., (2020). Modifiable lifestyle factors and severe COVID-19 risk: Evidence from Mendelian randomization analysis. medRxiv 2020.10.19.20215525.

Longdon B., Brockhurst M.A., Russell C.A., Welch J.J., Jiggins F.M., (2014). The Evolution and Genetics of Virus Host Shifts. PLoS Pathog 10(11): e1004395.

Lu, J., du Plessis, L., Liu, Z., Hill, V., Kang, M., Lin, H., Sun, J., François, S., Kraemer, M.U.G., Faria, N.R., McCrone, J.T., Peng, J., Xiong, Q., Yuan, R., Zeng, L., Zhou, P., Liang, C., Yi, L., Liu, J., Xiao, J., Hu, J., Liu, T., Ma, W., Li, W., Su, J., Zheng, H., Peng, B., Fang, S., Su, W., Li, K., Sun, R., Bai, R., Tang, X., Liang, M., Quick, J., Song, T., Rambaut, A., Loman, N., Raghwani, J., Pybus, O.G., Ke, C., (2020). Genomic Epidemiology of SARS-CoV-2 in Guangdong Province, China. Cell 181, 997–1003.e9.

Murrell, B., Moola, S., Mabona, A., Weighill, T., Sheward, D., Kosakovsky Pond, S.L., Scheffler, K., (2013). FUBAR: A Fast, Unconstrained Bayesian AppRoximation for Inferring Selection. Mol. Biol. Evol. 30, 1196–1205.

Nelson, G., Buzko, O., Spilman, P., Niazi, K., Rabizadeh, S., Soon-Shiong, P., (2021). Molecular dynamic simulation reveals E484K mutation enhances spike RBD-ACE2 affinity and the combination of E484K, K417N and N501Y mutations (501Y.V2 variant) induces conformational change greater than N501Y mutant alone, potentially resulting in an escape mutant. bioRxiv 2021.01.13.426558.

Nonaka, C.K.V., Franco, M.M., Gräf, T., Mendes, A.V.A., Aguiar, R.S. de, Giovanetti, M., Souza, B.S. de F., (2021). Genomic Evidence of a Sars-Cov-2 Reinfection Case With E484K Spike Mutation in Brazil. Preprints, 2021010132.

Okonechnikov, K., Golosova, O., Fursov, M., UGENE team, (2012). Unipro UGENE: a unified bioinformatics toolkit. Bioinforma. Oxf. Engl. 28, 1166–1167.

Pinto, D., Park, Y.-J., Beltramello, M., Walls, A.C., Tortorici, M.A., Bianchi, S., Jaconi, S., Culap, K., Zatta, F., De Marco, A., Peter, A., Guarino, B., Spreafico, R., Cameroni, E., Case, J.B., Chen, R.E., Havenar-Daughton, C., Snell, G., Telenti, A., Virgin, H.W., Lanzavecchia, A., Diamond, M.S., Fink, K., Veesler, D., Corti, D., (2020). Cross-neutralization of SARS-CoV-2 by a human monoclonal SARS-CoV antibody. Nature 583, 290–295.

Pond, S.L.K., Frost, S.D.W., Muse, S.V., (2005). HyPhy: hypothesis testing using phylogenies. Bioinformatics 21, 676–679.

Rambaut, A., Drummond, A.J., Xie, D., Baele, G., Suchard, M.A., (2018). Posterior Summarization in Bayesian Phylogenetics Using Tracer 1.7. Syst. Biol. 67, 901–904.

Rambaut, A., Holmes, E.C., O’Toole, Á., Hill, V., McCrone, J.T., Ruis, C., du Plessis, L., Pybus, O.G., (2020a). A dynamic nomenclature proposal for SARS-CoV-2 lineages to assist genomic epidemiology. Nat. Microbiol. 5, 1403–1407.

Rambaut, A., Loman, N., Pybus, O., Barclay, W., Barrett, J., Carabelli, A., Connor, T., Peacock, T., Robertson, D., Volz, E., (2020b). Preliminary genomic characterisation of an emergent SARS-CoV-2 lineage in the UK defined by a novel set of spike mutations [WWW Document]. Virological. URL https://virological.org/t/preliminary-genomic-characterisation-of-an-emergent-sars-cov-2-lineage-in-the-uk-defined-by-a-novel-set-of-spike-mutations/563 (accessed 1.4.21).

Shu, Y., McCauley, J., (2017). GISAID: Global initiative on sharing all influenza data – from vision to reality. Eurosurveillance 22.

Sironi, M., Cagliani, R., Forni, D. et al., (2015). Evolutionary insights into host–pathogen interactions from mammalian sequence data. Nat Rev Genet 16, 224–236.

Smith, M.D., Wertheim, J.O., Weaver, S., Murrell, B., Scheffler, K., Kosakovsky Pond, S.L., (2015). Less Is More: An Adaptive Branch-Site Random Effects Model for Efficient Detection of Episodic Diversifying Selection. Mol. Biol. Evol. 32, 1342–1353.

Starr, T.N., Greaney, A.J., Hilton, S.K., Ellis, D., Crawford, K.H.D., Dingens, A.S., Navarro, M.J., Bowen, J.E., Tortorici, M.A., Walls, A.C., King, N.P., Veesler, D., Bloom, J.D., (2020). Deep Mutational Scanning of SARS-CoV-2 Receptor Binding Domain Reveals Constraints on Folding and ACE2 Binding. Cell 182, 1295–1310.e20.

Tegally, H., Wilkinson, E., Giovanetti, M., Iranzadeh, A., Fonseca, V., Giandhari, J., Doolabh, D., Pillay, S., San, E.J., Msomi, N., Mlisana, K., Gottberg, A. von, Walaza, S., Allam, M., Ismail, A., Mohale, T., Glass, A.J., Engelbrecht, S., Zyl, G.V., Preiser, W., Petruccione, F., Sigal, A., Hardie, D., Marais, G., Hsiao, M., Korsman, S., Davies, M.-A., Tyers, L., Mudau, I., York, D., Maslo, C., Goedhals, D., Abrahams, S., Laguda-Akingba, O., Alisoltani-Dehkordi, A., Godzik, A., Wibmer, C.K., Sewell, B.T., Lourenço, J., Alcantara, L.C.J., Pond, S.L.K., Weaver, S., Martin, D., Lessells, R.J., Bhiman, J.N., Williamson, C., Oliveira, T. de, (2020). Emergence and rapid spread of a new severe acute respiratory syndrome-related coronavirus 2 (SARS-CoV-2) lineage with multiple spike mutations in South Africa. medRxiv 2020.12.21.20248640.

Tennessen, J.A., (2008). Positive selection drives a correlation between non-synonymous/synonymous divergence and functional divergence. Bioinformatics 24(12), 1421–1425.

Toyoshima, Y., Nemoto, K., Matsumoto, S., Nakamura, Y., Kiyotani, K., (2020). SARS-CoV-2 genomic variations associated with mortality rate of COVID-19. J. Hum. Genet. 65, 1075–1082.

van Dorp, L., Acman, M., Richard, D., Shaw, L. P., Ford, C. E., Ormond, L., Owen, C. J., Pang, J., Tan, C., Boshier, F., Ortiz, A. T., & Balloux, F., (2020). Emergence of genomic diversity and recurrent mutations in SARS-CoV-2. Infection, genetics and evolution: journal of molecular epidemiology and evolutionary genetics in infectious diseases 83, 104351.

Voloch, C.M., F, R. da S., Almeida, L.G.P. de, Cardoso, C.C., Brustolini, O.J., Gerber, A.L., Guimarães, A.P. de C., Mariani, D., Costa, R.M. da, Ferreira, O.C., Workgroup, C.-U., LNCC-Workgroup, Cavalcanti, A.C., Frauches, T.S., Mello, C.M.B. de, Galliez, R.M., Faffe, D.S., Castiñeiras, T.M.P.P., Tanuri, A., Vasconcelos, A.T.R. de, (2020). Genomic characterization of a novel SARS-CoV-2 lineage from Rio de Janeiro, Brazil. medRxiv 2020.12.23.20248598.

Volz, E., Hill, V., McCrone, J.T., Price, A., Jorgensen, D., O’Toole, Á., Southgate, J., Johnson, R., Jackson, B., Nascimento, F.F., Rey, S.M., Nicholls, S.M., Colquhoun, R.M., da Silva Filipe, A., Shepherd, J., Pascall, D.J., Shah, R., Jesudason, N., Li, K., Jarrett, R., Pacchiarini, N., Bull, M., Geidelberg, L., Siveroni, I., Goodfellow, I., Loman, N.J., Pybus, O.G., Robertson, D.L., Thomson, E.C., Rambaut, A., Connor, T.R., (2020). Evaluating the effects of SARS-CoV-2 Spike mutation D614G on transmissibility and pathogenicity. Cell 184(1), 64–75.e11.

Weisblum, Y., Schmidt, F., Zhang, F., Da Silva, J., Poston, D., Lorenzi, J.C., Muecksch, F., Rutkowska, M., Hoffmann, H.-H., Michailidis, E., Gaebler, C., Agudelo, M., Cho, A., Wang, Z., Gazumyan, A., Cipolla, M., Luchsinger, L., Hillyer, C.D., Caskey, M., Robbiani, D.F., Rice, C.M., Nussenzweig, M.C., Hatziioannou, T., Bieniasz, P.D., (2020). Escape from neutralizing antibodies by SARS-CoV-2 spike protein variants. eLife 9, e61312.

Wibmer, C.K., Ayres, F., Hermanus, T., Madzivhandila, M., Kgagudi, P., Lambson, B.E., Vermeulen, M., Berg, K. van den, Rossouw, T., Boswell, M., Ueckermann, V., Meiring, S., Gottberg, A. von, Cohen, C., Morris, L., Bhiman, J.N., Moore, P.L., (2021). SARS-CoV-2 501Y.V2 escapes neutralization by South African COVID-19 donor plasma. bioRxiv 2021.01.18.427166.

Xia, S., Lan, Q., Su, S., Wang, X., Xu, W., Liu, Z., Zhu, Y., Wang, Q., Lu, L., Jiang, S., (2020). The role of furin cleavage site in SARS-CoV-2 spike protein-mediated membrane fusion in the presence or absence of trypsin. Signal Transduct. Target. Ther. 5, 1–3.

Yang Z., 2007. PAML 4: phylogenetic analysis by maximum likelihood. Mol Biol Evol. 2007 Aug;24(8):1586–91. doi: 10.1093/molbev/msm088. Epub. May 4. PMID: 17483113.

Yu, F., Xiang, R., Deng, X., Wang, L., Yu, Z., Tian, S., Liang, R., Li, Y., Ying, T., Jiang, S., (2020). Receptor-binding domain-specific human neutralizing monoclonal antibodies against SARS-CoV and SARS-CoV-2. Signal Transduct. Target. Ther. 5, 1–12.

Zhu, N., Zhang, D., Wang, W., Li, X., Yang, B., Song, J., Zhao, X., Huang, B., Shi, W., Lu, R., Niu, P., Zhan, F., Ma, X., Wang, D., Xu, W., Wu, G., Gao, G.F., Tan, W., (2020). A Novel Coronavirus from Patients with Pneumonia in China, 2019. N. Engl. J. Med. 382, 727–733.

