## Supplementary material for "E484K as an innovative phylogenetic event for viral evolution: Genomic analysis of the E484K spike mutation in SARS-CoV-2 lineages from Brazil": Table S1

| Accession id | Collection date | Submission date | Location | Lineage | Mutations |
|---|---|---|---|---|---|
| EPI_ISL_717925 | 9/10/2020 | 20/12/2020 | Rio de Janeiro / Rio de Janeiro | P.2 | Spike D614G, Spike E484K, Spike F565L, Spike V1176F, N A119S, N G204R, N M234I, N R203K, NSP5 L205V, NSP7 L71F, NSP12 P323 |
| EPI_ISL_792562 | 13/10/2020 | 10/1/2021 | Paraiba / Joao Pessoa | P.2 | Spike D614G, Spike E484K, Spike S929I, Spike V1176F, N A119S, N G204R, N M234I, N P151S, N R203K, NSP2 A336T, NSP2 G339S, NSP5 L205V, NSP7 L71F, NSP12 P323L |
| EPI_ISL_832013 | 13/10/2020 | 15/1/2021 | Rio Grande do Sul / Esteio | B.1.1 | Spike D614G, Spike E484K, N A119S, N G204R, N M234I, N R203K, NSP12 P323L |
| EPI_ISL_832010 | 14/10/2020 | 15/1/2021 | Rio Grande do Sul / Esteio | P.2 | Spike D614G, Spike E484K, Spike V1176F, N A119S, N G204R, N M234I, N R203K, NSP5 L205V, NSP7 L71F, NSP12 P323L |
| EPI_ISL_756294 | 26/10/2020 | 2/1/2021 | Bahia/Salvador | P.2 | Spike D614G, Spike E484K, Spike V1176F, N A119S, N G204R, N M234I, N P207S, N R203K, NSP5 L205V, NSP7 L71F, NSP12 P323L |
| EPI_ISL_717921 | 26/10/2020 | 20/12/2020 | Rio de Janeiro / Cabo Frio | P.2 | Spike D614G, Spike E484K, Spike V1176F, N A119S, N G204R, N M234I, N R203K, NSP5 L205V, NSP7 L71F, NSP12 P323L |
| EPI_ISL_717926 | 26/10/2020 | 20/12/2020 | Rio de Janeiro / Duque de Caxias | P.2 | Spike D614G, Spike E484K, Spike F565L, Spike V1176F, N A119S, N G204R, N M234I, N R203K, NS8 I121L, NSP5 L205V, NSP7 L71F, NSP12 P323L |
| EPI_ISL_792645 | 27/10/2020 | 10/1/2021 | Parana / Umuarama | P.2 | Spike D614G, Spike E484K, Spike V1176F, N A119S, N G204R, N M234I, N R203K, NSP5 L205V, NSP7 L71F, NSP12 A185V, NSP12 P323L |
| EPI_ISL_717927 | 27/10/2020 | 20/12/2020 | Rio de Janeiro / Rio de Janeiro | P.2 | Spike D614G, Spike E484K, Spike V1176F, N A119S, N G204R, N M234I, N R203K, NSP5 L205V, NSP7 L71F, NSP12 P323L |
| EPI_ISL_717928 | 27/10/2020 | 20/12/2020 | Rio de Janeiro / Rio de Janeiro | P.2 | Spike D614G, Spike E484K, Spike V1176F, N A119S, N G204R, N M234I, N R203K, NSP2 S138L, NSP5 L205V, NSP7 L71F, NSP12 P323L, NSP16 M65I |
| EPI_ISL_717929 | 27/10/2020 | 20/12/2020 | Rio de Janeiro / Duque de Caxias | P.2 | Spike D614G, Spike E484K, Spike F565L, Spike V1176F, N A119S, N G204R, N M234I, N R203K, NS3 T12I, NSP2 A419V, NSP5 L205V, NSP7 L71F, NSP12 P323L, NSP14 A96V |
| EPI_ISL_717930 | 27/10/2020 | 20/12/2020 | Rio de Janeiro / Rio de Janeiro | P.2 | Spike D614G, Spike E484K, Spike L5F, Spike V1176F, N A119S, N G204R, N M234I, N R203K, NSP2 S138L, NSP3 A690V, NSP5 L205V, NSP7 L71F, NSP12 G179S, NSP12 P323L |
| EPI_ISL_717931 | 27/10/2020 | 20/12/2020 | Rio de Janeiro / Rio de Janeiro | P.2 | Spike D614G, Spike E484K, Spike V1176F, N A119S, N G204R, N M234I, N R203K, NSP3 S1134L, NSP5 L205V, NSP7 L71F, NSP12 P323L |
| EPI_ISL_717922 | 29/10/2020 | 20/12/2020 | Rio de Janeiro / Rio de Janeiro | P.2 | Spike D614G, Spike E484K, Spike F565L, Spike V1176F, N A119S, N G204R, N M234I, N R203K, NS8 I121L, NSP3 A333V, NSP5 L205V, NSP7 L71F, NSP12 P323L |
| EPI_ISL_717924 | 29/10/2020 | 20/12/2020 | Rio de Janeiro / Rio de Janeiro | P.2 | Spike D614G, Spike E484K, Spike T1238I, Spike V1176F, N A119S, N G204R, N M234I, N R203K, NS3 S74C, NSP2 Q395H, NSP5 L205V, NSP6 T181I, NSP7 L71F, NSP12 P323L |
| EPI_ISL_717932 | 29/10/2020 | 20/12/2020 | Rio de Janeiro / Rio de Janeiro | P.2 | Spike D614G, Spike E484K, Spike V1176F, N A119S, N G204R, N M234I, N R203K, NSP5 L205V, NSP7 L71F, NSP12 P323L |
| EPI_ISL_717933 | 29/10/2020 | 20/12/2020 | Rio de Janeiro / Rio de Janeiro | P.2 | Spike D614G, Spike E484K, Spike V1176F, N A119S, N G204R, N M234I, N R203K, NSP5 L205V, NSP7 L71F, NSP12 P323L |
| EPI_ISL_717934 | 29/10/2020 | 20/12/2020 | Rio de Janeiro / Rio de Janeiro | P.2 | Spike D614G, Spike E484K, Spike F565L, Spike V1176F, N A119S, N G204R, N M234I, N R203K, NSP5 L205V, NSP7 L71F, NSP12 P323L |
| EPI_ISL_717935 | 29/10/2020 | 20/12/2020 | Rio de Janeiro / Rio de Janeiro | P.2 | Spike D614G, Spike E484K, Spike F565L, Spike V1176F, N A119S, N G204R, N M234I, N R203K, NS8 I121L, NSP5 L205V, NSP7 L71F, NSP12 P323L |
| EPI_ISL_717936 | 29/10/2020 | 20/12/2020 | Rio de Janeiro / Rio de Janeiro | P.2 | Spike D614G, Spike E484K, Spike V1176F, N A119S, N G204R, N M234I, N R203K, NSP5 L205V, NSP7 L71F, NSP12 P323L |
| EPI_ISL_717937 | 29/10/2020 | 20/12/2020 | Rio de Janeiro / Duque de Caxias | P.2 | Spike D614G, Spike E484K, Spike V1176F, N A119S, N G204R, N M234I, N R203K, NS3 G44R, NS3 N137Y, NSP5 L205V, NSP7 L71F, NSP12 P323L |
| EPI_ISL_717938 | 29/10/2020 | 20/12/2020 | Rio de Janeiro / Niterói | P.2 | Spike D614G, Spike E484K, Spike P1263L, Spike V1176F, N A119S, N G204R, N M234I, N R203K, NSP5 L205V, NSP7 L71F, NSP12 P323L |
| EPI_ISL_717939 | 29/10/2020 | 20/12/2020 | Rio de Janeiro / Rio de Janeiro | P.2 | Spike D614G, Spike E484K, Spike S12F, Spike V1176F, N A119S, N G204R, N M234I, N R203K, NSP3 M829I, NSP5 L205V, NSP7 L71F, NSP12 P323L |
| EPI_ISL_717940 | 30/10/2020 | 20/12/2020 | Rio de Janeiro / Niterói | P.2 | Spike D614G, Spike E484K, Spike V1176F, N A119S, N G204R, N M234I, N R203K, NSP3 I1468T, NSP5 L205V, NSP7 L71F, NSP12 P323L |
| EPI_ISL_717941 | 30/10/2020 | 20/12/2020 | Rio de Janeiro / Rio de Janeiro | P.2 | Spike D614G, Spike E484K, Spike F565L, Spike V1176F, N A119S, N G204R, N M234I, N R203K, NSP5 L205V, NSP7 L71F, NSP12 P323L |
| EPI_ISL_792646 | 3/11/2020 | 10/1/2021 | Parana / Maringa | P.2 | Spike D614G, Spike E484K, Spike V1176F, N A119S, N G204R, N M234I, N R203K, NSP2 L180S, NSP5 L205V, NSP7 L71F, NSP12 P323L |
| EPI_ISL_717942 | 3/11/2020 | 20/12/2020 | Rio de Janeiro / Rio de Janeiro | P.2 | Spike D614G, Spike E484K, Spike V1176F, N A119S, N G204R, N M234I, N R203K, NSP3 G1451C, NSP5 L205V, NSP6 T181I, NSP7 L71F, NSP12 P323L |
| EPI_ISL_717943 | 3/11/2020 | 20/12/2020 | Rio de Janeiro / Rio de Janeiro | P.2 | Spike D614G, Spike E484K, Spike T29I, Spike V1176F, N A119S, N G204R, N M234I, N R203K, NSP5 L205V, NSP7 L71F, NSP12 P323L |
| EPI_ISL_717944 | 3/11/2020 | 20/12/2020 | Rio de Janeiro / Rio de Janeiro | P.2 | Spike D614G, Spike E484K, Spike V1176F, N A119S, N G204R, N M234I, N R203K, NS3 S74C, NSP5 L205V, NSP7 L71F, NSP12 P323L |
| EPI_ISL_717945 | 3/11/2020 | 20/12/2020 | Rio de Janeiro / Rio de Janeiro | P.2 | Spike D614G, Spike E484K, Spike V1176F, N A119S, N G204R, N M234I, N R203K, NS3 S74C, NSP5 L205V, NSP7 L71F, NSP12 P323L |
| EPI_ISL_717946 | 3/11/2020 | 20/12/2020 | Rio de Janeiro / Rio de Janeiro | P.2 | Spike D614G, Spike E484K, Spike V1176F, N A119S, N G204R, N M234I, N R203K, NS3 S74C, NSP5 L205V, NSP7 L71F, NSP12 P323L |
| EPI_ISL_717947 | 3/11/2020 | 20/12/2020 | Rio de Janeiro / Rio de Janeiro | P.2 | Spike D614G, Spike E484K, Spike V1176F, N A119S, N G204R, N M234I, N R203K, NS3 G44R, NSP5 L205V, NSP6 A46V, NSP7 L71F, NSP12 P323L |
| EPI_ISL_717948 | 3/11/2020 | 20/12/2020 | Rio de Janeiro / Rio de Janeiro | P.2 | Spike A846S, Spike D614G, Spike E484K, Spike V1176F, N A119S, N G204R, N M234I, N R203K, NSP2 K239R, NSP5 L205V, NSP7 L71F, NSP12 P323L |

| | | | | | |
|---|---|---|---|---|---|
| EPI_ISL_717949 | 3/11/2020 | 20/12/2020 | Rio de Janeiro / Rio de Janeiro | P.2 | Spike D614G, Spike E484K, Spike V1176F, N A119S, N G204R, N M234I, N R203K, NSP2 D84N, NSP5 L205V, NSP7 L71F, NSP12 P323L |
| EPI_ISL_717950 | 3/11/2020 | 20/12/2020 | Rio de Janeiro / Rio de Janeiro | P.2 | Spike D614G, Spike E484K, Spike V1176F, N A119S, N G204R, N M234I, N R203K, NSP5 L205V, NSP7 L71F, NSP12 P323L |
| EPI_ISL_717951 | 4/11/2020 | 20/12/2020 | Rio de Janeiro / Rio de Janeiro | P.2 | Spike D614G, Spike E484K, Spike V1176F, N A119S, N G204R, N M234I, N R203K, NSP5 L205V, NSP7 L71F, NSP12 P323L |
| EPI_ISL_717952 | 4/11/2020 | 20/12/2020 | Rio de Janeiro / Rio de Janeiro | P.2 | Spike D614G, Spike E484K, Spike V1176F, N A119S, N G204R, N M234I, N R203K, NSP5 L205V, NSP7 L71F, NSP12 P323L |
| EPI_ISL_717953 | 4/11/2020 | 20/12/2020 | Rio de Janeiro / Rio de Janeiro | P.2 | Spike A846S, Spike D614G, Spike E484K, Spike V1176F, N A119S, N G204R, N M234I, N R203K, NSP2 K239R, NSP5 L205V, NSP7 L71F, NSP12 P323L |
| EPI_ISL_717954 | 4/11/2020 | 20/12/2020 | Rio de Janeiro / Rio de Janeiro | P.2 | Spike D614G, Spike E484K, Spike S12F, Spike V1176F, N A119S, N G204R, N M234I, N R203K, NSP3 M829I, NSP5 L205V, NSP7 L71F, NSP12 P323L |
| EPI_ISL_717955 | 5/11/2020 | 20/12/2020 | Rio de Janeiro / Rio de Janeiro | P.2 | Spike D614G, Spike E484K, Spike V1176F, N A119S, N G204R, N M234I, N R203K, NSP3 Q322P, NSP5 L205V, NSP6 L37F, NSP7 L71F, NSP12 P323L, NSP13 S453G |
| EPI_ISL_717956 | 5/11/2020 | 20/12/2020 | Rio de Janeiro / Rio de Janeiro | P.2 | Spike D614G, Spike E484K, Spike V1176F, N A119S, N G204R, N M234I, N R203K, NS3 G44R, NSP5 L205V, NSP6 A46V, NSP7 L71F, NSP12 P323L |
| EPI_ISL_717957 | 5/11/2020 | 20/12/2020 | Rio de Janeiro / Rio de Janeiro | P.2 | Spike D614G, Spike E484K, Spike V1176F, N A119S, N G204R, N G238C, N M234I, N R203K, NSP5 L205V, NSP7 L71F, NSP12 P323L |
| EPI_ISL_792560 | 6/11/2020 | 10/1/2021 | Amazonas / Manaus | P.2 | Spike D614G, Spike E484K, Spike V1176F, N A119S, N G204R, N M234I, N P151S, N R203K, NSP2 G339S, NSP5 L205V, NSP7 L71F, NSP12 P323L |
| EPI_ISL_792634 | 10/11/2020 | 10/1/2021 | Paraiba | P.2 | Spike D614G, Spike E484K, Spike V1176F, N A119S, N G204R, N M234I, N R203K, NS8 I121L, NSP2 S430L, NSP5 L205V, NSP7 L71F, NSP12 P323L |
| EPI_ISL_792635 | 10/11/2020 | 10/1/2021 | Paraiba | P.2 | Spike D614G, Spike E484K, Spike V1176F, N A119S, N G204R, N M234I, N R203K, NS8 I121L, NSP2 S430L, NSP5 L205V, NSP7 L71F, NSP12 P323L |
| EPI_ISL_833163 | 11/19/2020 | 1/17/2021 | Sao Paulo / Ribeirao do Sul | B.1.1.33 | Spike D614G, Spike E484K, N G204R, N I292T, N R203K, NS6 I33T, NS7b E33A, NSP3 A1711V, NSP3 S1285F, NSP6 F36L, NSP12 P323L, NSP15 K12N |
| EPI_ISL_792639 | 21/11/2020 | 10/1/2021 | Alagoas / Maceio | P.2 | Spike D614G, Spike E484K, Spike V1176F, N A119S, N G204R, N M234I, N R203K, NSP5 L205V, NSP7 L71F, NSP12 P323L |
| EPI_ISL_792650 | 23/11/2020 | 10/1/2021 | Parana / Uniao da Vitoria | P.2 | Spike D614G, Spike E484K, Spike V1176F, N A119S, N G204R, N M234I, N R203K, NSP3 G1849C, NSP3 V1834F, NSP5 L205V, NSP7 L71F, NSP12 P323L, NSP12 Q822H |
| EPI_ISL_792651 | 23/11/2020 | 10/1/2021 | Parana / Dois Vizinhos | P.2 | Spike D614G, Spike E484K, Spike V1176F, N A119S, N G204R, N M234I, N R203K, NSP3 A85V, NSP3 N1587S, NSP5 L205V, NSP7 L71F, NSP12 P323L, NSP16 T35I |
| EPI_ISL_779155 | 23/11/2020 | 7/1/2021 | Rio Grande do Sul / Carlos Barbosa | P.2 | Spike D614G, Spike E484K, Spike V1176F, N A119S, N G204R, N M234I, N R203K, NSP5 L205V, NSP7 L71F, NSP12 P323L, NSP14 M501I |
| EPI_ISL_770556 | 23/11/2020 | 6/1/2021 | Rio Grande do Sul / Canoas | P.2 | Spike D614G, Spike E484K, Spike V1176F, N A119S, N G204R, N M234I, N R203K, NSP5 L205V, NSP7 L71F, NSP12 P323L, NSP12 V174I |
| EPI_ISL_770557 | 24/11/2020 | 6/1/2021 | Rio Grande do Sul / Butia | P.2 | Spike D614G, Spike E484K, Spike V1176F, M A2S, N A119S, N G204R, N M234I, N R203K, NSP2 E167D, NSP5 L205V, NSP7 L71F, NSP12 P323L |
| EPI_ISL_770559 | 24/11/2020 | 6/1/2021 | Rio Grande do Sul / Garibaldi | P.2 | Spike D614G, Spike E484K, Spike V1176F, N A119S, N G204R, N M234I, N R203K, NSP4 K86R, NSP5 L205V, NSP7 L71F, NSP12 P323L |
| EPI_ISL_770560 | 24/11/2020 | 6/1/2021 | Rio Grande do Sul / Garibaldi | P.2 | Spike D614G, Spike E484K, Spike V1176F, N A119S, N G204R, N M234I, N R203K, NSP5 L205V, NSP7 L71F, NSP12 P323L |
| EPI_ISL_770561 | 24/11/2020 | 6/1/2021 | Rio Grande do Sul / Garibaldi | P.2 | Spike D614G, Spike E484K, Spike V1176F, N A119S, N G204R, N M234I, N R203K, NSP4 K86R, NSP5 L205V, NSP7 L71F, NSP12 P323L |
| EPI_ISL_770563 | 24/11/2020 | 6/1/2021 | Rio Grande do Sul / Garibaldi | P.2 | Spike D614G, Spike E484K, Spike V1176F, N A119S, N G204R, N M234I, N R203K, NSP3 L561F, NSP5 L205V, NSP7 L71F, NSP12 P323L, NSP15 P205L |
| EPI_ISL_770564 | 24/11/2020 | 6/1/2021 | Rio Grande do Sul / Garibaldi | P.2 | Spike D614G, Spike E484K, Spike V1176F, M S197T, N A119S, N G204R, N M234I, N R203K, NSP4 K86R, NSP5 L205V, NSP7 L71F, NSP12 P323L |
| EPI_ISL_770566 | 24/11/2020 | 6/1/2021 | Rio Grande do Sul / Linha Nova | P.2 | Spike D614G, Spike E484K, Spike V1176F, N A119S, N A211V, N D402Y, N G204R, N M234I, N R203K, NS3 A99S, NSP3 V381A, NSP5 L205V, NSP7 L71F, NSP12 P323L, NSP15 L162F |
| EPI_ISL_770570 | 24/11/2020 | 6/1/2021 | Rio Grande do Sul / Carlos Barbosa | P.2 | Spike D614G, Spike E484K, Spike V1176F, N A119S, N G204R, N M234I, N R203K, NSP5 L205V, NSP7 L71F, NSP12 P323L, NSP14 M501I |
| EPI_ISL_770553 | 24/11/2020 | 6/1/2021 | Rio Grande do Sul / Canoas | P.2 | Spike A626S, Spike D614G, Spike E484K, Spike V1176F, N A119S, N G204R, N M234I, N R203K, NSP5 L205V, NSP7 L71F, NSP12 P323L |
| EPI_ISL_770578 | 24/11/2020 | 6/1/2021 | Rio Grande do Sul / Canoas | P.2 | Spike D614G, Spike D627G, Spike E484K, Spike V1176F, E L19F, N A119S, N G204R, N M234I, N R203K, NS3 L52F, NSP2 E612K, NSP4 S386F, NSP5 L205V, NSP7 L71F, NSP12 P323L |
| EPI_ISL_770581 | 24/11/2020 | 6/1/2021 | Rio Grande do Sul / Canoas | P.2 | Spike D614G, Spike E484K, Spike V1176F, N A119S, N G204R, N M234I, N R203K, NSP5 L205V, NSP7 L71F, NSP12 P323L |
| EPI_ISL_770584 | 24/11/2020 | 6/1/2021 | Rio Grande do Sul / Canoas | P.2 | Spike D614G, Spike E484K, Spike V1176F, N A119S, N G204R, N M234I, N R203K, NSP3 S1375F, NSP5 L205V, NSP7 L71F, NSP12 P323L, NSP13 V386A |
| EPI_ISL_770598 | 24/11/2020 | 6/1/2021 | Rio Grande do Sul / Novo Hamburgo | P.2 | Spike D614G, Spike E484K, Spike V1176F, N A119S, N G204R, N M234I, N R203K, NSP2 K454N, NSP5 L205V, NSP7 L71F, NSP12 P323L |

| | | | | | |
|---|---|---|---|---|---|
| EPI_ISL_792652 | 25/11/2020 | 10/1/2021 | Parana / Cascavel | P.2 | Spike D614G, Spike E484K, Spike V1176F, N A119S, N G204R, N M234I, N R203K, NS3 S165Y, NSP3 V1897I, NSP5 L205V, NSP7 L71F, NSP9 T19I, NSP12 P323L |
| EPI_ISL_770565 | 25/11/2020 | 6/1/2021 | Rio Grande do Sul / Barao | P.2 | Spike D614G, Spike E484K, Spike V1176F, N A119S, N G204R, N M234I, N R203K, NS3 T151I, NS6 W27L, NSP3 E95D, NSP3 L327F, NSP5 L205V, NSP7 L71F, NSP8 P133S, NSP12 D477A, NSP12 P323L |
| EPI_ISL_770568 | 25/11/2020 | 6/1/2021 | Rio Grande do Sul / Linha Nova | P.2 | Spike D614G, Spike E484K, Spike V483F, Spike V1176F, N A119S, N G204R, N M234I, N R203K, NSP3 V1759L, NSP5 L205V, NSP7 L71F, NSP12 P323L, NSP14 M501I |
| EPI_ISL_770571 | 25/11/2020 | 6/1/2021 | Rio Grande do Sul / Carlos Barbosa | P.2 | Spike D614G, Spike E484K, Spike V1176F, N A119S, N G204R, N M234I, N R203K, NSP3 T720A, NSP5 L205V, NSP7 L71F, NSP12 P323L, NSP16 T140I |
| EPI_ISL_770579 | 25/11/2020 | 6/1/2021 | Rio Grande do Sul / Canoas | P.2 | Spike D614G, Spike E484K, Spike I1115V, Spike V1176F, N A119S, N G204R, N M234I, N R203K, NSP2 L271F, NSP5 L205V, NSP7 L71F, NSP12 P323L |
| EPI_ISL_779159 | 25/11/2020 | 7/1/2021 | Rio Grande do Sul / Canoas | P.2 | Spike D614G, Spike E484K, Spike V1176F, N A119S, N G204R, N M234I, N R203K, NSP5 L205V, NSP6 L37F, NSP7 L71F, NSP12 P323L |
| EPI_ISL_770580 | 25/11/2020 | 6/1/2021 | Rio Grande do Sul / Canoas | P.2 | Spike D614G, Spike E484K, Spike L5F, Spike V1176F, N A119S, N G204R, N M234I, N R203K, NS3 T89I, NSP5 L205V, NSP7 L71F, NSP12 P323L, NSP16 A188S |
| EPI_ISL_770583 | 25/11/2020 | 6/1/2021 | Rio Grande do Sul / Canoas | P.2 | Spike D614G, Spike E484K, Spike V1176F, N A119S, N G204R, N G238C, N M234I, N R203K, NS3 D27Y, NSP5 L205V, NSP7 L71F, NSP12 A16V, NSP12 P323L |
| EPI_ISL_770587 | 25/11/2020 | 6/1/2021 | Rio Grande do Sul / Butia | P.2 | Spike D614G, Spike E484K, Spike V1176F, N A119S, N D341Y, N G204R, N M234I, N R203K, NS3 G172C, NSP5 L205V, NSP7 L71F, NSP12 P323L, NSP14 H95Y |
| EPI_ISL_770589 | 25/11/2020 | 6/1/2021 | Rio Grande do Sul / Garibaldi | P.2 | Spike D614G, Spike E484K, Spike V1176F, N A119S, N G204R, N M234I, N R203K, NSP3 K1679N, NSP3 S692F, NSP3 V1897I, NSP5 L205V, NSP7 L71F, NSP9 T19I, NSP12 P323L |
| EPI_ISL_770591 | 25/11/2020 | 6/1/2021 | Rio Grande do Sul / Garibaldi | P.2 | Spike D614G, Spike E484K, Spike V1176F, N A119S, N G204R, N M234I, N R203K, NSP4 K86R, NSP5 L205V, NSP7 L71F, NSP12 P323L |
| EPI_ISL_770592 | 25/11/2020 | 6/1/2021 | Rio Grande do Sul / Garibaldi | P.2 | Spike D614G, Spike E484K, Spike V1176F, N A119S, N G204R, N M234I, N R203K, NSP5 L205V, NSP7 L71F, NSP12 P169A, NSP12 P323L |
| EPI_ISL_770593 | 25/11/2020 | 6/1/2021 | Rio Grande do Sul / Garibaldi | P.2 | Spike D614G, Spike E484K, Spike V1176F, N A119S, N G204R, N M234I, N R203K, NSP5 L205V, NSP7 L71F, NSP12 P323L |
| EPI_ISL_770594 | 25/11/2020 | 6/1/2021 | Rio Grande do Sul / Garibaldi | P.2 | Spike D614G, Spike E484K, Spike V1176F, N A119S, N G204R, N M234I, N R203K, NSP5 L205V, NSP7 L71F, NSP12 P169A, NSP12 P323L |
| EPI_ISL_770595 | 25/11/2020 | 6/1/2021 | Rio Grande do Sul / Garibaldi | P.2 | Spike D614G, Spike E484K, Spike V1176F, N A119S, N G204R, N M234I, N R203K, NSP4 K86R, NSP5 L205V, NSP7 L71F, NSP12 P323L |
| EPI_ISL_770596 | 25/11/2020 | 6/1/2021 | Rio Grande do Sul / Garibaldi | P.2 | Spike D614G, Spike E484K, Spike V1176F, N A119S, N G204R, N M234I, N R203K, NSP5 L205V, NSP7 L71F, NSP12 P169A, NSP12 P323L |
| EPI_ISL_770552 | 25/11/2020 | 6/1/2021 | Rio Grande do Sul / Morro Reuter | P.2 | Spike D614G, Spike E484K, Spike V1176F, N A119S, N D401Y, N G204R, N M234I, N N48del, N R203K, NSP3 T754I, NSP5 L205V, NSP7 L71F, NSP12 P323L |
| EPI_ISL_770602 | 26/11/2020 | 6/1/2021 | Rio Grande do Sul / Garibaldi | P.2 | Spike D614G, Spike E484K, Spike V1176F, N A119S, N G204R, N M234I, N R203K, NSP4 K86R, NSP5 L205V, NSP7 L71F, NSP12 P323L |
| EPI_ISL_770603 | 26/11/2020 | 6/1/2021 | Rio Grande do Sul / Garibaldi | P.2 | Spike D614G, Spike E484K, Spike V1176F, N A119S, N G204R, N M234I, N R203K, NSP5 L205V, NSP6 L37F, NSP7 L71F, NSP12 P323L |
| EPI_ISL_770604 | 26/11/2020 | 6/1/2021 | Rio Grande do Sul / Garibaldi | P.2 | Spike D614G, Spike E484K, Spike V1176F, N A119S, N G204R, N M234I, N R203K, NSP5 L205V, NSP7 L71F, NSP12 P323L, NSP14 M501I |
| EPI_ISL_770605 | 26/11/2020 | 6/1/2021 | Rio Grande do Sul / Garibaldi | P.2 | Spike D614G, Spike E484K, Spike V1176F, N A119S, N G204R, N M234I, N R203K, NSP5 L205V, NSP7 L71F, NSP12 P169A, NSP12 P323L |
| EPI_ISL_770606 | 26/11/2020 | 6/1/2021 | Rio Grande do Sul / Garibaldi | P.2 | Spike D614G, Spike E484K, Spike V1176F, N A119S, N G204R, N M234I, N R203K, NSP5 L205V, NSP7 L71F, NSP12 P323L, NSP14 M501I |
| EPI_ISL_770607 | 26/11/2020 | 6/1/2021 | Rio Grande do Sul / Garibaldi | P.2 | Spike D614G, Spike E484K, Spike V1176F, N A119S, N G204R, N M234I, N R203K, NSP5 L205V, NSP7 L71F, NSP12 P169A, NSP12 P323L |
| EPI_ISL_770617 | 26/11/2020 | 6/1/2021 | Rio Grande do Sul / Carlos Barbosa | P.2 | Spike D614G, Spike E484K, Spike P1263S, Spike V1176F, N A119S, N G204R, N M234I, N R203K, NS7b D36G, NSP5 L205V, NSP7 L71F, NSP12 P323L, NSP15 P205L |
| EPI_ISL_833161 | 11/26/2020 | 1/17/2021 | Sao Paulo / Sao Paulo | P.2 | Spike A879S, Spike D614G, Spike E484K, Spike V1176F, N A119S, N G204R, N M234I, N R203K, NSP5 L205V, NSP7 L71F, NSP12 P323L |
| EPI_ISL_770616 | 27/11/2020 | 6/1/2021 | Rio Grande do Sul / Carlos Barbosa | P.2 | Spike D614G, Spike E484K, Spike V1176F, N A119S, N G204R, N M234I, N R203K, NSP5 L205V, NSP7 L71F, NSP12 P323L, NSP14 M501I |
| EPI_ISL_770618 | 27/11/2020 | 6/1/2021 | Rio Grande do Sul / Butia | P.2 | Spike D614G, Spike E484K, Spike V1176F, N A119S, N D377Y, N G204R, N M234I, N R203K, NSP5 L205V, NSP5 P96L, NSP7 L71F, NSP12 P323L, NSP15 V9F |
| EPI_ISL_770619 | 27/11/2020 | 6/1/2021 | Rio Grande do Sul / Butia | P.2 | Spike D614G, Spike E484K, Spike V1176F, M A2S, N A119S, N G204R, N M234I, N R203K, NSP2 E167D, NSP5 L205V, NSP7 L71F, NSP12 P323L |
| EPI_ISL_770620 | 27/11/2020 | 6/1/2021 | Rio Grande do Sul / Garibaldi | P.2 | Spike D614G, Spike E484K, Spike V1176F, N A119S, N G204R, N M234I, N R203K, NSP4 K86R, NSP5 L205V, NSP7 L71F, NSP12 P323L |
| EPI_ISL_770554 | 27/11/2020 | 6/1/2021 | Rio Grande do Sul / Garibaldi | P.2 | Spike A626S, Spike D614G, Spike E484K, Spike V1176F, N A119S, N G204R, N M234I, N R203K, NSP5 L205V, NSP7 L71F, NSP12 P323L |
| EPI_ISL_770621 | 27/11/2020 | 6/1/2021 | Rio Grande do Sul / Garibaldi | P.2 | Spike D614G, Spike E484K, Spike V1176F, N A119S, N G204R, N M234I, N R203K, NSP4 K86R, NSP5 L205V, NSP7 L71F, NSP12 P323L |
| EPI_ISL_770622 | 27/11/2020 | 6/1/2021 | Rio Grande do Sul / Garibaldi | P.2 | Spike D614G, Spike E484K, Spike V1176F, N A119S, N G204R, N M234I, N R203K, NSP5 L205V, NSP7 L71F, NSP12 P169A, NSP12 P323L |
| EPI_ISL_770624 | 27/11/2020 | 6/1/2021 | Rio Grande do Sul / Garibaldi | P.2 | Spike D614G, Spike E484K, Spike V1176F, N A119S, N G204R, N M234I, N R203K, NSP5 L205V, NSP7 L71F, NSP12 P169A, NSP12 P323L |

| | | | | | |
|---|---|---|---|---|---|
| EPI_ISL_770625 | 27/11/2020 | 6/1/2021 | Rio Grande do Sul / Garibaldi | P.2 | Spike D614G, Spike E484K, Spike T572I, Spike V1176F, N A119S, N G204R, N M234I, N R203K, NSP4 K86R, NSP5 L205V, NSP7 L71F, NSP12 P323L |
| EPI_ISL_770628 | 28/11/2020 | 6/1/2021 | Rio Grande do Sul / Garibaldi | P.2 | Spike D614G, Spike E484K, Spike V1176F, N A119S, N G204R, N M234I, N R203K, NSP4 K86R, NSP5 L205V, NSP7 L71F, NSP12 P323L |
| EPI_ISL_833158 | 11/30/2020 | 1/17/2021 | Sao Paulo / Sao Bernardo do Campo | P.2 | Spike D614G, Spike E484K, Spike V1176F, N A119S, N G204R, N G238C, N M234I, N R203K, NSP3 D9G, NSP3 L728I, NSP3 S315R, NSP3 S1406F, NSP5 L205V, NSP7 L71F, NSP12 P323L, NSP14 M62I |
| EPI_ISL_792642 | 4/12/2020 | 10/1/2021 | Alagoas / Maceio | P.2 | Spike D614G, Spike E484K, Spike V1176F, N A119S, N G204R, N M234I, N R203K, NSP5 L205V, NSP7 L71F, NSP12 P323L |
| EPI_ISL_833137 | 12/4/2020 | 1/17/2021 | Amazonas / Manaus | P.1 | Spike D138Y, Spike D614G, Spike E484K, Spike H655Y, Spike K417T, Spike L18F, Spike N501Y, Spike P26S, Spike R190S, Spike T20N, Spike T1027I, Spike V1176F, N G204R, N P80R, N R203K, NS3 S253P, NS8 E92K, NSP3 K977Q, NSP3 S370L, NSP6 F108del, NSP6 G107del, NSP6 S106del, NSP12 P323L, NSP13 E341D |
| EPI_ISL_755642 | 5/12/2020 | 1/1/2021 | São Paulo / Mogi das Cruzes | P.2 | Spike A626S, Spike A892V, Spike D614G, Spike E484K, Spike V1176F, N A119S, N G204R, N G238C, N M234I, N P279L, N R203K, NSP3 R748S, NSP5 L205V, NSP7 L71F, NSP9 T34I, NSP12 P323L |
| EPI_ISL_755649 | 6/12/2020 | 1/1/2021 | São Paulo / Sao Bernardo do Campo | B.1.1.33 | Spike D614G, Spike E484K, N G204R, N I292T, N R203K, NS3 V202L, NS6 I33T, NS7b E33A, NSP2 I296V, NSP3 A1711V, NSP3 S1285F, NSP6 F36L, NSP12 P323L, NSP15 K12N |
| EPI_ISL_836143 | 12/7/2020 | 1/18/2021 | Sao Paulo / Atibaia | P.2 | Spike D614G, Spike E484K, Spike V1176F, N A119S, N G204R, N M234I, N R203K, NSP2 Q427R, NSP5 L205V, NSP7 L71F, NSP12 P323L |
| EPI_ISL_755645 | 8/12/2020 | 1/1/2021 | São Paulo / Taboao da Serra | P.2 | Spike D614G, Spike E484K, Spike V1176F, N A119S, N G204R, N M234I, N R203K, NS8 I121L, NSP3 G307C, NSP3 P654S, NSP5 L205V, NSP7 L71F, NSP12 P323L |
| EPI_ISL_836977 | 12/8/2020 | 1/18/2021 | Sao Paulo / Sao Jose dos Campos | P.2 | Spike D614G, Spike E484K, Spike V1176F, N A119S, N G204R, N M234I, N R203K, NSP3 G1451C, NSP5 L205V, NSP7 L71F, NSP12 P323L |
| EPI_ISL_755652 | 9/12/2020 | 1/1/2021 | São Paulo / Embu das Artes | B.1.1.33 | Spike D614G, Spike E484K, M V23L, N G204R, N I292T, N R203K, NS6 I33T, NS7b E33A, NSP3 A1711V, NSP3 S1285F, NSP6 F36L, NSP7 L71F, NSP12 P323L, NSP15 K12N, NSP16 G113C |
| EPI_ISL_755651 | 10/12/2020 | 1/1/2021 | São Paulo / Poa | P.2 | Spike D614G, Spike E484K, Spike L5F, Spike V1176F, N A119S, N G204R, N M234I, N R203K, NSP5 L205V, NSP7 L71F, NSP12 P323L |
| EPI_ISL_755653 | 12/12/2020 | 1/1/2021 | São Paulo / Guararema | P.2 | Spike D614G, Spike E484K, Spike V1176F, M H125Y, N A119S, N G204R, N G238C, N M234I, N R203K, NS7a Q21H, NSP3 T819I, NSP5 L205V, NSP7 L71F, NSP12 P323L |
| EPI_ISL_804827 | 15/12/2020 | 12/1/2021 | Amazonas | P.2 | Spike A688V, Spike D614G, Spike E484K, Spike V1176F, N A119S, N G204R, N M234I, N R203K, NSP3 T183I, NSP5 L205V, NSP7 L71F, NSP12 P323L, NSP13 P53L |
| EPI_ISL_804819 | 16/12/2020 | 12/1/2021 | Amazonas | P.1 | Spike D614G, Spike E484K, Spike H655Y, Spike K417T, Spike N501Y, Spike T1027I, Spike V1176F, N G204R, N P80R, N R203K, NS3 S253P, NS8 E92K, NSP3 K977Q, NSP3 S370L, NSP6 F108del, NSP6 G107del, NSP6 S106del, NSP12 P323L, NSP13 E341D |
| EPI_ISL_833167 | 12/16/2020 | 1/17/2021 | Amazonas | P.1 | Spike D138Y, Spike D614G, Spike E484K, Spike H655Y, Spike K417T, Spike L18F, Spike N501Y, Spike P26S, Spike R190S, Spike T20N, Spike T1027I, Spike V1176F, N G204R, N P80R, N R203K, N T16M, NS3 S253P, NS8 E92K, NSP3 K977Q, NSP3 S370L, NSP6 F108del, NSP6 G107del, NSP6 S106del, NSP12 P323L, NSP13 E341D |
| EPI_ISL_804824 | 17/12/2020 | 12/1/2021 | Amazonas | P.1 | Spike D614G, Spike E484K, Spike K417T, Spike N501Y, Spike T1027I, Spike V1176F, N G204R, N P80R, N R203K, NS3 S253P, NS8 E92K, NSP3 K977Q, NSP3 S370L, NSP6 F108del, NSP6 G107del, NSP6 S106del, NSP12 P323L, NSP13 E341D |
| EPI_ISL_804823 | 17/12/2020 | 12/1/2021 | Amazonas | P.1 | Spike D138Y, Spike D614G, Spike E484K, Spike H655Y, Spike K417T, Spike L18F, Spike N501Y, Spike P26S, Spike R190S, Spike T20N, Spike T1027I, Spike V1176F, N G204R, N P80R, N R203K, NS3 S253P, NS8 E92K, NSP3 K977Q, NSP3 S370L, NSP6 F108del, NSP6 G107del, NSP6 S106del, NSP12 P323L, NSP13 E341D |
| EPI_ISL_804828 | 17/12/2020 | 12/1/2021 | Amazonas | B.1.1.33 | Spike A344S, Spike D614G, Spike E484K, N G204R, N I292T, N R203K, NS6 I33T, NS6 I60T, NS7b E33A, NSP1 T170I, NSP2 P13L, NSP3 A54T, NSP3 A1711V, NSP6 F36L, NSP12 P323L |
| EPI_ISL_833171 | 12/17/2020 | 1/17/2021 | Amazonas | P.1 | Spike D138Y, Spike D614G, Spike E484K, Spike H655Y, Spike K417T, Spike L18F, Spike N501Y, Spike P26S, Spike R190S, Spike T20N, Spike T1027I, Spike V1176F, N G204R, N P80R, N R203K, NS3 S253P, NS8 E92K, NSP3 K977Q, NSP3 S370L, NSP6 F108del, NSP6 G107del, NSP6 S106del, NSP12 P323L, NSP13 E341D |
| EPI_ISL_833172 | 12/17/2020 | 1/17/2021 | Amazonas | P.1 | Spike D138Y, Spike D614G, Spike E484K, Spike H655Y, Spike K417T, Spike L18F, Spike N501Y, Spike P26S, Spike R190S, Spike T20N, Spike T1027I, Spike V1176F, N G204R, N P80R, N R203K, NS3 S253P, NS8 E92K, NSP3 K977Q, NSP3 S370L, NSP6 F108del, NSP6 G107del, NSP6 S106del, NSP12 P323L, NSP13 E341D |
| EPI_ISL_833176 | 12/17/2020 | 1/17/2021 | Amazonas | P.2 | Spike D614G, Spike E484K, Spike V1176F, N A119S, N G204R, N M234I, N R203K, NSP5 L205V, NSP7 L71F, NSP12 P323L |
| EPI_ISL_804829 | 18/12/2020 | 12/1/2021 | Amazonas | P.2 | Spike D614G, Spike E484K, Spike V1176F, N A119S, N G204R, N M234I, N R203K, NS3 T151I, NS8 I39M, NSP3 E95D, NSP3 L327F, NSP5 L205V, NSP7 L71F, NSP12 D477A, NSP12 P323L |

| | | | | | |
|---|---|---|---|---|---|
| EPI_ISL_833173 | 12/18/2020 | 1/17/2021 | Amazonas | P.1 | Spike D138Y, Spike D614G, Spike E484K, Spike H655Y, Spike K417T, Spike L18F, Spike N501Y, Spike P26S, Spike R190S, Spike T20N, Spike T1027I, Spike V1176F, N G204R, N P80R, N R203K, NS3 S253P, NS8 E92K, NSP3 K977Q, NSP3 S370L, NSP6 F108del, NSP6 G107del, NSP6 S106del, NSP12 P323L, NSP13 E341D |
| EPI_ISL_804821 | 21/12/2020 | 12/1/2021 | Amazonas | P.1 | Spike D614G, Spike E484K, Spike H655Y, Spike K417T, Spike N501Y, Spike T1027I, Spike V1176F, N G204R, N P80R, N R203K, NS3 S253P, NS8 E92K, NSP3 K977Q, NSP3 S370L, NSP6 F108del, NSP6 G107del, NSP6 S106del, NSP12 P323L, NSP13 E341D |
| EPI_ISL_804814 | 21/12/2020 | 12/1/2021 | Amazonas | P.1 | Spike D614G, Spike E484K, Spike H655Y, Spike K417T, Spike N501Y, Spike T1027I, Spike V1176F, N G204R, N P80R, N R203K, NS3 S253P, NS8 E92K, NSP3 K977Q, NSP6 F108del, NSP6 G107del, NSP6 S106del, NSP12 P323L, NSP13 E341D |
| EPI_ISL_804820 | 21/12/2020 | 12/1/2021 | Amazonas | P.1 | Spike D614G, Spike E484K, Spike H655Y, Spike K417T, Spike N501Y, Spike T1027I, Spike V1176F, N G204R, N P80R, N R203K, NS3 S253P, NS8 E92K, NSP3 K977Q, NSP3 S370L, NSP3 S1735F, NSP6 F108del, NSP6 G107del, NSP6 S106del, NSP12 P323L, NSP13 E341D |
| EPI_ISL_833138 | 12/21/2020 | 1/17/2021 | Amazonas / Manaus | P.1 | Spike D138Y, Spike D614G, Spike E484K, Spike H655Y, Spike K417T, Spike L18F, Spike N501Y, Spike P26S, Spike R190S, Spike T20N, Spike T1027I, Spike V1176F, N G204R, N P80R, N R203K, NS3 S253P, NS8 E92K, NSP3 K977Q, NSP3 S370L, NSP6 F108del, NSP6 G107del, NSP6 S106del, NSP12 P323L, NSP13 E341D |
| EPI_ISL_804816 | 22/12/2020 | 12/1/2021 | Amazonas | P.2 | Spike A688V, Spike D614G, Spike E484K, Spike V1176F, N A119S, N G204R, N M234I, N R203K, NSP5 L205V, NSP7 L71F, NSP12 P323L, NSP13 P53L |
| EPI_ISL_833139 | 12/22/2020 | 1/17/2021 | Amazonas / Manaus | P.1 | Spike D138Y, Spike D614G, Spike E484K, Spike H655Y, Spike K417T, Spike L18F, Spike N501Y, Spike P26S, Spike R190S, Spike T20N, Spike T1027I, Spike V1176F, N G204R, N P80R, N R203K, NS3 S253P, NS8 E92K, NSP3 K977Q, NSP3 S370L, NSP6 F108del, NSP6 G107del, NSP6 S106del, NSP12 P323L, NSP13 E341D, NSP14 P443S |
| EPI_ISL_833174 | 12/22/2020 | 1/17/2021 | Amazonas | P.1 | Spike D138Y, Spike D614G, Spike E484K, Spike H655Y, Spike K417T, Spike L18F, Spike N501Y, Spike P26S, Spike R190S, Spike T20N, Spike T1027I, Spike V1176F, N G204R, N P80R, N R203K, NS3 S253P, NS8 E92K, NSP3 K977Q, NSP3 S370L, NSP6 F108del, NSP6 G107del, NSP6 S106del, NSP12 P323L, NSP13 E341D |
| EPI_ISL_833175 | 12/22/2020 | 1/17/2021 | Amazonas | P.2 | Spike A688V, Spike D614G, Spike E484K, Spike V1176F, N A119S, N G204R, N M234I, N R203K, NSP3 T183I, NSP5 L205V, NSP7 L71F, NSP12 P323L, NSP13 P53L |
| EPI_ISL_833140 | 12/23/2020 | 1/17/2021 | Amazonas / Manaus | P.1 | Spike A27V, Spike D138Y, Spike D614G, Spike E484K, Spike H655Y, Spike K417T, Spike L18F, Spike N501Y, Spike P26S, Spike R190S, Spike T20N, Spike T1027I, Spike V1176F, N G204R, N P80R, N R203K, NS3 S253P, NS8 E92K, NSP3 K977Q, NSP3 S370L, NSP6 F108del, NSP6 G107del, NSP6 S106del, NSP12 P323L, NSP13 E341D |
| EPI_ISL_833169 | 12/23/2020 | 1/17/2021 | Amazonas | P.1 | Spike D138Y, Spike D614G, Spike E484K, Spike H655Y, Spike K417T, Spike L18F, Spike N501Y, Spike P26S, Spike R190S, Spike T20N, Spike T1027I, Spike V1176F, N G204R, N P80R, N R203K, NS3 S253P, NS8 E92K, NSP3 K977Q, NSP3 S370L, NSP6 F108del, NSP6 G107del, NSP6 S106del, NSP12 P323L, NSP13 E341D |
| EPI_ISL_833170 | 12/23/2020 | 1/17/2021 | Amazonas | P.1 | Spike D138Y, Spike D614G, Spike E484K, Spike H655Y, Spike K417T, Spike L18F, Spike N501Y, Spike P26S, Spike R190S, Spike T20N, Spike T1027I, Spike V1176F, N A119S, N G204R, N P80R, N R203K, NS3 S253P, NS8 E92K, NSP3 K977Q, NSP3 S370L, NSP6 F108del, NSP6 G107del, NSP6 S106del, NSP12 P323L, NSP13 E341D |
| EPI_ISL_833136 | 12/29/2020 | 1/17/2021 | Amazonas / Manaus | P.1 | Spike D138Y, Spike D614G, Spike E484K, Spike H655Y, Spike K417T, Spike L18F, Spike N501Y, Spike P26S, Spike R190S, Spike T20N, Spike T1027I, Spike V1176F, N A267V, N G204R, N P80R, N P151L, N R203K, NS3 S253P, NS8 E92K, NSP3 K977Q, NSP3 S370L, NSP6 F108del, NSP6 G107del, NSP6 S106del, NSP12 P323L, NSP13 E341D |
| EPI_ISL_811149 | 12/30/2020 | 1/13/2021 | Amazonas / Manaus | B.1.1.28 | Spike D138Y, Spike D614G, Spike E484K, Spike H655Y, Spike K417T, Spike L18F, Spike N501Y, Spike P26S, Spike R190S, Spike T20N, Spike T1027I, Spike V1176F, N G204R, N P80R, N R203K, NS3 S253P, NS8 E92K, NSP3 K977Q, NSP3 S370L, NSP6 F108del, NSP6 G107del, NSP6 S106del, NSP12 P323L, NSP13 E341D, NSP13 T216N |

We gratefully acknowledge the following Authors from the Originating laboratories responsible for obtaining the specimens, as well as the Submitting laboratories where the genome data were generated and shared via GISAID, on which this research is based.

All Submitters of data may be contacted directly via www.gisaid.org

Authors are sorted alphabetically.

| Accession ID | Originating Laboratory | Submitting Laboratory | Authors |
|---|---|---|---|
| EPI_ISL_717921, EPI_ISL_717922, EPI_ISL_717924, EPI_ISL_717925, EPI_ISL_717926, EPI_ISL_717927, EPI_ISL_717928, EPI_ISL_717929, EPI_ISL_717930, EPI_ISL_717931, EPI_ISL_717932, EPI_ISL_717933, EPI_ISL_717934, EPI_ISL_717935, EPI_ISL_717936, EPI_ISL_717937, EPI_ISL_717938, EPI_ISL_717939, EPI_ISL_717940, EPI_ISL_717941, EPI_ISL_717942, EPI_ISL_717943, EPI_ISL_717944, EPI_ISL_717945, EPI_ISL_717946, EPI_ISL_717947, EPI_ISL_717948, EPI_ISL_717949, EPI_ISL_717950, EPI_ISL_717951, EPI_ISL_717952, EPI_ISL_717953, EPI_ISL_717954, EPI_ISL_717955, EPI_ISL_717956, EPI_ISL_717957 | | | |
| see above | Laboratorio de Virologia Molecular / UFRJ | Bioinformatics Laboratory / LNCC | Carolina M Voloch, Ronaldo da Silva F Jr, Luiz G P de Almeida, Cynthia C Cardoso, Otavio Bustrolini, Alexandra L Gerber, Ana Paula de C Guimarães, Diana Mariani, Andréa Cony Cavalcanti, Claudia dos Santos Rodrigues, Terezinha M P P Castiñeira, Amilcar Tanuri, Ana Tereza R de Vasconcelos |
| EPI_ISL_755642 | Instituto Adolfo Lutz - Central | Instituto Adolfo Lutz, Interdiciplinary Procedures Center, Strategic Laboratory | Claudio Tavares Sacchi, Claudia Regina Gonçalves, Erica Valessa Ramos Gomes, Karoline Rodrigues Campos |
| EPI_ISL_755645 | Lab LOC - Itapecerica da Serra | Instituto Adolfo Lutz, Interdiciplinary Procedures Center, Strategic Laboratory | Claudio Tavares Sacchi, Claudia Regina Gonçalves, Erica Valessa Ramos Gomes, Karoline Rodrigues Campos |
| EPI_ISL_755649 | Instituto Adolfo Lutz - Regional de Santo Andre | Instituto Adolfo Lutz, Interdiciplinary Procedures Center, Strategic Laboratory | Claudio Tavares Sacchi, Claudia Regina Gonçalves, Erica Valessa Ramos Gomes, Karoline Rodrigues Campos |
| EPI_ISL_755651 | Instituto Adolfo Lutz - Central | Instituto Adolfo Lutz, Interdiciplinary Procedures Center, Strategic Laboratory | Claudio Tavares Sacchi, Claudia Regina Gonçalves, Erica Valessa Ramos Gomes, Karoline Rodrigues Campos |
| EPI_ISL_755652 | Lab LOC - Itapecerica da Serra | Instituto Adolfo Lutz, Interdiciplinary Procedures Center, Strategic Laboratory | Claudio Tavares Sacchi, Claudia Regina Gonçalves, Erica Valessa Ramos Gomes, Karoline Rodrigues Campos |
| EPI_ISL_755653 | Instituto Adolfo Lutz - Central | Instituto Adolfo Lutz, Interdiciplinary Procedures Center, Strategic Laboratory | Claudio Tavares Sacchi, Claudia Regina Gonçalves, Erica Valessa Ramos Gomes, Karoline Rodrigues Campos |
| EPI_ISL_756294 | Center for Biotechnology and Cell Therapy, São Rafael Hospital, Salvador, Brazil | Center for Biotechnology and Cell Therapy, São Rafael Hospital, Salvador, Brazil | Carolina Kymie Vasques Nonaka, Marília Miranda Franco, Tiago Gräf, Ana Verena Almeida Mendes, Renato Santana de Aguiar, Marta Giovanetti, Bruno Solano de Freitas Souza |
| EPI_ISL_770552, EPI_ISL_770553, EPI_ISL_770554, EPI_ISL_770556, EPI_ISL_770557, EPI_ISL_770559, EPI_ISL_770560, EPI_ISL_770561, EPI_ISL_770563, EPI_ISL_770564, EPI_ISL_770565, EPI_ISL_770566, EPI_ISL_770568, EPI_ISL_770570, EPI_ISL_770571, EPI_ISL_770578, EPI_ISL_770579, EPI_ISL_770580, EPI_ISL_770581, EPI_ISL_770583, EPI_ISL_770584, EPI_ISL_770587, EPI_ISL_770589, EPI_ISL_770591, EPI_ISL_770592, EPI_ISL_770593, EPI_ISL_770594, EPI_ISL_770595, EPI_ISL_770596, EPI_ISL_770598, EPI_ISL_770602, EPI_ISL_770603, EPI_ISL_770604, EPI_ISL_770605, EPI_ISL_770606, EPI_ISL_770607, EPI_ISL_770616, EPI_ISL_770617, EPI_ISL_770618, EPI_ISL_770619, EPI_ISL_770620, EPI_ISL_770621, EPI_ISL_770622, EPI_ISL_770624, EPI_ISL_770625, EPI_ISL_770628, EPI_ISL_779155, EPI_ISL_779159 | | | |
| see above | Laboratório de Microbiologia Molecular - Universidade FEEVALE | Bioinformatics Laboratory / LNCC | Felipe Benites, Fernando Rosado Spilki, Alana Witt Hansen, Juliane Deise Fleck, Juliana Schons, Meriane Demoliner, Ana Karolina Eisen Antunes, Fagner Henrique Heldt, Larissa Mallmann, Bruna Hermann, Ana Luiza Ziulkoski, Vyctoria Goes, Karoline Schallenberger, Matheus Nunes Weber, Paula Rodrigues de Almeida, Alessandra Pavan Lamarca da Silva, Ronaldo da Silva F Jr , Luiz G P de Almeida, Alexandra L Gerber , Ana Paula de C Guimarães,Ana Tereza R de Vasconcelos |
| EPI_ISL_792560 | Laboratorio de Ecologia de Doencas Transmissiveis na Amazonia, Instituto Leonidas e Maria Deane - Fiocruz Amazonia | Laboratorio de Ecologia de Doencas Transmissiveis na Amazonia, Instituto Leonidas e Maria Deane - Fiocruz Amazonia | Valdinete Nascimento, Victor Souza, André Corado, Fernanda Nascimento, George Silva, Ágatha Costa, Karina Pessoa, Debora Duarte, Luciana Gonçalves, Maria Júlia Brandão, Michele Jesus, Felipe Naveca |
| EPI_ISL_792562, EPI_ISL_792634, EPI_ISL_792635 | LACEN-PB | Laboratory of Respiratory Viruses and Measles, Oswaldo Cruz Institute, FIOCRUZ | Paola Resende, Luciana Appolinario, Fernando Motta, Anna Carolina Paixao, Ana Carolina Mendonca, João Felipe Bezerra, Romero Henrique Teixeira de Vasconcelos, Dalane Loudal Florentino Teixeira, Thiago Franco de Oliveira Carneiro, Marilda Siqueira |
| EPI_ISL_792639, EPI_ISL_792642 | LACEN-AL | Laboratory of Respiratory Viruses and Measles, Oswaldo Cruz Institute, FIOCRUZ | Paola Resende, Luciana Appolinario, Fernando Motta, Anna Carolina Paixao, Ana Carolina Mendonca, Anderson Brandao Leite, Marilda Siqueira |
| EPI_ISL_792645, EPI_ISL_792646, EPI_ISL_792650, EPI_ISL_792651, EPI_ISL_792652 | LACEN-PR | Laboratory of Respiratory Viruses and Measles, Oswaldo Cruz Institute, FIOCRUZ | Paola Resende, Luciana Appolinario, Fernando Motta, Anna Carolina Paixao, Ana Carolina Mendonca, Maria do Carmo Debur, Irina Nastassja Riediger, Marilda Siqueira |
| EPI_ISL_804814, EPI_ISL_804816, EPI_ISL_804819, EPI_ISL_804820, EPI_ISL_804821, EPI_ISL_804823, EPI_ISL_804824, EPI_ISL_804827, EPI_ISL_804828, EPI_ISL_804829 | DB Diagnosticos do Brasil | Laboratório de Parasitologia Médica - Instituto de Medicina Tropical - Universidade de São Paulo | Nuno Faria, Ingra Morales Claro, Darlan Candido, Lucas A. Moyses Franco, Pamela dos Santos Andrade, Thais de Moura Coletti, Camila A. Maia da Silva, Flavia Cristina Sales, Erika Regina Manuli, Renato A. Santana, Nelson Gaburo, Cecília da Cunha Camilo, Nelson Abrahim Fraiji, Myuki Alfaia Esashika Crispim, Maria do Perpétuo Socorro Sampaio Carvalho, Andrew Rambaut, Nick Loman, Oliver G. Pybus, Ester C. Sabino; DB; HEMOAM; CDL; CADDE Genomic Network. |
| EPI_ISL_811149 | Laboratorio de Ecologia de Doencas Transmissiveis na Amazonia, Instituto Leonidas e Maria Deane - Fiocruz Amazonia | Laboratorio de Ecologia de Doencas Transmissiveis na Amazonia, Instituto Leonidas e Maria Deane - Fiocruz Amazonia | Valdinete Nascimento, Victor Souza, André Corado, Fernanda Nascimento, George Silva, Ágatha Costa, Debora Duarte, Luciana Gonçalves, Matilde Mejía, Karina Pessoa, Maria Júlia Brandão, Michele Jesus, Felipe Naveca |
| EPI_ISL_832010, EPI_ISL_832013 | Laboratório de Microbiologia Molecular - Universidade FEEVALE | Universidade Federal de Ciências da Saúde de Porto Alegre | Vinícius Bonetti Franceschi, Amanda de Menezes Mayer, Gabriel Dickin Caldana, Carla Andretta Moreira Neves, Patrícia Aline Gröhs Ferrareze, Gabriela Bettella Cybis, Ricardo Ariel Zimerman, Lívia Kmetzsch, Fernando Rosado Spilki, Claudia Elizabeth Thompson |
| EPI_ISL_833136, EPI_ISL_833137, EPI_ISL_833138, EPI_ISL_833139, EPI_ISL_833140 | Laboratorio de Ecologia de Doencas Transmissiveis na Amazonia, Instituto Leonidas e Maria Deane - Fiocruz Amazonia | Laboratorio de Ecologia de Doencas Transmissiveis na Amazonia, Instituto Leonidas e Maria Deane - Fiocruz Amazonia | Valdinete Nascimento, Victor Souza, André Corado, Fernanda Nascimento, George Silva, Ágatha Costa, Debora Duarte, Karina Pessoa, Matilde Mejía, Luciana Gonçalves, Maria Júlia Brandão, Michele Jesus, Felipe Naveca |
| EPI_ISL_833158 | Instituto Adolfo Lutz - Regional de Santo Andre | Instituto Adolfo Lutz, Interdiciplinary Procedures Center, Strategic Laboratory | Claudio Tavares Sacchi, Claudia Regina Gonçalves, Erica Valessa Ramos Gomes, Karoline Rodrigues Campos |
| EPI_ISL_833161 | Instituto Adolfo Lutz - Central | Instituto Adolfo Lutz, Interdiciplinary Procedures Center, Strategic Laboratory | Claudio Tavares Sacchi, Claudia Regina Gonçalves, Erica Valessa Ramos Gomes, Karoline Rodrigues Campos |
| EPI_ISL_833163 | Instituto Adolfo Lutz - Regional de Marilia | Instituto Adolfo Lutz, Interdiciplinary Procedures Center, Strategic Laboratory | Claudio Tavares Sacchi, Claudia Regina Gonçalves, Erica Valessa Ramos Gomes, Karoline Rodrigues Campos |
| EPI_ISL_833167, EPI_ISL_833169, EPI_ISL_833170, EPI_ISL_833171, EPI_ISL_833172, EPI_ISL_833173, EPI_ISL_833174, EPI_ISL_833175, EPI_ISL_833176 | DB Diagnosticos do Brasil | Instituto Adolfo Lutz, Interdiciplinary Procedures Center, Strategic Laboratory | Claudio Tavares Sacchi, Claudia Regina Gonçalves, Erica Valessa Ramos Gomes, Karoline Rodrigues Campos |
| EPI_ISL_836143 | Hospital de Campanha COVID-19 de Mairipora | Instituto Adolfo Lutz, Interdiciplinary Procedures Center, Strategic Laboratory | Claudio Tavares Sacchi, Claudia Regina Gonçalves, Erica Valessa Ramos Gomes, Karoline Rodrigues Campos |
| EPI_ISL_836977 | Hospital Municiplal Dr. Jose de Carvalho Florence | Instituto Adolfo Lutz, Interdiciplinary Procedures Center, Strategic Laboratory | Claudio Tavares Sacchi, Claudia Regina Gonçalves, Erica Valessa Ramos Gomes, Karoline Rodrigues Campos |
