## Supplementary material for "E484K as an innovative phylogenetic event for viral evolution: Genomic analysis of the E484K spike mutation in SARS-CoV-2 lineages from Brazil": Table S3

We gratefully acknowledge the following Authors from the Originating laboratories responsible for obtaining the specimens and the Submitting laboratories where genetic sequence data were generated and shared via the GISAID Initiative, on which this research is based.

| Virus name | Accession No. | Collected | Originating Lab | Submitting Lab | Authors |
|---|---|---|---|---|---|
| Brazil/SP-01/2020 | EPI_ISL_412964 | 2020-02-25 | Hospital Israelita Albert Einstein | Instituto Adolfo Lutz Interdisciplinary Procedures Center Strategic Laboratory | Jaqueline Goes de Jesus et al |
| Brazil/SP-02/2020 | EPI_ISL_413016 | 2020-02-28 | Hospital Israelita Albert Einstein | Instituto Adolfo Lutz, Interdisciplinary Procedures Center, Strategic Laboratory | Jaqueline Goes de Jesus et al |
| Brazil/ES-225/2020 | EPI_ISL_415128 | 2020-02-29 | LACEN/ES - Laboratório Central de Saúde Pública do Espírito Santo | Instituto Oswaldo Cruz FIOCRUZ - Laboratory of Respiratory Viruses and Measles (LVRS) | Paola Resende et al |
| Brazil/SP-05/2020 | EPI_ISL_414016 | 2020-02-29 | Hospital São Joaquim Beneficencia Portuguesa | Instituto Adolfo Lutz, Interdiciplinary Procedures Center, Strategic Laboratory | Claudio Tavares Sacchi et al |
| Brazil/SP-06/2020 | EPI_ISL_414015 | 2020-02-29 | Hospital São Joaquim Beneficencia Portuguesa | Instituto Adolfo Lutz, Interdiciplinary Procedures Center, Strategic Laboratory | Claudio Tavares Sacchi et al |
| Brazil/BA-312/2020 | EPI_ISL_415105 | 2020-03-04 | Laboratório Central de Saúde Pública Professor Gonçalo Moniz – LACEN/BA | Instituto Oswaldo Cruz FIOCRUZ - Laboratory of Respiratory Viruses and Measles (LVRS) | Paola Resende et al |
| Brazil/RJ-314/2020 | EPI_ISL_414045 | 2020-03-04 | LACEN RJ - Laboratório Central de Saúde Pública Noel Nutels | Instituto Oswaldo Cruz FIOCRUZ - Laboratory of Respiratory Viruses and Measles (LVRS) | Paola Resende et al |
| Brazil/SP-10/2020 | EPI_ISL_416032 | 2020-03-04 | National Influenza Center - Instituto Adolfo Lutz | Instituto Adolfo Lutz, Interdiciplinary Procedures Center, Strategic Laboratory | Claudio Tavares Sacchi et al |
| Brazil/BA-510/2020 | EPI_ISL_427293 | 2020-03-06 | LACEN-BA - Laboratório Central de Saúde Pública Professor Gonçalo Moniz | Instituto Oswaldo Cruz FIOCRUZ - Laboratory of Respiratory Viruses and Measles (LVRS) | Paola Resende et al |
| Brazil/RS-2525/2020 | EPI_ISL_729804 | 2020-03-09 | Laboratorio Central de Saude Publica do Estado do Rio Grande do Sul (LACEN-RS) | Laboratory of Respiratory Viruses and Measles, Oswaldo Cruz Institute, FIOCRUZ | Paola Resende et al |
| Brazil/SC-766/2020 | EPI_ISL_427305 | 2020-03-10 | LACEN-SC - Laboratorio Central de Santa Catarina | Instituto Oswaldo Cruz FIOCRUZ - Laboratory of Respiratory Viruses and Measles (LVRS) | Paola Resende et al |
| Brazil/SC-769/2020 | EPI_ISL_427306 | 2020-03-10 | LACEN-SC - Laboratorio Central de Santa Catarina | Instituto Oswaldo Cruz FIOCRUZ - Laboratory of Respiratory Viruses and Measles (LVRS) | Paola Resende et al |
| Brazil/SC-770/2020 | EPI_ISL_541370 | 2020-03-12 | LACEN/SC | Laboratory of Respiratory Viruses and Measles, Oswaldo Cruz Institute, FIOCRUZ | Paola Resende et al |
| Brazil/SE-6535/2020 | EPI_ISL_541375 | 2020-03-12 | LACEN/SE | Laboratory of Respiratory Viruses and Measles, Oswaldo Cruz Institute, FIOCRUZ | Paola Resende et al |
| Brazil/DF-0001/2020 | EPI_ISL_426580 | 2020-03-13 | Instituto Sabin | Laboratory of Virology | Fernando L Melo et al |
| Brazil/DF-619i/2020 | EPI_ISL_427295 | 2020-03-13 | Instituto Oswaldo Cruz FIOCRUZ - Laboratory of Respiratory Viruses and Measles (LVRS) | Instituto Oswaldo Cruz FIOCRUZ - Laboratory of Respiratory Viruses and Measles (LVRS) | Paola Resende et al |
| Brazil/MG-CV9/2020 | EPI_ISL_429672 | 2020-03-13 | Central Public Health Laboratory/Octávio Magalhães Institute (IOM) from the Ezequiel Dias Foundation (FUNED) | Instituto Octávio Magalhães / Fundação Ezequiel Dias (IOM/Funed) | Talita Adelino et al |
| Brazil/SC-771/2020 | EPI_ISL_541371 | 2020-03-13 | LACEN/SC | Laboratory of Respiratory Viruses and Measles, Oswaldo Cruz Institute, FIOCRUZ | Paola Resende et al |
| Brazil/CE-L10-CD171/2020 | EPI_ISL_476184 | 2020-03-14 | DB Diagnósticos do Brasil | Instituto de Medicina Tropical da Univesidade de São Paulo | Samples: Nelson Gaburo Jr et al |
| Brazil/RN-IEC-162277/2020 | EPI_ISL_524798 | 2020-03-14 | Evandro Chagas Institute | Evandro Chagas Institute | Santos et al |
| Brazil/AM-02/2020 | EPI_ISL_417034 | 2020-03-16 | Laboratorio de Ecologia de Doencas Transmissiveis na Amazonia, Instituto Leonidas e Maria Deane - Fiocruz Amazonia | Laboratorio de Ecologia de Doencas Transmissiveis na Amazonia, Instituto Leonidas e Maria Deane - Fiocruz Amazonia | Valdinete Nascimento et al |
| Brazil/MG-CV16/2020 | EPI_ISL_429676 | 2020-03-16 | Central Public Health Laboratory/Octávio Magalhães Institute (IOM) from the Ezequiel Dias Foundation (FUNED) | Instituto Octávio Magalhães / Fundação Ezequiel Dias (IOM/Funed) | Talita Adelino et al |

| | | | | | |
|---|---|---|---|---|---|
| Brazil/PR-5619/2020 | EPI_ISL_541342 | 2020-03-16 | LACEN/PR | Laboratory of Respiratory Viruses and Measles, Oswaldo Cruz Institute, FIOCRUZ | Paola Resende et al |
| Brazil/RJ-L10-CD154/2020 | EPI_ISL_476178 | 2020-03-16 | DB Diagnósticos do Brasil | Instituto de Medicina Tropical da Univesidade de São Paulo | Samples: Nelson Gaburo Jr et al |
| Brazil/RJ-L10-CD158/2020 | EPI_ISL_476180 | 2020-03-16 | DB Diagnósticos do Brasil | Instituto de Medicina Tropical da Univesidade de São Paulo | Samples: Nelson Gaburo Jr et al |
| Brazil/RJ-MD63-1a/2020 | EPI_ISL_528637 | 2020-03-16 | LVM/UFRJ | Bioinformatics Laboratory / LNCC | Gustavo D. P. Silva et al |
| Brazil/RJ-MD63-1b/2020 | EPI_ISL_528638 | 2020-03-16 | LVM/UFRJ | Bioinformatics Laboratory / LNCC | Gustavo D. P. Silva et al |
| Brazil/RS-2529/2020 | EPI_ISL_729806 | 2020-03-16 | Laboratorio Central de Saude Publica do Estado do Rio Grande do Sul (LACEN-RS) | Laboratory of Respiratory Viruses and Measles, Oswaldo Cruz Institute, FIOCRUZ | Paola Resende et al |
| Brazil/RS-L11-CD180/2020 | EPI_ISL_476192 | 2020-03-16 | DB Diagnósticos do Brasil | Instituto de Medicina Tropical da Univesidade de São Paulo | Samples: Nelson Gaburo Jr et al |
| Brazil/MG-0101/2020 | EPI_ISL_417925 | 2020-03-17 | Laboratório Simili | Bioinformatics Laboratory / LNCC | Filipe Romero et al |
| Brazil/MG-CV32/2020 | EPI_ISL_429688 | 2020-03-17 | Central Public Health Laboratory/Octávio Magalhães Institute (IOM) from the Ezequiel Dias Foundation (FUNED) | Instituto Octávio Magalhães / Fundação Ezequiel Dias (IOM/Funed) | Talita Adelino et al |
| Brazil/PR-5617/2020 | EPI_ISL_541340 | 2020-03-17 | LACEN/PR | Laboratory of Respiratory Viruses and Measles, Oswaldo Cruz Institute, FIOCRUZ | Paola Resende et al |
| Brazil/PR-5618/2020 | EPI_ISL_541341 | 2020-03-17 | LACEN/PR | Laboratory of Respiratory Viruses and Measles, Oswaldo Cruz Institute, FIOCRUZ | Paola Resende et al |
| Brazil/RS-2528/2020 | EPI_ISL_729805 | 2020-03-17 | Laboratorio Central de Saude Publica do Estado do Rio Grande do Sul (LACEN-RS) | Laboratory of Respiratory Viruses and Measles, Oswaldo Cruz Institute, FIOCRUZ | Paola Resende et al |
| Brazil/SP-272/2020 | EPI_ISL_524462 | 2020-03-17 | Hospital Metropolitano | Instituto Adolfo Lutz, Interdiciplinary Procedures Center, Strategic Laboratory | Claudio Tavares Sacchi et al |
| Brazil/AC-IEC162535/2020 | EPI_ISL_458139 | 2020-03-18 | Evandro Chagas Institute | Evandro Chagas Institute | Santos et al |
| Brazil/AL-837/2020 | EPI_ISL_427292 | 2020-03-18 | LACEN-AL - Laboratorio Central de Alagoas | Instituto Oswaldo Cruz FIOCRUZ - Laboratory of Respiratory Viruses and Measles (LVRS) | Paola Resende et al |
| Brazil/MG-00301/2020 | EPI_ISL_623104 | 2020-03-18 | Simile Medicina Diagnóstica | Bioinformatics Laboratory / LNCC | Carolina M Voloch et al |
| Brazil/un-SP02cc/2020 | EPI_ISL_450506 | 2020-03-18 | Clinical Laboratory, Hospital Israelita Albert Einstein | Clinical Laboratory, Hospital Israelita Albert Einstein | Malta et al |
| Brazil/PB-IEC161853/2020 | EPI_ISL_524801 | 2020-03-19 | Evandro Chagas Institute | Evandro Chagas Institute | Santos et al |
| Brazil/PR-5621/2020 | EPI_ISL_541344 | 2020-03-19 | LACEN/PR | Laboratory of Respiratory Viruses and Measles, Oswaldo Cruz Institute, FIOCRUZ | Paola Resende et al |
| Brazil/RS-2539/2020 | EPI_ISL_729808 | 2020-03-19 | Laboratorio Central de Saude Publica do Estado do Rio Grande do Sul (LACEN-RS) | Laboratory of Respiratory Viruses and Measles, Oswaldo Cruz Institute, FIOCRUZ | Paola Resende et al |
| Brazil/SE-6533/2020 | EPI_ISL_541374 | 2020-03-19 | LACEN/SE | Laboratory of Respiratory Viruses and Measles, Oswaldo Cruz Institute, FIOCRUZ | Paola Resende et al |
| Brazil/SE-6536/2020 | EPI_ISL_541376 | 2020-03-19 | LACEN/SE | Laboratory of Respiratory Viruses and Measles, Oswaldo Cruz Institute, FIOCRUZ | Paola Resende et al |
| Brazil/SP-639/2020 | EPI_ISL_515554 | 2020-03-19 | Pronto Socorro Municipal de Perus | Instituto Adolfo Lutz, Interdiciplinary Procedures Center, Strategic Laboratory | Claudio Tavares Sacchi et al |
| Brazil/SP-L11-CD200/2020 | EPI_ISL_476201 | 2020-03-19 | DB Diagnósticos do Brasil | Instituto de Medicina Tropical da Univesidade de São Paulo | Samples: Nelson Gaburo Jr et al |
| Brazil/MG-CV42/2020 | EPI_ISL_429695 | 2020-03-20 | Central Public Health Laboratory/Octávio Magalhães Institute (IOM) from the Ezequiel Dias Foundation (FUNED) | Instituto Octávio Magalhães / Fundação Ezequiel Dias (IOM/Funed) | Talita Adelino et al |
| Brazil/PR-5620/2020 | EPI_ISL_541343 | 2020-03-20 | LACEN/PR | Laboratory of Respiratory Viruses and Measles, Oswaldo Cruz Institute, FIOCRUZ | Paola Resende et al |

| | | | | | |
|---|---|---|---|---|---|
| Brazil/RJ-763/2020 | EPI_ISL_427302 | 2020-03-20 | Instituto Oswaldo Cruz FIOCRUZ - Laboratory of Respiratory Viruses and Measles (LVRS) | Instituto Oswaldo Cruz FIOCRUZ - Laboratory of Respiratory Viruses and Measles (LVRS) | Paola Resende et al |
| Brazil/RS-2546/2020 | EPI_ISL_729810 | 2020-03-20 | Laboratorio Central de Saude Publica do Estado do Rio Grande do Sul (LACEN-RS) | Laboratory of Respiratory Viruses and Measles, Oswaldo Cruz Institute, FIOCRUZ | Paola Resende et al |
| Brazil/RS-2550/2020 | EPI_ISL_729812 | 2020-03-20 | Laboratorio Central de Saude Publica do Estado do Rio Grande do Sul (LACEN-RS) | Laboratory of Respiratory Viruses and Measles, Oswaldo Cruz Institute, FIOCRUZ | Paola Resende et al |
| Brazil/SE-6530/2020 | EPI_ISL_541373 | 2020-03-20 | LACEN/SE | Laboratory of Respiratory Viruses and Measles, Oswaldo Cruz Institute, FIOCRUZ | Paola Resende et al |
| Brazil/SP-537/2020 | EPI_ISL_471546 | 2020-03-20 | AMA DR Jose Soares Hungria | Instituto Adolfo Lutz, Interdiciplinary Procedures Center, Strategic Laboratory | Claudio Tavares Sacchi et al |
| Brazil/un-HIAE-SP04/2020 | EPI_ISL_486429 | 2020-03-20 | unknown | Clinical Laboratory, Hospital Israelita Albert Einstein | Malta et al |
| Brazil/CE-L12-CD224/2020 | EPI_ISL_476218 | 2020-03-21 | DB Diagnósticos do Brasil | Instituto de Medicina Tropical da Univesidade de São Paulo | Samples: Nelson Gaburo Jr et al |
| Brazil/DF-891/2020 | EPI_ISL_427298 | 2020-03-22 | Instituto Oswaldo Cruz FIOCRUZ - Laboratory of Respiratory Viruses and Measles (LVRS) | Instituto Oswaldo Cruz FIOCRUZ - Laboratory of Respiratory Viruses and Measles (LVRS) | Paola Resende et al |
| Brazil/RS-2549/2020 | EPI_ISL_729811 | 2020-03-22 | Laboratorio Central de Saude Publica do Estado do Rio Grande do Sul (LACEN-RS) | Laboratory of Respiratory Viruses and Measles, Oswaldo Cruz Institute, FIOCRUZ | Paola Resende et al |
| Brazil/MA-IEC162157/2020 | EPI_ISL_524797 | 2020-03-24 | Evandro Chagas Institute | Evandro Chagas Institute | Santos et al |
| Brazil/SC-L16-CD314/2020 | EPI_ISL_476278 | 2020-03-24 | DB Diagnósticos do Brasil | Instituto de Medicina Tropical da Univesidade de São Paulo | Samples: Nelson Gaburo Jr et al |
| Brazil/SP-154/2020 | EPI_ISL_515555 | 2020-03-24 | Hospital Geral de Vila Nova Cachoeirinha | Instituto Adolfo Lutz, Interdiciplinary Procedures Center, Strategic Laboratory | Claudio Tavares Sacchi et al |
| Brazil/SP-RP4/2020 | EPI_ISL_613707 | 2020-03-24 | Laboratory of Molecular Biology, Blood Center of Ribeirão Preto | Laboratory of Molecular Biology, Blood Center of Ribeirão Preto, Faculty of Medicine of Ribeirão Preto, University of São Paulo | Svetoslav N Slavov et al |
| Brazil/RJ-1111/2020 | EPI_ISL_456076 | 2020-03-25 | LACEN RJ - Laboratório Central de Saúde Pública Noel Nutels | Laboratory of Respiratory Viruses and Measles, Oswaldo Cruz Institute, FIOCRUZ | Paola Resende et al |
| Brazil/RS-2553/2020 | EPI_ISL_729813 | 2020-03-25 | Laboratorio Central de Saude Publica do Estado do Rio Grande do Sul (LACEN-RS) | Laboratory of Respiratory Viruses and Measles, Oswaldo Cruz Institute, FIOCRUZ | Paola Resende et al |
| Brazil/RS-2554/2020 | EPI_ISL_729814 | 2020-03-25 | Laboratorio Central de Saude Publica do Estado do Rio Grande do Sul (LACEN-RS) | Laboratory of Respiratory Viruses and Measles, Oswaldo Cruz Institute, FIOCRUZ | Paola Resende et al |
| Brazil/SP-190/2020 | EPI_ISL_523982 | 2020-03-25 | Hospital do Servidor Público Estadual Francisco Morato de Oliveira | Instituto Adolfo Lutz, Interdiciplinary Procedures Center, Strategic Laboratory | Claudio Tavares Sacchi et al |
| Brazil/SP-625/2020 | EPI_ISL_515528 | 2020-03-25 | Hospital Sao Paulo de Ensino da Unifesp | Instituto Adolfo Lutz, Interdiciplinary Procedures Center, Strategic Laboratory | Claudio Tavares Sacchi et al |
| Brazil/PR-5623/2020 | EPI_ISL_541346 | 2020-03-26 | LACEN/PR | Laboratory of Respiratory Viruses and Measles, Oswaldo Cruz Institute, FIOCRUZ | Paola Resende et al |
| Brazil/SP-147/2020 | EPI_ISL_468312 | 2020-03-26 | Hospital Municipal Dr Ignacio Proenca de Gouvea | Instituto Adolfo Lutz, Interdiciplinary Procedures Center, Strategic Laboratory | Claudio Tavares Sacchi et al |
| Brazil/SP-149/2020 | EPI_ISL_468314 | 2020-03-26 | CTA Centro de Testagem e Aconselhamento | Instituto Adolfo Lutz, Interdiciplinary Procedures Center, Strategic Laboratory | Claudio Tavares Sacchi et al |
| Brazil/RS-2556/2020 | EPI_ISL_729815 | 2020-03-27 | Laboratorio Central de Saude Publica do Estado do Rio Grande do Sul (LACEN-RS) | Laboratory of Respiratory Viruses and Measles, Oswaldo Cruz Institute, FIOCRUZ | Paola Resende et al |
| Brazil/SE-6529/2020 | EPI_ISL_541372 | 2020-03-27 | LACEN/SE | Laboratory of Respiratory Viruses and Measles, Oswaldo Cruz Institute, FIOCRUZ | Paola Resende et al |
| Brazil/PR-5622/2020 | EPI_ISL_541345 | 2020-03-28 | LACEN/PR | Laboratory of Respiratory Viruses and Measles, Oswaldo Cruz Institute, FIOCRUZ | Paola Resende et al |
| Brazil/PR-L16-CD333/2020 | EPI_ISL_476282 | 2020-03-28 | DB Diagnósticos do Brasil | Instituto de Medicina Tropical da Univesidade de São Paulo | Samples: Nelson Gaburo Jr et al |

| | | | | | |
|---|---|---|---|---|---|
| Brazil/RJ-0920/2020 | EPI_ISL_636838 | 2020-03-29 | Laboratório de Imunofarmacologia - Instituto Oswaldo Cruz | Laboratório de Imunofarmacologia - Instituto Oswaldo Cruz | Souza et al |
| Brazil/SP-551/2020 | EPI_ISL_471554 | 2020-03-29 | Hospital Bosque da Saúde | Instituto Adolfo Lutz, Interdiciplinary Procedures Center, Strategic Laboratory | Claudio Tavares Sacchi et al |
| Brazil/CE-L16-CD344/2020 | EPI_ISL_476288 | 2020-03-30 | DB Diagnósticos do Brasil | Instituto de Medicina Tropical da Univesidade de São Paulo | Samples: Nelson Gaburo Jr et al |
| Brazil/CE-L16-CD346/2020 | EPI_ISL_476289 | 2020-03-30 | DB Diagnósticos do Brasil | Instituto de Medicina Tropical da Univesidade de São Paulo | Samples: Nelson Gaburo Jr et al |
| Brazil/RJ-899/2020 | EPI_ISL_456071 | 2020-03-30 | Laboratory of Respiratory Viruses and Measles, Oswaldo Cruz Institute, FIOCRUZ | Laboratory of Respiratory Viruses and Measles, Oswaldo Cruz Institute, FIOCRUZ | Paola Resende et al |
| Brazil/SP-294/2020 | EPI_ISL_527862 | 2020-03-30 | Hospital Municipal de Urgência | Instituto Adolfo Lutz, Interdiciplinary Procedures Center, Strategic Laboratory | Claudio Tavares Sacchi et al |
| Brazil/SP-122/2020 | EPI_ISL_515522 | 2020-03-31 | UPA 24HS de Itatiba | Instituto Adolfo Lutz, Interdiciplinary Procedures Center, Strategic Laboratory | Claudio Tavares Sacchi et al |
| Brazil/SP-125/2020 | EPI_ISL_515524 | 2020-04-01 | PS Municipal Dr Lauro Ribas Braga | Instituto Adolfo Lutz, Interdiciplinary Procedures Center, Strategic Laboratory | Claudio Tavares Sacchi et al |
| Brazil/SP-L22-CD474/2020 | EPI_ISL_476373 | 2020-04-01 | Hospital da Clínicas da Faculdade de Medicina da Universidade de São Paulo | Instituto de Medicina Tropical da Univesidade de São Paulo | Samples: Ingra Morales Claro et al |
| Brazil/CE-0206/2020 | EPI_ISL_470573 | 2020-04-02 | Hermes Pardini | Bioinformatics Laboratory / LNCC | Alexandra Gerber et al |
| Brazil/SP-124/2020 | EPI_ISL_515523 | 2020-04-02 | PS Municipal Dr Lauro Ribas Braga | Instituto Adolfo Lutz, Interdiciplinary Procedures Center, Strategic Laboratory | Claudio Tavares Sacchi et al |
| Brazil/RS-2564/2020 | EPI_ISL_729816 | 2020-04-03 | Laboratorio Central de Saude Publica do Estado do Rio Grande do Sul (LACEN-RS) | Laboratory of Respiratory Viruses and Measles, Oswaldo Cruz Institute, FIOCRUZ | Paola Resende et al |
| Brazil/AP-IEC162966/2020 | EPI_ISL_458142 | 2020-04-05 | Evandro Chagas Institute | Evandro Chagas Institute | Santos et al |
| Brazil/PA-IEC162802/2020 | EPI_ISL_458140 | 2020-04-07 | Evandro Chagas Institute | Evandro Chagas Institute | Santos et al |
| Brazil/PE-IAM08/2020 | EPI_ISL_500460 | 2020-04-07 | LACEN/PE | WallauLab, Aggeu Magalhaes Institute | Marcelo Henrique Santos Paiva et al |
| Brazil/PE-IAM89/2020 | EPI_ISL_500486 | 2020-04-07 | LACEN/PE | WallauLab, Aggeu Magalhaes Institute | Marcelo Henrique Santos Paiva et al |
| Brazil/PE-IAM109/2020 | EPI_ISL_572334 | 2020-04-08 | LACEN/PE | WallauLab, Aggeu Magalhaes Institute | Marcelo Henrique Santos Paiva et al |
| Brazil/PE-IAM221/2020 | EPI_ISL_500473 | 2020-04-08 | LACEN/PE | WallauLab, Aggeu Magalhaes Institute | Marcelo Henrique Santos Paiva et al |
| Brazil/PE-IAM30/2020 | EPI_ISL_500477 | 2020-04-08 | LACEN/PE | WallauLab, Aggeu Magalhaes Institute | Marcelo Henrique Santos Paiva et al |
| Brazil/PE-IAM48/2020 | EPI_ISL_500482 | 2020-04-08 | LACEN/PE | WallauLab, Aggeu Magalhaes Institute | Marcelo Henrique Santos Paiva et al |
| Brazil/PE-IAM67/2020 | EPI_ISL_500483 | 2020-04-08 | LACEN/PE | WallauLab, Aggeu Magalhaes Institute | Marcelo Henrique Santos Paiva et al |
| Brazil/PE-IAM84/2020 | EPI_ISL_500484 | 2020-04-08 | LACEN/PE | WallauLab, Aggeu Magalhaes Institute | Marcelo Henrique Santos Paiva et al |
| Brazil/PE-IAM87/2020 | EPI_ISL_500485 | 2020-04-08 | LACEN/PE | WallauLab, Aggeu Magalhaes Institute | Marcelo Henrique Santos Paiva et al |
| Brazil/RJ-INCA-C07/2020 | EPI_ISL_513514 | 2020-04-08 | Programa de Oncovirologia, Instituto Nacional de Câncer | Programa de Oncovirologia, Instituto Nacional de Câncer | Juliana D. Siqueira et al |
| Brazil/AP-IEC163359/2020 | EPI_ISL_524795 | 2020-04-09 | Evandro Chagas Institute | Evandro Chagas Institute | Santos et al |
| Brazil/PE-IAM103/2020 | EPI_ISL_500462 | 2020-04-09 | LACEN/PE | WallauLab, Aggeu Magalhaes Institute | Marcelo Henrique Santos Paiva et al |

| | | | | | |
|---|---|---|---|---|---|
| Brazil/PE-IAM138/2020 | EPI_ISL_500463 | 2020-04-09 | LACEN/PE | WallauLab, Aggeu Magalhaes Institute | Marcelo Henrique Santos Paiva et al |
| Brazil/MG-00302/2020 | EPI_ISL_623105 | 2020-04-13 | Simile Medicina Diagnóstica | Bioinformatics Laboratory / LNCC | Carolina M Voloch et al |
| Brazil/PE-IAM355/2020 | EPI_ISL_500874 | 2020-04-13 | LACEN/PE | WallauLab, Aggeu Magalhaes Institute | Marcelo Henrique Santos Paiva et al |
| Brazil/RJ-1953/2020 | EPI_ISL_541352 | 2020-04-13 | Laboratory of Respiratory Viruses and Measles, Oswaldo Cruz Institute, FIOCRUZ | Laboratory of Respiratory Viruses and Measles, Oswaldo Cruz Institute, FIOCRUZ | Paola Resende et al |
| Brazil/AP-IEC163972/2020 | EPI_ISL_458144 | 2020-04-15 | Evandro Chagas Institute | Evandro Chagas Institute | Santos et al |
| Brazil/AP-IEC164082/2020 | EPI_ISL_458143 | 2020-04-15 | Evandro Chagas Institute | Evandro Chagas Institute | Santos et al |
| Brazil/PA-IEC163469/2020 | EPI_ISL_524787 | 2020-04-15 | Evandro Chagas Institute | Evandro Chagas Institute | Santos et al |
| Brazil/SP-459/2020 | EPI_ISL_583490 | 2020-04-15 | Hospital Estadual Sumare | Instituto Adolfo Lutz, Interdiciplinary Procedures Center, Strategic Laboratory | Claudio Tavares Sacchi et al |
| Brazil/SP-461/2020 | EPI_ISL_583492 | 2020-04-15 | Santa Casa Anna Cintra | Instituto Adolfo Lutz, Interdiciplinary Procedures Center, Strategic Laboratory | Claudio Tavares Sacchi et al |
| Brazil/RJ-INCA-C22/2020 | EPI_ISL_513530 | 2020-04-16 | Programa de Oncovirologia, Instituto Nacional de Câncer | Programa de Oncovirologia, Instituto Nacional de Câncer | Juliana D. Siqueira et al |
| Brazil/RJ-INCA-C24/2020 | EPI_ISL_513532 | 2020-04-16 | Programa de Oncovirologia, Instituto Nacional de Câncer | Programa de Oncovirologia, Instituto Nacional de Câncer | Juliana D. Siqueira et al |
| Brazil/RJ-00337/2020 | EPI_ISL_623140 | 2020-04-17 | Laboratorio de Virologia Molecular / UFRJ | Bioinformatics Laboratory / LNCC | Carolina M Voloch et al |
| Brazil/RJ-INCA-C41/2020 | EPI_ISL_513549 | 2020-04-17 | Programa de Oncovirologia, Instituto Nacional de Câncer | Programa de Oncovirologia, Instituto Nacional de Câncer | Juliana D. Siqueira et al |
| Brazil/SP-283/2020 | EPI_ISL_527860 | 2020-04-17 | Hospital Municipal de Parelheiros Josanias Castanha Braga | Instituto Adolfo Lutz, Interdiciplinary Procedures Center, Strategic Laboratory | Claudio Tavares Sacchi et al |
| Brazil/SP-209/2020 | EPI_ISL_523990 | 2020-04-19 | AMA Jardim Peri | Instituto Adolfo Lutz, Interdiciplinary Procedures Center, Strategic Laboratory | Claudio Tavares Sacchi et al |
| Brazil/PE-IAM383/2020 | EPI_ISL_572335 | 2020-04-20 | LACEN/PE | WallauLab, Aggeu Magalhaes Institute | Marcelo Henrique Santos Paiva et al |
| Brazil/RS-6213/2020 | EPI_ISL_729834 | 2020-04-20 | Laboratorio Central de Saude Publica do Estado do Rio Grande do Sul (LACEN-RS) | Laboratory of Respiratory Viruses and Measles, Oswaldo Cruz Institute, FIOCRUZ | Paola Resende et al |
| Brazil/SE-6583/2020 | EPI_ISL_541389 | 2020-04-20 | LACEN/SE | Laboratory of Respiratory Viruses and Measles, Oswaldo Cruz Institute, FIOCRUZ | Paola Resende et al |
| Brazil/SE-6601/2020 | EPI_ISL_541392 | 2020-04-20 | LACEN/SE | Laboratory of Respiratory Viruses and Measles, Oswaldo Cruz Institute, FIOCRUZ | Paola Resende et al |
| Brazil/SE-6606/2020 | EPI_ISL_541394 | 2020-04-20 | LACEN/SE | Laboratory of Respiratory Viruses and Measles, Oswaldo Cruz Institute, FIOCRUZ | Paola Resende et al |
| Brazil/SE-6607/2020 | EPI_ISL_541395 | 2020-04-20 | LACEN/SE | Laboratory of Respiratory Viruses and Measles, Oswaldo Cruz Institute, FIOCRUZ | Paola Resende et al |
| Brazil/SE-6608/2020 | EPI_ISL_541396 | 2020-04-20 | LACEN/SE | Laboratory of Respiratory Viruses and Measles, Oswaldo Cruz Institute, FIOCRUZ | Paola Resende et al |
| Brazil/SE-6603/2020 | EPI_ISL_541393 | 2020-04-21 | LACEN/SE | Laboratory of Respiratory Viruses and Measles, Oswaldo Cruz Institute, FIOCRUZ | Paola Resende et al |
| Brazil/RJ-00327/2020 | EPI_ISL_623130 | 2020-04-22 | Laboratorio de Virologia Molecular / UFRJ | Bioinformatics Laboratory / LNCC | Carolina M Voloch et al |
| Brazil/RJ-00339/2020 | EPI_ISL_623142 | 2020-04-22 | Laboratorio de Virologia Molecular / UFRJ | Bioinformatics Laboratory / LNCC | Carolina M Voloch et al |
| Brazil/RJ-INCA-C64/2020 | EPI_ISL_513557 | 2020-04-22 | Programa de Oncovirologia, Instituto Nacional de Câncer | Programa de Oncovirologia, Instituto Nacional de Câncer | Juliana D. Siqueira et al |
| Brazil/RJ-INCA-C69/2020 | EPI_ISL_513561 | 2020-04-22 | Programa de Oncovirologia, Instituto Nacional de Câncer | Programa de Oncovirologia, Instituto Nacional de Câncer | Juliana D. Siqueira et al |

| | | | | | |
|---|---|---|---|---|---|
| Brazil/SP-516/2020 | EPI_ISL_468321 | 2020-04-22 | Hospital Universitario da USP | Instituto Adolfo Lutz, Interdiciplinary Procedures Center, Strategic Laboratory | Claudio Tavares Sacchi et al |
| Brazil/PA-IEC164173/2020 | EPI_ISL_458146 | 2020-04-23 | Evandro Chagas Institute | Evandro Chagas Institute | Santos et al |
| Brazil/SP-203/2020 | EPI_ISL_523986 | 2020-04-23 | Ama Dr Jose Soares Hungria | Instituto Adolfo Lutz, Interdiciplinary Procedures Center, Strategic Laboratory | Claudio Tavares Sacchi et al |
| Brazil/MA-IEC164827/2020 | EPI_ISL_524788 | 2020-04-24 | Evandro Chagas Institute | Evandro Chagas Institute | Santos et al |
| Brazil/PA-IEC164218/2020 | EPI_ISL_458147 | 2020-04-24 | Evandro Chagas Institute | Evandro Chagas Institute | Santos et al |
| Brazil/RJ-00311/2020 | EPI_ISL_623114 | 2020-04-24 | Laboratorio de Virologia Molecular / UFRJ | Bioinformatics Laboratory / LNCC | Carolina M Voloch et al |
| Brazil/RJ-INCA-C77/2020 | EPI_ISL_513567 | 2020-04-24 | Programa de Oncovirologia, Instituto Nacional de Câncer | Programa de Oncovirologia, Instituto Nacional de Câncer | Juliana D. Siqueira et al |
| Brazil/SE-6567/2020 | EPI_ISL_541386 | 2020-04-24 | LACEN/SE | Laboratory of Respiratory Viruses and Measles, Oswaldo Cruz Institute, FIOCRUZ | Paola Resende et al |
| Brazil/SE-6568/2020 | EPI_ISL_541387 | 2020-04-24 | LACEN/SE | Laboratory of Respiratory Viruses and Measles, Oswaldo Cruz Institute, FIOCRUZ | Paola Resende et al |
| Brazil/MA-IEC165425/2020 | EPI_ISL_524794 | 2020-04-25 | Evandro Chagas Institute | Evandro Chagas Institute | Santos et al |
| Brazil/PA-IEC164239/2020 | EPI_ISL_458141 | 2020-04-26 | Evandro Chagas Institute | Evandro Chagas Institute | Santos et al |
| Brazil/SE-6539/2020 | EPI_ISL_541377 | 2020-04-26 | LACEN/SE | Laboratory of Respiratory Viruses and Measles, Oswaldo Cruz Institute, FIOCRUZ | Paola Resende et al |
| Brazil/SE-6544/2020 | EPI_ISL_541378 | 2020-04-26 | LACEN/SE | Laboratory of Respiratory Viruses and Measles, Oswaldo Cruz Institute, FIOCRUZ | Paola Resende et al |
| Brazil/SE-6549/2020 | EPI_ISL_541379 | 2020-04-26 | LACEN/SE | Laboratory of Respiratory Viruses and Measles, Oswaldo Cruz Institute, FIOCRUZ | Paola Resende et al |
| Brazil/SE-6563/2020 | EPI_ISL_541385 | 2020-04-26 | LACEN/SE | Laboratory of Respiratory Viruses and Measles, Oswaldo Cruz Institute, FIOCRUZ | Paola Resende et al |
| Brazil/PA-IEC164684/2020 | EPI_ISL_458148 | 2020-04-27 | Evandro Chagas Institute | Evandro Chagas Institute | Santos et al |
| Brazil/PE-IAM719/2020 | EPI_ISL_572353 | 2020-04-27 | LACEN/PE | WallauLab, Aggeu Magalhaes Institute | Marcelo Henrique Santos Paiva et al |
| Brazil/SE-6550/2020 | EPI_ISL_541380 | 2020-04-27 | LACEN/SE | Laboratory of Respiratory Viruses and Measles, Oswaldo Cruz Institute, FIOCRUZ | Paola Resende et al |
| Brazil/SE-6555/2020 | EPI_ISL_541381 | 2020-04-27 | LACEN/SE | Laboratory of Respiratory Viruses and Measles, Oswaldo Cruz Institute, FIOCRUZ | Paola Resende et al |
| Brazil/SE-6556/2020 | EPI_ISL_541382 | 2020-04-27 | LACEN/SE | Laboratory of Respiratory Viruses and Measles, Oswaldo Cruz Institute, FIOCRUZ | Paola Resende et al |
| Brazil/SE-6557/2020 | EPI_ISL_541383 | 2020-04-27 | LACEN/SE | Laboratory of Respiratory Viruses and Measles, Oswaldo Cruz Institute, FIOCRUZ | Paola Resende et al |
| Brazil/SE-6561/2020 | EPI_ISL_541384 | 2020-04-27 | LACEN/SE | Laboratory of Respiratory Viruses and Measles, Oswaldo Cruz Institute, FIOCRUZ | Paola Resende et al |
| Brazil/MA-IEC165398/2020 | EPI_ISL_524790 | 2020-04-28 | Evandro Chagas Institute | Evandro Chagas Institute | Santos et al |
| Brazil/PA-IEC164747/2020 | EPI_ISL_524783 | 2020-04-28 | Evandro Chagas Institute | Evandro Chagas Institute | Santos et al |
| Brazil/MA-IEC162466/2020 | EPI_ISL_524791 | 2020-04-29 | Evandro Chagas Institute | Evandro Chagas Institute | Santos et al |
| Brazil/RJ-00359/2020 | EPI_ISL_623162 | 2020-04-29 | Laboratorio de Virologia Molecular / UFRJ | Bioinformatics Laboratory / LNCC | Carolina M Voloch et al |

| | | | | | |
|---|---|---|---|---|---|
| Brazil/SE-6591/2020 | EPI_ISL_541390 | 2020-04-29 | LACEN/SE | Laboratory of Respiratory Viruses and Measles, Oswaldo Cruz Institute, FIOCRUZ | Paola Resende et al |
| Brazil/SE-6594/2020 | EPI_ISL_541391 | 2020-04-29 | LACEN/SE | Laboratory of Respiratory Viruses and Measles, Oswaldo Cruz Institute, FIOCRUZ | Paola Resende et al |
| Brazil/PA-IEC165302/2020 | EPI_ISL_524786 | 2020-05-01 | Evandro Chagas Institute | Evandro Chagas Institute | Santos et al |
| Brazil/PE-IAM961/2020 | EPI_ISL_572366 | 2020-05-02 | LACEN/PE | WallauLab, Aggeu Magalhaes Institute | Marcelo Henrique Santos Paiva et al |
| Brazil/PE-IAM965/2020 | EPI_ISL_572367 | 2020-05-02 | LACEN/PE | WallauLab, Aggeu Magalhaes Institute | Marcelo Henrique Santos Paiva et al |
| Brazil/SP-243/2020 | EPI_ISL_693196 | 2020-05-02 | Hospital Santa Clara | Instituto Adolfo Lutz, Interdiciplinary Procedures Center, Strategic Laboratory | Claudio Tavares Sacchi et al |
| Brazil/SP-321/2020 | EPI_ISL_534316 | 2020-05-02 | OS Mun Santana Lauro Ribas Braga | Instituto Adolfo Lutz, Interdiciplinary Procedures Center, Strategic Laboratory | Claudio Tavares Sacchi et al |
| Brazil/PE-IAM1082/2020 | EPI_ISL_572375 | 2020-05-03 | LACEN/PE | WallauLab, Aggeu Magalhaes Institute | Marcelo Henrique Santos Paiva et al |
| Brazil/PE-IAM889/2020 | EPI_ISL_572358 | 2020-05-03 | LACEN/PE | WallauLab, Aggeu Magalhaes Institute | Marcelo Henrique Santos Paiva et al |
| Brazil/PE-IAM900/2020 | EPI_ISL_572359 | 2020-05-03 | LACEN/PE | WallauLab, Aggeu Magalhaes Institute | Marcelo Henrique Santos Paiva et al |
| Brazil/PE-IAM914/2020 | EPI_ISL_572360 | 2020-05-03 | LACEN/PE | WallauLab, Aggeu Magalhaes Institute | Marcelo Henrique Santos Paiva et al |
| Brazil/PE-IAM939/2020 | EPI_ISL_572361 | 2020-05-03 | LACEN/PE | WallauLab, Aggeu Magalhaes Institute | Marcelo Henrique Santos Paiva et al |
| Brazil/RJ-00316/2020 | EPI_ISL_623119 | 2020-05-04 | Laboratorio de Virologia Molecular / UFRJ | Bioinformatics Laboratory / LNCC | Carolina M Voloch et al |
| Brazil/RJ-INCA-I34/2020 | EPI_ISL_513575 | 2020-05-04 | Programa de Oncovirologia, Instituto Nacional de Câncer | Programa de Oncovirologia, Instituto Nacional de Câncer | Juliana D. Siqueira et al |
| Brazil/RJ-INCA-I35/2020 | EPI_ISL_513576 | 2020-05-04 | Programa de Oncovirologia, Instituto Nacional de Câncer | Programa de Oncovirologia, Instituto Nacional de Câncer | Juliana D. Siqueira et al |
| Brazil/RJ-INCA-I39/2020 | EPI_ISL_513578 | 2020-05-04 | Programa de Oncovirologia, Instituto Nacional de Câncer | Programa de Oncovirologia, Instituto Nacional de Câncer | Juliana D. Siqueira et al |
| Brazil/RJ-INCA-I43/2020 | EPI_ISL_513579 | 2020-05-04 | Programa de Oncovirologia, Instituto Nacional de Câncer | Programa de Oncovirologia, Instituto Nacional de Câncer | Juliana D. Siqueira et al |
| Brazil/SP-249/2020 | EPI_ISL_693198 | 2020-05-04 | Santa Casa de Misericordia de Sao Paulo - Hospital Central | Instituto Adolfo Lutz, Interdiciplinary Procedures Center, Strategic Laboratory | Claudio Tavares Sacchi et al |
| Brazil/SP-317/2020 | EPI_ISL_534314 | 2020-05-04 | Hospital Universitario da USP de SP | Instituto Adolfo Lutz, Interdiciplinary Procedures Center, Strategic Laboratory | Claudio Tavares Sacchi et al |
| Brazil/MA-IEC166716/2020 | EPI_ISL_524789 | 2020-05-05 | Evandro Chagas Institute | Evandro Chagas Institute | Santos et al |
| Brazil/PA-IEC165313/2020 | EPI_ISL_524785 | 2020-05-05 | Evandro Chagas Institute | Evandro Chagas Institute | Santos et al |
| Brazil/RJ-00362/2020 | EPI_ISL_623165 | 2020-05-05 | Laboratorio de Virologia Molecular / UFRJ | Bioinformatics Laboratory / LNCC | Carolina M Voloch et al |
| Brazil/RJ-INCA-I51/2020 | EPI_ISL_513581 | 2020-05-05 | Programa de Oncovirologia, Instituto Nacional de Câncer | Programa de Oncovirologia, Instituto Nacional de Câncer | Juliana D. Siqueira et al |
| Brazil/RO-06/2020 | EPI_ISL_514136 | 2020-05-05 | Rondônia Central Public Health Laboratory (LACEN/RO), vinculated to State Health Secretariat of Rondônia (SESAU/RO) | Molecular Virology Laboratory of Oswaldo Cruz Foundation of Rondônia | Luan Felipo Botelho-Souza et al |
| Brazil/RO-07/2020 | EPI_ISL_514137 | 2020-05-05 | Rondônia Central Public Health Laboratory (LACEN/RO), vinculated to State Health Secretariat of Rondônia (SESAU/RO) | Molecular Virology Laboratory of Oswaldo Cruz Foundation of Rondônia | Luan Felipo Botelho-Souza et al |

| | | | | | |
|---|---|---|---|---|---|
| Brazil/SP-256/2020 | EPI_ISL_534320 | 2020-05-05 | Hospital do Serv Pub ESTAFCO Morato de Oliveira | Instituto Adolfo Lutz, Interdiciplinary Procedures Center, Strategic Laboratory | Claudio Tavares Sacchi et al |
| Brazil/SP-324/2020 | EPI_ISL_534317 | 2020-05-05 | Hospital Geral de Itapevi | Instituto Adolfo Lutz, Interdiciplinary Procedures Center, Strategic Laboratory | Claudio Tavares Sacchi et al |
| Brazil/PE-IAM1148/2020 | EPI_ISL_572381 | 2020-05-06 | LACEN/PE | WallauLab, Aggeu Magalhaes Institute | Marcelo Henrique Santos Paiva et al |
| Brazil/PE-IAM991/2020 | EPI_ISL_572371 | 2020-05-06 | LACEN/PE | WallauLab, Aggeu Magalhaes Institute | Marcelo Henrique Santos Paiva et al |
| Brazil/PE-IAM992/2020 | EPI_ISL_572372 | 2020-05-06 | LACEN/PE | WallauLab, Aggeu Magalhaes Institute | Marcelo Henrique Santos Paiva et al |
| Brazil/RS-6177/2020 | EPI_ISL_729820 | 2020-05-06 | Laboratorio Central de Saude Publica do Estado do Rio Grande do Sul (LACEN-RS) | Laboratory of Respiratory Viruses and Measles, Oswaldo Cruz Institute, FIOCRUZ | Paola Resende et al |
| Brazil/RS-6180/2020 | EPI_ISL_729797 | 2020-05-06 | Laboratorio Central de Saude Publica do Estado do Rio Grande do Sul (LACEN-RS) | Laboratory of Respiratory Viruses and Measles, Oswaldo Cruz Institute, FIOCRUZ | Paola Resende et al |
| Brazil/RS-6183/2020 | EPI_ISL_729822 | 2020-05-06 | Laboratorio Central de Saude Publica do Estado do Rio Grande do Sul (LACEN-RS) | Laboratory of Respiratory Viruses and Measles, Oswaldo Cruz Institute, FIOCRUZ | Paola Resende et al |
| Brazil/SP-255/2020 | EPI_ISL_534319 | 2020-05-06 | Hospital do Serv Pub ESTAFCO Morato de Oliveira | Instituto Adolfo Lutz, Interdiciplinary Procedures Center, Strategic Laboratory | Claudio Tavares Sacchi et al |
| Brazil/MA-IEC166867/2020 | EPI_ISL_524799 | 2020-05-08 | Evandro Chagas Institute | Evandro Chagas Institute | Santos et al |
| Brazil/PR-LRV-01/2020 | EPI_ISL_510535 | 2020-05-08 | Molecular Virology, Instituto Carlos Chagas / Fiocruz Paraná | Universidade Federal do Parana (UFPR) | Suzukawa et al |
| Brazil/RJ-00346/2020 | EPI_ISL_623149 | 2020-05-08 | Laboratorio de Virologia Molecular / UFRJ | Bioinformatics Laboratory / LNCC | Carolina M Voloch et al |
| Brazil/RS-6184/2020 | EPI_ISL_729823 | 2020-05-08 | Laboratorio Central de Saude Publica do Estado do Rio Grande do Sul (LACEN-RS) | Laboratory of Respiratory Viruses and Measles, Oswaldo Cruz Institute, FIOCRUZ | Paola Resende et al |
| Brazil/PE-IAM1254/2020 | EPI_ISL_572384 | 2020-05-09 | LACEN/PE | WallauLab, Aggeu Magalhaes Institute | Marcelo Henrique Santos Paiva et al |
| Brazil/RS-6187/2020 | EPI_ISL_729798 | 2020-05-09 | Laboratorio Central de Saude Publica do Estado do Rio Grande do Sul (LACEN-RS) | Laboratory of Respiratory Viruses and Measles, Oswaldo Cruz Institute, FIOCRUZ | Paola Resende et al |
| Brazil/PE-IAM1309/2020 | EPI_ISL_572386 | 2020-05-10 | LACEN/PE | WallauLab, Aggeu Magalhaes Institute | Marcelo Henrique Santos Paiva et al |
| Brazil/RS-6188/2020 | EPI_ISL_729824 | 2020-05-10 | Laboratorio Central de Saude Publica do Estado do Rio Grande do Sul (LACEN-RS) | Laboratory of Respiratory Viruses and Measles, Oswaldo Cruz Institute, FIOCRUZ | Paola Resende et al |
| Brazil/RS-6189/2020 | EPI_ISL_729825 | 2020-05-10 | Laboratorio Central de Saude Publica do Estado do Rio Grande do Sul (LACEN-RS) | Laboratory of Respiratory Viruses and Measles, Oswaldo Cruz Institute, FIOCRUZ | Paola Resende et al |
| Brazil/PE-IAM1315/2020 | EPI_ISL_572388 | 2020-05-11 | LACEN/PE | WallauLab, Aggeu Magalhaes Institute | Marcelo Henrique Santos Paiva et al |
| Brazil/RJ-00340/2020 | EPI_ISL_623143 | 2020-05-11 | Laboratorio de Virologia Molecular / UFRJ | Bioinformatics Laboratory / LNCC | Carolina M Voloch et al |
| Brazil/RO-01/2020 | EPI_ISL_514131 | 2020-05-11 | Rondônia Central Public Health Laboratory (LACEN/RO), vinculated to State Health Secretariat of Rondônia (SESAU/RO) | Molecular Virology Laboratory of Oswaldo Cruz Foundation of Rondônia | Luan Felipo Botelho-Souza et al |
| Brazil/RO-02/2020 | EPI_ISL_514132 | 2020-05-11 | Rondônia Central Public Health Laboratory (LACEN/RO), vinculated to State Health Secretariat of Rondônia (SESAU/RO) | Molecular Virology Laboratory of Oswaldo Cruz Foundation of Rondônia | Luan Felipo Botelho-Souza et al |
| Brazil/RO-04/2020 | EPI_ISL_514134 | 2020-05-11 | Rondônia Central Public Health Laboratory (LACEN/RO), vinculated to State Health Secretariat of Rondônia (SESAU/RO) | Molecular Virology Laboratory of Oswaldo Cruz Foundation of Rondônia | Luan Felipo Botelho-Souza et al |
| Brazil/RS-6195/2020 | EPI_ISL_729794 | 2020-05-11 | Laboratorio Central de Saude Publica do Estado do Rio Grande do Sul (LACEN-RS) | Laboratory of Respiratory Viruses and Measles, Oswaldo Cruz Institute, FIOCRUZ | Paola Resende et al |
| Brazil/RS-6192/2020 | EPI_ISL_729827 | 2020-05-12 | Laboratorio Central de Saude Publica do Estado do Rio Grande do Sul (LACEN-RS) | Laboratory of Respiratory Viruses and Measles, Oswaldo Cruz Institute, FIOCRUZ | Paola Resende et al |

| | | | | | |
|---|---|---|---|---|---|
| Brazil/PA-IEC166687/2020 | EPI_ISL_524800 | 2020-05-13 | Evandro Chagas Institute | Evandro Chagas Institute | Santos et al |
| Brazil/PE-IAM1468/2020 | EPI_ISL_572396 | 2020-05-14 | LACEN/PE | WallauLab, Aggeu Magalhaes Institute | Marcelo Henrique Santos Paiva et al |
| Brazil/RS-6203/2020 | EPI_ISL_729831 | 2020-05-14 | Laboratorio Central de Saude Publica do Estado do Rio Grande do Sul (LACEN-RS) | Laboratory of Respiratory Viruses and Measles, Oswaldo Cruz Institute, FIOCRUZ | Paola Resende et al |
| Brazil/RS-6205/2020 | EPI_ISL_729832 | 2020-05-18 | Laboratorio Central de Saude Publica do Estado do Rio Grande do Sul (LACEN-RS) | Laboratory of Respiratory Viruses and Measles, Oswaldo Cruz Institute, FIOCRUZ | Paola Resende et al |
| Brazil/RS-6208/2020 | EPI_ISL_729833 | 2020-05-19 | Laboratorio Central de Saude Publica do Estado do Rio Grande do Sul (LACEN-RS) | Laboratory of Respiratory Viruses and Measles, Oswaldo Cruz Institute, FIOCRUZ | Paola Resende et al |
| Brazil/RS-6215/2020 | EPI_ISL_729835 | 2020-05-20 | Laboratorio Central de Saude Publica do Estado do Rio Grande do Sul (LACEN-RS) | Laboratory of Respiratory Viruses and Measles, Oswaldo Cruz Institute, FIOCRUZ | Paola Resende et al |
| Brazil/SP-253/2020 | EPI_ISL_693199 | 2020-05-20 | Hospital do Servidor Publico Estadual Francisco Morato de Oliveira | Instituto Adolfo Lutz, Interdiciplinary Procedures Center, Strategic Laboratory | Claudio Tavares Sacchi et al |
| Brazil/RS-6218/2020 | EPI_ISL_729836 | 2020-05-27 | Laboratorio Central de Saude Publica do Estado do Rio Grande do Sul (LACEN-RS) | Laboratory of Respiratory Viruses and Measles, Oswaldo Cruz Institute, FIOCRUZ | Paola Resende et al |
| Brazil/RS-6219/2020 | EPI_ISL_729795 | 2020-05-27 | Laboratorio Central de Saude Publica do Estado do Rio Grande do Sul (LACEN-RS) | Laboratory of Respiratory Viruses and Measles, Oswaldo Cruz Institute, FIOCRUZ | Paola Resende et al |
| Brazil/RS-6222/2020 | EPI_ISL_729837 | 2020-05-28 | Laboratorio Central de Saude Publica do Estado do Rio Grande do Sul (LACEN-RS) | Laboratory of Respiratory Viruses and Measles, Oswaldo Cruz Institute, FIOCRUZ | Paola Resende et al |
| Brazil/RS-6228/2020 | EPI_ISL_729800 | 2020-05-28 | Laboratorio Central de Saude Publica do Estado do Rio Grande do Sul (LACEN-RS) | Laboratory of Respiratory Viruses and Measles, Oswaldo Cruz Institute, FIOCRUZ | Paola Resende et al |
| Brazil/RS-6232/2020 | EPI_ISL_729840 | 2020-05-28 | Laboratorio Central de Saude Publica do Estado do Rio Grande do Sul (LACEN-RS) | Laboratory of Respiratory Viruses and Measles, Oswaldo Cruz Institute, FIOCRUZ | Paola Resende et al |
| Brazil/RJ-0720R2/2020 | EPI_ISL_636837 | 2020-05-29 | Laboratório de Imunofarmacologia - Instituto Oswaldo Cruz | Laboratório de Imunofarmacologia - Instituto Oswaldo Cruz | Souza et al |
| Brazil/RS-6226/2020 | EPI_ISL_729846 | 2020-05-29 | Laboratorio Central de Saude Publica do Estado do Rio Grande do Sul (LACEN-RS) | Laboratory of Respiratory Viruses and Measles, Oswaldo Cruz Institute, FIOCRUZ | Paola Resende et al |
| Brazil/SP-673/2020 | EPI_ISL_693232 | 2020-05-30 | Hospital e Pronto Socorro Portinari | Instituto Adolfo Lutz, Interdiciplinary Procedures Center, Strategic Laboratory | Claudio Tavares Sacchi et al |
| Brazil/RJ-00524/2020 | EPI_ISL_717910 | 2020-05-31 | LACEN Dr. Francisco Rimolo Neto | Bioinformatics Laboratory / LNCC | Carolina M Voloch et al |
| Brazil/RS-6242/2020 | EPI_ISL_729843 | 2020-05-31 | Laboratorio Central de Saude Publica do Estado do Rio Grande do Sul (LACEN-RS) | Laboratory of Respiratory Viruses and Measles, Oswaldo Cruz Institute, FIOCRUZ | Paola Resende et al |
| Brazil/RJ-00410/2020 | EPI_ISL_717812 | 2020-06-01 | Laboratorio de Virologia Molecular / UFRJ | Bioinformatics Laboratory / LNCC | Carolina M Voloch et al |
| Brazil/RJ-00411/2020 | EPI_ISL_717813 | 2020-06-01 | Laboratorio de Virologia Molecular / UFRJ | Bioinformatics Laboratory / LNCC | Carolina M Voloch et al |
| Brazil/RJ-00450/2020 | EPI_ISL_717845 | 2020-06-01 | Laboratorio de Virologia Molecular / UFRJ | Bioinformatics Laboratory / LNCC | Carolina M Voloch et al |
| Brazil/RJ-00451/2020 | EPI_ISL_717846 | 2020-06-01 | Laboratorio de Virologia Molecular / UFRJ | Bioinformatics Laboratory / LNCC | Carolina M Voloch et al |
| Brazil/RJ-00455/2020 | EPI_ISL_717850 | 2020-06-01 | Laboratorio de Virologia Molecular / UFRJ | Bioinformatics Laboratory / LNCC | Carolina M Voloch et al |
| Brazil/RJ-00456/2020 | EPI_ISL_717851 | 2020-06-01 | Laboratorio de Virologia Molecular / UFRJ | Bioinformatics Laboratory / LNCC | Carolina M Voloch et al |
| Brazil/RJ-00457/2020 | EPI_ISL_717852 | 2020-06-01 | Laboratorio de Virologia Molecular / UFRJ | Bioinformatics Laboratory / LNCC | Carolina M Voloch et al |
| Brazil/RJ-00525/2020 | EPI_ISL_717911 | 2020-06-01 | LACEN Dr. Francisco Rimolo Neto | Bioinformatics Laboratory / LNCC | Carolina M Voloch et al |
| Brazil/RJ-UFRJ-9331/2020 | EPI_ISL_492036 | 2020-06-01 | Instituto de Biologia do Exército | Laboratório Metabolismo Macromolecular FirminoTorres de Castro, Instituto de Biofísica Carlos Chagas Filho, Universidade Federal do Rio de Janeiro | Bianca Catarina Azevedo Cabral et al |
| Brazil/RS-6240/2020 | EPI_ISL_729841 | 2020-06-01 | Laboratorio Central de Saude Publica do Estado do Rio Grande do Sul (LACEN-RS) | Laboratory of Respiratory Viruses and Measles, Oswaldo Cruz Institute, FIOCRUZ | Paola Resende et al |
| Brazil/RS-6241/2020 | EPI_ISL_729842 | 2020-06-01 | Laboratorio Central de Saude Publica do Estado do Rio Grande do Sul (LACEN-RS) | Laboratory of Respiratory Viruses and Measles, Oswaldo Cruz Institute, FIOCRUZ | Paola Resende et al |

| | | | | | |
|---|---|---|---|---|---|
| Brazil/SP-341/2020 | EPI_ISL_547571 | 2020-06-01 | Hospital Municipal Antônio Giglio | Instituto Adolfo Lutz, Interdiciplinary Procedures Center, Strategic Laboratory | Claudio Tavares Sacchi et al |
| Brazil/RS-6243/2020 | EPI_ISL_729847 | 2020-06-02 | Laboratorio Central de Saude Publica do Estado do Rio Grande do Sul (LACEN-RS) | Laboratory of Respiratory Viruses and Measles, Oswaldo Cruz Institute, FIOCRUZ | Paola Resende et al |
| Brazil/RJ-00438/2020 | EPI_ISL_717833 | 2020-06-03 | LACEN Dr. Francisco Rimolo Neto | Bioinformatics Laboratory / LNCC | Carolina M Voloch et al |
| Brazil/SP-376/2020 | EPI_ISL_603023 | 2020-06-07 | Vigilância em Saúde Visa Sul | Instituto Adolfo Lutz, Interdiciplinary Procedures Center, Strategic Laboratory | Claudio Tavares Sacchi et al |
| Brasil/RS11069/2020 | EPI_ISL_831474 | 2020-06-12 | Laboratório de Microbiologia Molecular - Universidade FEEVALE | Universidade Federal de Ciências da Saúde de Porto Alegre | Vinícius Bonetti Franceschi et al |
| Brazil/RS-11262/2020 | EPI_ISL_831645 | 2020-06-15 | Laboratório de Microbiologia Molecular - Universidade FEEVALE | Universidade Federal de Ciências da Saúde de Porto Alegre | Vinícius Bonetti Franceschi et al |
| Brazil/SP-356/2020 | EPI_ISL_547575 | 2020-06-16 | SVO Jundiaí | Instituto Adolfo Lutz, Interdiciplinary Procedures Center, Strategic Laboratory | Claudio Tavares Sacchi et al |
| Brazil/SP-666/2020 | EPI_ISL_693229 | 2020-06-16 | Hospital 8 de Maio | Instituto Adolfo Lutz, Interdiciplinary Procedures Center, Strategic Laboratory | Claudio Tavares Sacchi et al |
| Brazil/RS-11574/2020 | EPI_ISL_831660 | 2020-06-17 | Laboratório de Microbiologia Molecular - Universidade FEEVALE | Universidade Federal de Ciências da Saúde de Porto Alegre | Vinícius Bonetti Franceschi et al |
| Brazil/SP-389/2020 | EPI_ISL_574597 | 2020-06-18 | Secretaria Municipal de Saude de Jarinu | Instituto Adolfo Lutz, Interdiciplinary Procedures Center, Strategic Laboratory | Claudio Tavares Sacchi et al |
| Brazil/SP-353/2020 | EPI_ISL_583496 | 2020-06-20 | UPA Jandira | Instituto Adolfo Lutz, Interdiciplinary Procedures Center, Strategic Laboratory | Claudio Tavares Sacchi et al |
| Brazil/SP-383/2020 | EPI_ISL_603024 | 2020-06-20 | Santa Casa de Misericordia de Araçatuba | Instituto Adolfo Lutz, Interdiciplinary Procedures Center, Strategic Laboratory | Claudio Tavares Sacchi et al |
| Brazil/SP-361/2020 | EPI_ISL_693206 | 2020-06-21 | Hospital Municipal Mario Gatti | Instituto Adolfo Lutz, Interdiciplinary Procedures Center, Strategic Laboratory | Claudio Tavares Sacchi et al |
| Brazil/SP-393/2020 | EPI_ISL_583502 | 2020-06-21 | Serv de Vig Sanitaria Epidemio e CTRL de Zoonoses Guaruja | Instituto Adolfo Lutz, Interdiciplinary Procedures Center, Strategic Laboratory | Claudio Tavares Sacchi et al |
| Brazil/SP-394/2020 | EPI_ISL_583503 | 2020-06-22 | CTA Centro de Testagem e Aconselhamento | Instituto Adolfo Lutz, Interdiciplinary Procedures Center, Strategic Laboratory | Claudio Tavares Sacchi et al |
| Brazil/SP-577/2020 | EPI_ISL_693205 | 2020-06-22 | Hospital de Campanha Covid-19 Assis | Instituto Adolfo Lutz, Interdiciplinary Procedures Center, Strategic Laboratory | Claudio Tavares Sacchi et al |
| Brazil/PE-COV0260/2020 | EPI_ISL_502875 | 2020-06-24 | LACEN/PE | LABBE, Federal University of Pernambuco | WILSON JOSE DA SILVA JUNIOR et al |
| Brazil/SP-668/2020 | EPI_ISL_693231 | 2020-06-27 | Pronto Socorro Municipal de Santa Branca | Instituto Adolfo Lutz, Interdiciplinary Procedures Center, Strategic Laboratory | Claudio Tavares Sacchi et al |
| Brazil/SP-370/2020 | EPI_ISL_583500 | 2020-06-29 | Centro de Saude I Tacito Leite de Carvalho e Silva | Instituto Adolfo Lutz, Interdiciplinary Procedures Center, Strategic Laboratory | Claudio Tavares Sacchi et al |
| Brazil/SP-398/2020 | EPI_ISL_603028 | 2020-06-30 | Hospital Municipal Santa Ana | Instituto Adolfo Lutz, Interdiciplinary Procedures Center, Strategic Laboratory | Claudio Tavares Sacchi et al |
| Brazil/SP-600/2020 | EPI_ISL_515525 | 2020-06-30 | National Influenza Center - Instituto Adolfo Lutz | Instituto Adolfo Lutz, Interdiciplinary Procedures Center, Strategic Laboratory | Claudio Tavares Sacchi et al |
| Brazil/SP-688/2020 | EPI_ISL_693234 | 2020-06-30 | Upa Vereador Jose Da Rocha Goncalves | Instituto Adolfo Lutz, Interdiciplinary Procedures Center, Strategic Laboratory | Claudio Tavares Sacchi et al |
| Brazil/SP-696/2020 | EPI_ISL_693238 | 2020-06-30 | Secao Centro de Diagnostico Secedi | Instituto Adolfo Lutz, Interdiciplinary Procedures Center, Strategic Laboratory | Claudio Tavares Sacchi et al |
| Brazil/SP-400/2020 | EPI_ISL_693208 | 2020-07-01 | Hospital Municipal Antonio Giglio | Instituto Adolfo Lutz, Interdiciplinary Procedures Center, Strategic Laboratory | Claudio Tavares Sacchi et al |
| Brazil/SP-401/2020 | EPI_ISL_693209 | 2020-07-01 | Hospital Municipal Antonio Giglio | Instituto Adolfo Lutz, Interdiciplinary Procedures Center, Strategic Laboratory | Claudio Tavares Sacchi et al |
| Brazil/SP-694/2020 | EPI_ISL_693237 | 2020-07-01 | UPA Santa Isabel | Instituto Adolfo Lutz, Interdiciplinary Procedures Center, Strategic Laboratory | Claudio Tavares Sacchi et al |

| | | | | | |
|---|---|---|---|---|---|
| Brazil/RJ-00463/2020 | EPI_ISL_717858 | 2020-07-02 | Laboratorio de Virologia Molecular / UFRJ | Bioinformatics Laboratory / LNCC | Carolina M Voloch et al |
| Brazil/RS-13367/2020 | EPI_ISL_831681 | 2020-07-02 | Laboratório de Microbiologia Molecular - Universidade FEEVALE | Universidade Federal de Ciências da Saúde de Porto Alegre | Vinícius Bonetti Franceschi et al |
| Brazil/SP-692/2020 | EPI_ISL_693236 | 2020-07-02 | Hospital Santa Marcelina Sao Paulo | Instituto Adolfo Lutz, Interdiciplinary Procedures Center, Strategic Laboratory | Claudio Tavares Sacchi et al |
| Brazil/SP-719/2020 | EPI_ISL_693245 | 2020-07-02 | UPA Santa Isabel | Instituto Adolfo Lutz, Interdiciplinary Procedures Center, Strategic Laboratory | Claudio Tavares Sacchi et al |
| Brazil/SP-590/2020 | EPI_ISL_693216 | 2020-07-05 | Unidade de Vigilância Epiudemiológica de Araras | Instituto Adolfo Lutz, Interdiciplinary Procedures Center, Strategic Laboratory | Claudio Tavares Sacchi et al |
| Brazil/SP-591/2020 | EPI_ISL_693217 | 2020-07-05 | Unidade de Vigilância Epiudemiológica de Araras | Instituto Adolfo Lutz, Interdiciplinary Procedures Center, Strategic Laboratory | Claudio Tavares Sacchi et al |
| Brazil/SP-415/2020 | EPI_ISL_693210 | 2020-07-06 | Pronto-Socorro Dr. Osmar Mesquita | Instituto Adolfo Lutz, Interdiciplinary Procedures Center, Strategic Laboratory | Claudio Tavares Sacchi et al |
| Brazil/SP-589/2020 | EPI_ISL_693215 | 2020-07-06 | Secretaria Municipal de Saúde de Iracemapolis | Instituto Adolfo Lutz, Interdiciplinary Procedures Center, Strategic Laboratory | Claudio Tavares Sacchi et al |
| Brazil/SP-417/2020 | EPI_ISL_547579 | 2020-07-07 | Santa Casa de Misericórdia de Araçatuba | Instituto Adolfo Lutz, Interdiciplinary Procedures Center, Strategic Laboratory | Claudio Tavares Sacchi et al |
| Brazil/RJ-00466/2020 | EPI_ISL_717861 | 2020-07-08 | Laboratorio de Virologia Molecular / UFRJ | Bioinformatics Laboratory / LNCC | Carolina M Voloch et al |
| Brazil/RJ-00467/2020 | EPI_ISL_717862 | 2020-07-08 | Laboratorio de Virologia Molecular / UFRJ | Bioinformatics Laboratory / LNCC | Carolina M Voloch et al |
| Brazil/SP-425/2020 | EPI_ISL_574594 | 2020-07-10 | Hospital Escola da Universidade de Taubate | Instituto Adolfo Lutz, Interdiciplinary Procedures Center, Strategic Laboratory | Claudio Tavares Sacchi et al |
| Brazil/SP-438/2020 | EPI_ISL_603035 | 2020-07-10 | Secretaria Municipal de Saúde | Instituto Adolfo Lutz, Interdiciplinary Procedures Center, Strategic Laboratory | Claudio Tavares Sacchi et al |
| Brazil/SP-439/2020 | EPI_ISL_603036 | 2020-07-10 | Hospital Santa Ana | Instituto Adolfo Lutz, Interdiciplinary Procedures Center, Strategic Laboratory | Claudio Tavares Sacchi et al |
| Brazil/SP-433/2020 | EPI_ISL_603030 | 2020-07-11 | Hospital Domingos Leonardo Ceravolo Presidente Prudente | Instituto Adolfo Lutz, Interdiciplinary Procedures Center, Strategic Laboratory | Claudio Tavares Sacchi et al |
| Brazil/SP-437/2020 | EPI_ISL_603034 | 2020-07-11 | Departamento de Vigilância à Saúde | Instituto Adolfo Lutz, Interdiciplinary Procedures Center, Strategic Laboratory | Claudio Tavares Sacchi et al |
| Brazil/RS-15270/2020 | EPI_ISL_729801 | 2020-07-13 | Laboratorio Central de Saude Publica do Estado do Rio Grande do Sul (LACEN-RS) | Laboratory of Respiratory Viruses and Measles, Oswaldo Cruz Institute, FIOCRUZ | Paola Resende et al |
| Brazil/SP-436/2020 | EPI_ISL_603033 | 2020-07-13 | Vigilancia Epidemiologica de São Bernardo do Campo | Instituto Adolfo Lutz, Interdiciplinary Procedures Center, Strategic Laboratory | Claudio Tavares Sacchi et al |
| Brazil/RS-15371/2020 | EPI_ISL_831688 | 2020-07-15 | Laboratório de Microbiologia Molecular - Universidade FEEVALE | Universidade Federal de Ciências da Saúde de Porto Alegre | Vinícius Bonetti Franceschi et al |
| Brazil/SP-427/2020 | EPI_ISL_603021 | 2020-07-15 | Pronto Socorro Dr. Conrado Cesarino Nuvolini | Instituto Adolfo Lutz, Interdiciplinary Procedures Center, Strategic Laboratory | Claudio Tavares Sacchi et al |
| Brazil/SP-429/2020 | EPI_ISL_603022 | 2020-07-16 | Departamento de Vigilância à Saúde | Instituto Adolfo Lutz, Interdiciplinary Procedures Center, Strategic Laboratory | Claudio Tavares Sacchi et al |
| Brazil/RS-15273/2020 | EPI_ISL_729844 | 2020-07-17 | Laboratorio Central de Saude Publica do Estado do Rio Grande do Sul (LACEN-RS) | Laboratory of Respiratory Viruses and Measles, Oswaldo Cruz Institute, FIOCRUZ | Paola Resende et al |
| Brazil/RS-15274/2020 | EPI_ISL_729850 | 2020-07-17 | Laboratorio Central de Saude Publica do Estado do Rio Grande do Sul (LACEN-RS) | Laboratory of Respiratory Viruses and Measles, Oswaldo Cruz Institute, FIOCRUZ | Paola Resende et al |
| Brazil/RS-15275/2020 | EPI_ISL_729856 | 2020-07-18 | Laboratorio Central de Saude Publica do Estado do Rio Grande do Sul (LACEN-RS) | Laboratory of Respiratory Viruses and Measles, Oswaldo Cruz Institute, FIOCRUZ | Paola Resende et al |
| Brazil/SP-579/2020 | EPI_ISL_693214 | 2020-07-19 | Unidade de Pronto Atendimento Central de Caraguatatuba | Instituto Adolfo Lutz, Interdiciplinary Procedures Center, Strategic Laboratory | Claudio Tavares Sacchi et al |
| Brazil/RJ-DCV05/2020 | EPI_ISL_509430 | 2020-07-20 | Centro de Desenvolvimento Tecnologico em Saude, Fundacao Oswaldo Cruz | Centro de Desenvolvimento Tecnologico em Saude, Fundacao Oswaldo Cruz | Souza et al |
| Brazil/RJ-DCV1/2020 | EPI_ISL_509431 | 2020-07-20 | Centro de Desenvolvimento Tecnologico em Saude, Fundacao Oswaldo Cruz | Centro de Desenvolvimento Tecnologico em Saude, Fundacao Oswaldo Cruz | Souza et al |

| | | | | | |
|---|---|---|---|---|---|
| Brazil/RJ-DCV3/2020 | EPI_ISL_509432 | 2020-07-20 | Centro de Desenvolvimento Tecnologico em Saude, Fundacao Oswaldo Cruz | Centro de Desenvolvimento Tecnologico em Saude, Fundacao Oswaldo Cruz | Souza et al |
| Brazil/RJ-DCV5/2020 | EPI_ISL_509433 | 2020-07-20 | Centro de Desenvolvimento Tecnologico em Saude, Fundacao Oswaldo Cruz | Centro de Desenvolvimento Tecnologico em Saude, Fundacao Oswaldo Cruz | Souza et al |
| Brazil/RJ-DCVN2/2020 | EPI_ISL_529139 | 2020-07-20 | Centro de Desenvolvimento Tecnologico em Saude, Fundacao Oswaldo Cruz | Centro de Desenvolvimento Tecnologico em Saude, Fundacao Oswaldo Cruz | Souza et al |
| Brazil/RJ-DCVN3/2020 | EPI_ISL_529140 | 2020-07-20 | Centro de Desenvolvimento Tecnologico em Saude, Fundacao Oswaldo Cruz | Centro de Desenvolvimento Tecnologico em Saude, Fundacao Oswaldo Cruz | Souza et al |
| Brazil/RJ-DCVTV507Q/2020 | EPI_ISL_509435 | 2020-07-20 | Centro de Desenvolvimento Tecnologico em Saude, Fundacao Oswaldo Cruz | Centro de Desenvolvimento Tecnologico em Saude, Fundacao Oswaldo Cruz | Souza et al |
| Brazil/RS-15276/2020 | EPI_ISL_729857 | 2020-07-20 | Laboratorio Central de Saude Publica do Estado do Rio Grande do Sul (LACEN-RS) | Laboratory of Respiratory Viruses and Measles, Oswaldo Cruz Institute, FIOCRUZ | Paola Resende et al |
| Brazil/SP-440/2020 | EPI_ISL_603037 | 2020-07-20 | Hospital Geral de Pedreira | Instituto Adolfo Lutz, Interdiciplinary Procedures Center, Strategic Laboratory | Claudio Tavares Sacchi et al |
| Brazil/RS-15278/2020 | EPI_ISL_729848 | 2020-07-27 | Laboratorio Central de Saude Publica do Estado do Rio Grande do Sul (LACEN-RS) | Laboratory of Respiratory Viruses and Measles, Oswaldo Cruz Institute, FIOCRUZ | Paola Resende et al |
| Brazil/RJ-00406/2020 | EPI_ISL_717786 | 2020-07-28 | LACEN RJ - Noel Nutels | Bioinformatics Laboratory / LNCC | Carolina M Voloch et al |
| Brazil/RS-15279/2020 | EPI_ISL_729845 | 2020-07-28 | Laboratorio Central de Saude Publica do Estado do Rio Grande do Sul (LACEN-RS) | Laboratory of Respiratory Viruses and Measles, Oswaldo Cruz Institute, FIOCRUZ | Paola Resende et al |
| Brazil/RS-15280/2020 | EPI_ISL_729851 | 2020-07-28 | Laboratorio Central de Saude Publica do Estado do Rio Grande do Sul (LACEN-RS) | Laboratory of Respiratory Viruses and Measles, Oswaldo Cruz Institute, FIOCRUZ | Paola Resende et al |
| Brazil/RJ-00517/2020 | EPI_ISL_717906 | 2020-07-29 | LACEN RJ - Noel Nutels | Bioinformatics Laboratory / LNCC | Carolina M Voloch et al |
| Brazil/RJ-00521/2020 | EPI_ISL_717789 | 2020-07-29 | LACEN RJ - Noel Nutels | Bioinformatics Laboratory / LNCC | Carolina M Voloch et al |
| Brazil/RJ-00520/2020 | EPI_ISL_717908 | 2020-08-01 | LACEN RJ - Noel Nutels | Bioinformatics Laboratory / LNCC | Carolina M Voloch et al |
| Brazil/RJ-00468/2020 | EPI_ISL_717863 | 2020-08-03 | Laboratorio de Virologia Molecular / UFRJ | Bioinformatics Laboratory / LNCC | Carolina M Voloch et al |
| Brazil/RJ-00469/2020 | EPI_ISL_717864 | 2020-08-03 | Laboratorio de Virologia Molecular / UFRJ | Bioinformatics Laboratory / LNCC | Carolina M Voloch et al |
| Brazil/RJ-00470/2020 | EPI_ISL_717865 | 2020-08-03 | Laboratorio de Virologia Molecular / UFRJ | Bioinformatics Laboratory / LNCC | Carolina M Voloch et al |
| Brazil/RS-19337/2020 | EPI_ISL_831689 | 2020-08-03 | Laboratório de Microbiologia Molecular - Universidade FEEVALE | Universidade Federal de Ciências da Saúde de Porto Alegre | Vinícius Bonetti Franceschi et al |
| Brazil/RJ-00472/2020 | EPI_ISL_717867 | 2020-08-04 | Laboratorio de Virologia Molecular / UFRJ | Bioinformatics Laboratory / LNCC | Carolina M Voloch et al |
| Brazil/RJ-00473/2020 | EPI_ISL_717868 | 2020-08-05 | Laboratorio de Virologia Molecular / UFRJ | Bioinformatics Laboratory / LNCC | Carolina M Voloch et al |
| Brazil/RS-15281/2020 | EPI_ISL_729852 | 2020-08-07 | Laboratorio Central de Saude Publica do Estado do Rio Grande do Sul (LACEN-RS) | Laboratory of Respiratory Viruses and Measles, Oswaldo Cruz Institute, FIOCRUZ | Paola Resende et al |
| Brazil/RS-15283/2020 | EPI_ISL_729802 | 2020-08-09 | Laboratorio Central de Saude Publica do Estado do Rio Grande do Sul (LACEN-RS) | Laboratory of Respiratory Viruses and Measles, Oswaldo Cruz Institute, FIOCRUZ | Paola Resende et al |
| Brazil/RS-15284/2020 | EPI_ISL_729853 | 2020-08-12 | Laboratorio Central de Saude Publica do Estado do Rio Grande do Sul (LACEN-RS) | Laboratory of Respiratory Viruses and Measles, Oswaldo Cruz Institute, FIOCRUZ | Paola Resende et al |
| Brazil/RS-15288/2020 | EPI_ISL_729849 | 2020-08-12 | Laboratorio Central de Saude Publica do Estado do Rio Grande do Sul (LACEN-RS) | Laboratory of Respiratory Viruses and Measles, Oswaldo Cruz Institute, FIOCRUZ | Paola Resende et al |
| Brazil/RS-15287/2020 | EPI_ISL_729860 | 2020-08-13 | Laboratorio Central de Saude Publica do Estado do Rio Grande do Sul (LACEN-RS) | Laboratory of Respiratory Viruses and Measles, Oswaldo Cruz Institute, FIOCRUZ | Paola Resende et al |
| Brazil/RS-15285/2020 | EPI_ISL_729859 | 2020-08-14 | Laboratorio Central de Saude Publica do Estado do Rio Grande do Sul (LACEN-RS) | Laboratory of Respiratory Viruses and Measles, Oswaldo Cruz Institute, FIOCRUZ | Paola Resende et al |
| Brazil/RS-15286/2020 | EPI_ISL_729803 | 2020-08-14 | Laboratorio Central de Saude Publica do Estado do Rio Grande do Sul (LACEN-RS) | Laboratory of Respiratory Viruses and Measles, Oswaldo Cruz Institute, FIOCRUZ | Paola Resende et al |
| Brazil/RS-15289/2020 | EPI_ISL_729854 | 2020-08-14 | Laboratorio Central de Saude Publica do Estado do Rio Grande do Sul (LACEN-RS) | Laboratory of Respiratory Viruses and Measles, Oswaldo Cruz Institute, FIOCRUZ | Paola Resende et al |
| Brazil/RS-15290/2020 | EPI_ISL_729799 | 2020-08-15 | Laboratorio Central de Saude Publica do Estado do Rio Grande do Sul (LACEN-RS) | Laboratory of Respiratory Viruses and Measles, Oswaldo Cruz Institute, FIOCRUZ | Paola Resende et al |

| | | | | | |
|---|---|---|---|---|---|
| Brazil/RS-15291/2020 | EPI_ISL_729855 | 2020-08-15 | Laboratorio Central de Saude Publica do Estado do Rio Grande do Sul (LACEN-RS) | Laboratory of Respiratory Viruses and Measles, Oswaldo Cruz Institute, FIOCRUZ | Paola Resende et al |
| Brazil/SP-163/2020 | EPI_ISL_523959 | 2020-08-15 | Pronto Socorro Municipal de Perus | Instituto Adolfo Lutz, Interdiciplinary Procedures Center, Strategic Laboratory | Claudio Tavares Sacchi et al |
| Brazil/RS-15292/2020 | EPI_ISL_729861 | 2020-08-16 | Laboratorio Central de Saude Publica do Estado do Rio Grande do Sul (LACEN-RS) | Laboratory of Respiratory Viruses and Measles, Oswaldo Cruz Institute, FIOCRUZ | Paola Resende et al |
| Brazil/RS-22838/2020 | EPI_ISL_831892 | 2020-08-19 | Laboratório de Microbiologia Molecular - Universidade FEEVALE | Universidade Federal de Ciências da Saúde de Porto Alegre | Vinícius Bonetti Franceschi et al |
| Brazil/RS-24285/2020 | EPI_ISL_831913 | 2020-08-24 | Laboratório de Microbiologia Molecular - Universidade FEEVALE | Universidade Federal de Ciências da Saúde de Porto Alegre | Vinícius Bonetti Franceschi et al |
| Brazil/RS-24277/2020 | EPI_ISL_831898 | 2020-08-26 | Laboratório de Microbiologia Molecular - Universidade FEEVALE | Universidade Federal de Ciências da Saúde de Porto Alegre | Vinícius Bonetti Franceschi et al |
| Brazil/RS-24565/2020 | EPI_ISL_831938 | 2020-08-27 | Laboratório de Microbiologia Molecular - Universidade FEEVALE | Universidade Federal de Ciências da Saúde de Porto Alegre | Vinícius Bonetti Franceschi et al |
| Brazil/RJ-00402/2020 | EPI_ISL_717806 | 2020-09-03 | Laboratorio de Virologia Molecular / UFRJ | Bioinformatics Laboratory / LNCC | Carolina M Voloch et al |
| Brazil/RJ-00412/2020 | EPI_ISL_717814 | 2020-09-03 | Laboratorio de Virologia Molecular / UFRJ | Bioinformatics Laboratory / LNCC | Carolina M Voloch et al |
| Brazil/RJ-00413/2020 | EPI_ISL_717815 | 2020-09-03 | Laboratorio de Virologia Molecular / UFRJ | Bioinformatics Laboratory / LNCC | Carolina M Voloch et al |
| Brazil/RJ-00414/2020 | EPI_ISL_717816 | 2020-09-03 | Laboratorio de Virologia Molecular / UFRJ | Bioinformatics Laboratory / LNCC | Carolina M Voloch et al |
| Brazil/RJ-00474/2020 | EPI_ISL_717869 | 2020-09-03 | Laboratorio de Virologia Molecular / UFRJ | Bioinformatics Laboratory / LNCC | Carolina M Voloch et al |
| Brazil/RS-25833/2020 | EPI_ISL_831939 | 2020-09-03 | Laboratório de Microbiologia Molecular - Universidade FEEVALE | Universidade Federal de Ciências da Saúde de Porto Alegre | Vinícius Bonetti Franceschi et al |
| Brazil/RJ-00415/2020 | EPI_ISL_717817 | 2020-09-04 | Laboratorio de Virologia Molecular / UFRJ | Bioinformatics Laboratory / LNCC | Carolina M Voloch et al |
| Brazil/RJ-00475/2020 | EPI_ISL_717870 | 2020-09-04 | Laboratorio de Virologia Molecular / UFRJ | Bioinformatics Laboratory / LNCC | Carolina M Voloch et al |
| Brazil/RJ-00476/2020 | EPI_ISL_717871 | 2020-09-04 | Laboratorio de Virologia Molecular / UFRJ | Bioinformatics Laboratory / LNCC | Carolina M Voloch et al |
| Brazil/RS-26977/2020 | EPI_ISL_831940 | 2020-09-10 | Laboratório de Microbiologia Molecular - Universidade FEEVALE | Universidade Federal de Ciências da Saúde de Porto Alegre | Vinícius Bonetti Franceschi et al |
| Brazil/RS-27623/2020 | EPI_ISL_832009 | 2020-09-16 | Laboratório de Microbiologia Molecular - Universidade FEEVALE | Universidade Federal de Ciências da Saúde de Porto Alegre | Vinícius Bonetti Franceschi et al |
| Brazil/RJ-00407/2020 | EPI_ISL_717809 | 2020-09-28 | LACEN Dr. Francisco Rimolo Neto | Bioinformatics Laboratory / LNCC | Carolina M Voloch et al |
| Brazil/PB-23855-R2/2020 | EPI_ISL_792562 | 2020-10-13 | LACEN-PB | Laboratory of Respiratory Viruses and Measles, Oswaldo Cruz Institute, FIOCRUZ | Paola Resende et al |
| Brazil/RS-31786/2020 | EPI_ISL_832011 | 2020-10-13 | Laboratório de Microbiologia Molecular - Universidade FEEVALE | Universidade Federal de Ciências da Saúde de Porto Alegre | Vinícius Bonetti Franceschi et al |
| Brazil/BA-2/2020 | EPI_ISL_756294 | 2020-10-26 | Center for Biotechnology and Cell Therapy, São Rafael Hospital, Salvador, Brazil | Center for Biotechnology and Cell Therapy, São Rafael Hospital, Salvador, Brazil | Carolina Kymie Vasques Nonaka et al |
| Brazil/RJ-00542/2020 | EPI_ISL_717926 | 2020-10-26 | Laboratorio de Virologia Molecular / UFRJ | Bioinformatics Laboratory / LNCC | Carolina M Voloch et al |
| Brazil/PR-28601/2020 | EPI_ISL_792645 | 2020-10-27 | LACEN-PR | Laboratory of Respiratory Viruses and Measles, Oswaldo Cruz Institute, FIOCRUZ | Paola Resende et al |
| Brazil/RJ-00543/2020 | EPI_ISL_717927 | 2020-10-27 | Laboratorio de Virologia Molecular / UFRJ | Bioinformatics Laboratory / LNCC | Carolina M Voloch et al |
| Brazil/RJ-00544/2020 | EPI_ISL_717928 | 2020-10-27 | Laboratorio de Virologia Molecular / UFRJ | Bioinformatics Laboratory / LNCC | Carolina M Voloch et al |
| Brazil/RJ-00545/2020 | EPI_ISL_717929 | 2020-10-27 | Laboratorio de Virologia Molecular / UFRJ | Bioinformatics Laboratory / LNCC | Carolina M Voloch et al |
| Brazil/RJ-00546/2020 | EPI_ISL_717930 | 2020-10-27 | Laboratorio de Virologia Molecular / UFRJ | Bioinformatics Laboratory / LNCC | Carolina M Voloch et al |
| Brazil/RJ-00547/2020 | EPI_ISL_717931 | 2020-10-27 | Laboratorio de Virologia Molecular / UFRJ | Bioinformatics Laboratory / LNCC | Carolina M Voloch et al |
| Brazil/RJ-00537/2020 | EPI_ISL_717922 | 2020-10-29 | Laboratorio de Virologia Molecular / UFRJ | Bioinformatics Laboratory / LNCC | Carolina M Voloch et al |
| Brazil/RJ-00540/2020 | EPI_ISL_717924 | 2020-10-29 | Laboratorio de Virologia Molecular / UFRJ | Bioinformatics Laboratory / LNCC | Carolina M Voloch et al |
| Brazil/RJ-00548/2020 | EPI_ISL_717932 | 2020-10-29 | Laboratorio de Virologia Molecular / UFRJ | Bioinformatics Laboratory / LNCC | Carolina M Voloch et al |

| | | | | | |
|---|---|---|---|---|---|
| Brazil/RJ-00549/2020 | EPI_ISL_717933 | 2020-10-29 | Laboratorio de Virologia Molecular / UFRJ | Bioinformatics Laboratory / LNCC | Carolina M Voloch et al |
| Brazil/RJ-00550/2020 | EPI_ISL_717934 | 2020-10-29 | Laboratorio de Virologia Molecular / UFRJ | Bioinformatics Laboratory / LNCC | Carolina M Voloch et al |
| Brazil/RJ-00551/2020 | EPI_ISL_717935 | 2020-10-29 | Laboratorio de Virologia Molecular / UFRJ | Bioinformatics Laboratory / LNCC | Carolina M Voloch et al |
| Brazil/RJ-00552/2020 | EPI_ISL_717936 | 2020-10-29 | Laboratorio de Virologia Molecular / UFRJ | Bioinformatics Laboratory / LNCC | Carolina M Voloch et al |
| Brazil/RJ-00553/2020 | EPI_ISL_717937 | 2020-10-29 | Laboratorio de Virologia Molecular / UFRJ | Bioinformatics Laboratory / LNCC | Carolina M Voloch et al |
| Brazil/RJ-00554/2020 | EPI_ISL_717938 | 2020-10-29 | Laboratorio de Virologia Molecular / UFRJ | Bioinformatics Laboratory / LNCC | Carolina M Voloch et al |
| Brazil/RJ-00555/2020 | EPI_ISL_717939 | 2020-10-29 | Laboratorio de Virologia Molecular / UFRJ | Bioinformatics Laboratory / LNCC | Carolina M Voloch et al |
| Brazil/RJ-00556/2020 | EPI_ISL_717940 | 2020-10-30 | Laboratorio de Virologia Molecular / UFRJ | Bioinformatics Laboratory / LNCC | Carolina M Voloch et al |
| Brazil/RJ-00557/2020 | EPI_ISL_717941 | 2020-10-30 | Laboratorio de Virologia Molecular / UFRJ | Bioinformatics Laboratory / LNCC | Carolina M Voloch et al |
| Brazil/PR-28602/2020 | EPI_ISL_792646 | 2020-11-03 | LACEN-PR | Laboratory of Respiratory Viruses and Measles, Oswaldo Cruz Institute, FIOCRUZ | Paola Resende et al |
| Brazil/RJ-00405/2020 | EPI_ISL_717808 | 2020-11-03 | Laboratorio de Virologia Molecular / UFRJ | Bioinformatics Laboratory / LNCC | Carolina M Voloch et al |
| Brazil/RJ-00558/2020 | EPI_ISL_717942 | 2020-11-03 | Laboratorio de Virologia Molecular / UFRJ | Bioinformatics Laboratory / LNCC | Carolina M Voloch et al |
| Brazil/RJ-00559/2020 | EPI_ISL_717943 | 2020-11-03 | Laboratorio de Virologia Molecular / UFRJ | Bioinformatics Laboratory / LNCC | Carolina M Voloch et al |
| Brazil/RJ-00560/2020 | EPI_ISL_717944 | 2020-11-03 | Laboratorio de Virologia Molecular / UFRJ | Bioinformatics Laboratory / LNCC | Carolina M Voloch et al |
| Brazil/RJ-00561/2020 | EPI_ISL_717945 | 2020-11-03 | Laboratorio de Virologia Molecular / UFRJ | Bioinformatics Laboratory / LNCC | Carolina M Voloch et al |
| Brazil/RJ-00562/2020 | EPI_ISL_717946 | 2020-11-03 | Laboratorio de Virologia Molecular / UFRJ | Bioinformatics Laboratory / LNCC | Carolina M Voloch et al |
| Brazil/RJ-00563/2020 | EPI_ISL_717947 | 2020-11-03 | Laboratorio de Virologia Molecular / UFRJ | Bioinformatics Laboratory / LNCC | Carolina M Voloch et al |
| Brazil/RJ-00564/2020 | EPI_ISL_717948 | 2020-11-03 | Laboratorio de Virologia Molecular / UFRJ | Bioinformatics Laboratory / LNCC | Carolina M Voloch et al |
| Brazil/RJ-00565/2020 | EPI_ISL_717949 | 2020-11-03 | Laboratorio de Virologia Molecular / UFRJ | Bioinformatics Laboratory / LNCC | Carolina M Voloch et al |
| Brazil/RJ-00566/2020 | EPI_ISL_717950 | 2020-11-03 | Laboratorio de Virologia Molecular / UFRJ | Bioinformatics Laboratory / LNCC | Carolina M Voloch et al |
| Brazil/RJ-00567/2020 | EPI_ISL_717951 | 2020-11-04 | Laboratorio de Virologia Molecular / UFRJ | Bioinformatics Laboratory / LNCC | Carolina M Voloch et al |
| Brazil/RJ-00568/2020 | EPI_ISL_717952 | 2020-11-04 | Laboratorio de Virologia Molecular / UFRJ | Bioinformatics Laboratory / LNCC | Carolina M Voloch et al |
| Brazil/RJ-00569/2020 | EPI_ISL_717953 | 2020-11-04 | Laboratorio de Virologia Molecular / UFRJ | Bioinformatics Laboratory / LNCC | Carolina M Voloch et al |
| Brazil/RJ-00570/2020 | EPI_ISL_717954 | 2020-11-04 | Laboratorio de Virologia Molecular / UFRJ | Bioinformatics Laboratory / LNCC | Carolina M Voloch et al |
| Brazil/RJ-00428/2020 | EPI_ISL_717830 | 2020-11-05 | Laboratorio de Virologia Molecular / UFRJ | Bioinformatics Laboratory / LNCC | Carolina M Voloch et al |
| Brazil/RJ-00504/2020 | EPI_ISL_717898 | 2020-11-05 | Laboratorio de Virologia Molecular / UFRJ | Bioinformatics Laboratory / LNCC | Carolina M Voloch et al |
| Brazil/RJ-00571/2020 | EPI_ISL_717955 | 2020-11-05 | Laboratorio de Virologia Molecular / UFRJ | Bioinformatics Laboratory / LNCC | Carolina M Voloch et al |
| Brazil/RJ-00572/2020 | EPI_ISL_717956 | 2020-11-05 | Laboratorio de Virologia Molecular / UFRJ | Bioinformatics Laboratory / LNCC | Carolina M Voloch et al |
| Brazil/RJ-00573/2020 | EPI_ISL_717957 | 2020-11-05 | Laboratorio de Virologia Molecular / UFRJ | Bioinformatics Laboratory / LNCC | Carolina M Voloch et al |
| Brazil/AM-20142410MC/2020 | EPI_ISL_792560 | 2020-11-06 | Laboratorio de Ecologia de Doencas Transmissiveis na Amazonia, Instituto Leonidas e Maria Deane - Fiocruz Amazonia | Laboratorio de Ecologia de Doencas Transmissiveis na Amazonia, Instituto Leonidas e Maria Deane - Fiocruz Amazonia | Valdinete Nascimento et al |
| Brazil/PB-29646/2020 | EPI_ISL_792634 | 2020-11-10 | LACEN-PB | Laboratory of Respiratory Viruses and Measles, Oswaldo Cruz Institute, FIOCRUZ | Paola Resende et al |
| Brazil/PB-29647/2020 | EPI_ISL_792635 | 2020-11-10 | LACEN-PB | Laboratory of Respiratory Viruses and Measles, Oswaldo Cruz Institute, FIOCRUZ | Paola Resende et al |
| Brazil/SP-931/2020 | EPI_ISL_833163 | 2020-11-19 | Instituto Adolfo Lutz - Regional de Marilia | Instituto Adolfo Lutz, Interdiciplinary Procedures Center, Strategic Laboratory | Claudio Tavares Sacchi et al |
| Brazil/AL-28256/2020 | EPI_ISL_792639 | 2020-11-21 | LACEN-AL | Laboratory of Respiratory Viruses and Measles, Oswaldo Cruz Institute, FIOCRUZ | Paola Resende et al |
| Brazil/SP-844R2/2020 | EPI_ISL_708530 | 2020-11-21 | Secretaria Municipal de Saude de Fernandópolis | Instituto Adolfo Lutz, Interdiciplinary Procedures Center, Strategic Laboratory | Claudio Tavares Sacchi et al |

| | | | | | |
|---|---|---|---|---|---|
| Brazil/AL-28875/2020 | EPI_ISL_792641 | 2020-11-23 | LACEN-AL | Laboratory of Respiratory Viruses and Measles, Oswaldo Cruz Institute, FIOCRUZ | Paola Resende et al |
| Brazil/PR-28606/2020 | EPI_ISL_792650 | 2020-11-23 | LACEN-PR | Laboratory of Respiratory Viruses and Measles, Oswaldo Cruz Institute, FIOCRUZ | Paola Resende et al |
| Brazil/PR-28607/2020 | EPI_ISL_792651 | 2020-11-23 | LACEN-PR | Laboratory of Respiratory Viruses and Measles, Oswaldo Cruz Institute, FIOCRUZ | Paola Resende et al |
| Brazil/PR-28610/2020 | EPI_ISL_792654 | 2020-11-23 | LACEN-PR | Laboratory of Respiratory Viruses and Measles, Oswaldo Cruz Institute, FIOCRUZ | Paola Resende et al |
| Brazil/RS-00601/2020 | EPI_ISL_779155 | 2020-11-23 | Laboratório de Microbiologia Molecular - Universidade FEEVALE | Bioinformatics Laboratory / LNCC | Felipe Benites et al |
| Brazil/RS-00602/2020 | EPI_ISL_779156 | 2020-11-23 | Laboratório de Microbiologia Molecular - Universidade FEEVALE | Bioinformatics Laboratory / LNCC | Felipe Benites et al |
| Brazil/RS-00603/2020 | EPI_ISL_770555 | 2020-11-23 | Laboratório de Microbiologia Molecular - Universidade FEEVALE | Bioinformatics Laboratory / LNCC | Felipe Benites et al |
| Brazil/RS-00604/2020 | EPI_ISL_770556 | 2020-11-23 | Laboratório de Microbiologia Molecular - Universidade FEEVALE | Bioinformatics Laboratory / LNCC | Felipe Benites et al |
| Brazil/RS-00622/2020 | EPI_ISL_770573 | 2020-11-23 | Laboratório de Microbiologia Molecular - Universidade FEEVALE | Bioinformatics Laboratory / LNCC | Felipe Benites et al |
| Brazil/RS-00625/2020 | EPI_ISL_770575 | 2020-11-23 | Laboratório de Microbiologia Molecular - Universidade FEEVALE | Bioinformatics Laboratory / LNCC | Felipe Benites et al |
| Brazil/RS-00626/2020 | EPI_ISL_770576 | 2020-11-23 | Laboratório de Microbiologia Molecular - Universidade FEEVALE | Bioinformatics Laboratory / LNCC | Felipe Benites et al |
| Brazil/RS-00605/2020 | EPI_ISL_770557 | 2020-11-24 | Laboratório de Microbiologia Molecular - Universidade FEEVALE | Bioinformatics Laboratory / LNCC | Felipe Benites et al |
| Brazil/RS-00606/2020 | EPI_ISL_770558 | 2020-11-24 | Laboratório de Microbiologia Molecular - Universidade FEEVALE | Bioinformatics Laboratory / LNCC | Felipe Benites et al |
| Brazil/RS-00607/2020 | EPI_ISL_770559 | 2020-11-24 | Laboratório de Microbiologia Molecular - Universidade FEEVALE | Bioinformatics Laboratory / LNCC | Felipe Benites et al |
| Brazil/RS-00608/2020 | EPI_ISL_770560 | 2020-11-24 | Laboratório de Microbiologia Molecular - Universidade FEEVALE | Bioinformatics Laboratory / LNCC | Felipe Benites et al |
| Brazil/RS-00609/2020 | EPI_ISL_770561 | 2020-11-24 | Laboratório de Microbiologia Molecular - Universidade FEEVALE | Bioinformatics Laboratory / LNCC | Felipe Benites et al |
| Brazil/RS-00611/2020 | EPI_ISL_770563 | 2020-11-24 | Laboratório de Microbiologia Molecular - Universidade FEEVALE | Bioinformatics Laboratory / LNCC | Felipe Benites et al |
| Brazil/RS-00612/2020 | EPI_ISL_770564 | 2020-11-24 | Laboratório de Microbiologia Molecular - Universidade FEEVALE | Bioinformatics Laboratory / LNCC | Felipe Benites et al |
| Brazil/RS-00614/2020 | EPI_ISL_770566 | 2020-11-24 | Laboratório de Microbiologia Molecular - Universidade FEEVALE | Bioinformatics Laboratory / LNCC | Felipe Benites et al |
| Brazil/RS-00618/2020 | EPI_ISL_770570 | 2020-11-24 | Laboratório de Microbiologia Molecular - Universidade FEEVALE | Bioinformatics Laboratory / LNCC | Felipe Benites et al |
| Brazil/RS-00620/2020 | EPI_ISL_770572 | 2020-11-24 | Laboratório de Microbiologia Molecular - Universidade FEEVALE | Bioinformatics Laboratory / LNCC | Felipe Benites et al |
| Brazil/RS-00627/2020 | EPI_ISL_770577 | 2020-11-24 | Laboratório de Microbiologia Molecular - Universidade FEEVALE | Bioinformatics Laboratory / LNCC | Felipe Benites et al |
| Brazil/RS-00628/2020 | EPI_ISL_770553 | 2020-11-24 | Laboratório de Microbiologia Molecular - Universidade FEEVALE | Bioinformatics Laboratory / LNCC | Felipe Benites et al |
| Brazil/RS-00629/2020 | EPI_ISL_770578 | 2020-11-24 | Laboratório de Microbiologia Molecular - Universidade FEEVALE | Bioinformatics Laboratory / LNCC | Felipe Benites et al |
| Brazil/RS-00633/2020 | EPI_ISL_770581 | 2020-11-24 | Laboratório de Microbiologia Molecular - Universidade FEEVALE | Bioinformatics Laboratory / LNCC | Felipe Benites et al |
| Brazil/RS-00634/2020 | EPI_ISL_770582 | 2020-11-24 | Laboratório de Microbiologia Molecular - Universidade FEEVALE | Bioinformatics Laboratory / LNCC | Felipe Benites et al |

| | | | | | |
|---|---|---|---|---|---|
| Brazil/RS-00636/2020 | EPI_ISL_770584 | 2020-11-24 | Laboratório de Microbiologia Molecular - Universidade FEEVALE | Bioinformatics Laboratory / LNCC | Felipe Benites et al |
| Brazil/RS-00637/2020 | EPI_ISL_770585 | 2020-11-24 | Laboratório de Microbiologia Molecular - Universidade FEEVALE | Bioinformatics Laboratory / LNCC | Felipe Benites et al |
| Brazil/RS-00640/2020 | EPI_ISL_770588 | 2020-11-24 | Laboratório de Microbiologia Molecular - Universidade FEEVALE | Bioinformatics Laboratory / LNCC | Felipe Benites et al |
| Brazil/RS-00653/2020 | EPI_ISL_770598 | 2020-11-24 | Laboratório de Microbiologia Molecular - Universidade FEEVALE | Bioinformatics Laboratory / LNCC | Felipe Benites et al |
| Brazil/PR-28608/2020 | EPI_ISL_792652 | 2020-11-25 | LACEN-PR | Laboratory of Respiratory Viruses and Measles, Oswaldo Cruz Institute, FIOCRUZ | Paola Resende et al |
| Brazil/RS-00613/2020 | EPI_ISL_770565 | 2020-11-25 | Laboratório de Microbiologia Molecular - Universidade FEEVALE | Bioinformatics Laboratory / LNCC | Felipe Benites et al |
| Brazil/RS-00615/2020 | EPI_ISL_770567 | 2020-11-25 | Laboratório de Microbiologia Molecular - Universidade FEEVALE | Bioinformatics Laboratory / LNCC | Felipe Benites et al |
| Brazil/RS-00616/2020 | EPI_ISL_770568 | 2020-11-25 | Laboratório de Microbiologia Molecular - Universidade FEEVALE | Bioinformatics Laboratory / LNCC | Felipe Benites et al |
| Brazil/RS-00619/2020 | EPI_ISL_770571 | 2020-11-25 | Laboratório de Microbiologia Molecular - Universidade FEEVALE | Bioinformatics Laboratory / LNCC | Felipe Benites et al |
| Brazil/RS-00630/2020 | EPI_ISL_770579 | 2020-11-25 | Laboratório de Microbiologia Molecular - Universidade FEEVALE | Bioinformatics Laboratory / LNCC | Felipe Benites et al |
| Brazil/RS-00631/2020 | EPI_ISL_779159 | 2020-11-25 | Laboratório de Microbiologia Molecular - Universidade FEEVALE | Bioinformatics Laboratory / LNCC | Felipe Benites et al |
| Brazil/RS-00632/2020 | EPI_ISL_770580 | 2020-11-25 | Laboratório de Microbiologia Molecular - Universidade FEEVALE | Bioinformatics Laboratory / LNCC | Felipe Benites et al |
| Brazil/RS-00635/2020 | EPI_ISL_770583 | 2020-11-25 | Laboratório de Microbiologia Molecular - Universidade FEEVALE | Bioinformatics Laboratory / LNCC | Felipe Benites et al |
| Brazil/RS-00638/2020 | EPI_ISL_770586 | 2020-11-25 | Laboratório de Microbiologia Molecular - Universidade FEEVALE | Bioinformatics Laboratory / LNCC | Felipe Benites et al |
| Brazil/RS-00639/2020 | EPI_ISL_770587 | 2020-11-25 | Laboratório de Microbiologia Molecular - Universidade FEEVALE | Bioinformatics Laboratory / LNCC | Felipe Benites et al |
| Brazil/RS-00642/2020 | EPI_ISL_770589 | 2020-11-25 | Laboratório de Microbiologia Molecular - Universidade FEEVALE | Bioinformatics Laboratory / LNCC | Felipe Benites et al |
| Brazil/RS-00643/2020 | EPI_ISL_770590 | 2020-11-25 | Laboratório de Microbiologia Molecular - Universidade FEEVALE | Bioinformatics Laboratory / LNCC | Felipe Benites et al |
| Brazil/RS-00645/2020 | EPI_ISL_770591 | 2020-11-25 | Laboratório de Microbiologia Molecular - Universidade FEEVALE | Bioinformatics Laboratory / LNCC | Felipe Benites et al |
| Brazil/RS-00646/2020 | EPI_ISL_770592 | 2020-11-25 | Laboratório de Microbiologia Molecular - Universidade FEEVALE | Bioinformatics Laboratory / LNCC | Felipe Benites et al |
| Brazil/RS-00647/2020 | EPI_ISL_770593 | 2020-11-25 | Laboratório de Microbiologia Molecular - Universidade FEEVALE | Bioinformatics Laboratory / LNCC | Felipe Benites et al |
| Brazil/RS-00648/2020 | EPI_ISL_770594 | 2020-11-25 | Laboratório de Microbiologia Molecular - Universidade FEEVALE | Bioinformatics Laboratory / LNCC | Felipe Benites et al |
| Brazil/RS-00649/2020 | EPI_ISL_770595 | 2020-11-25 | Laboratório de Microbiologia Molecular - Universidade FEEVALE | Bioinformatics Laboratory / LNCC | Felipe Benites et al |
| Brazil/RS-00650/2020 | EPI_ISL_770596 | 2020-11-25 | Laboratório de Microbiologia Molecular - Universidade FEEVALE | Bioinformatics Laboratory / LNCC | Felipe Benites et al |
| Brazil/RS-00652/2020 | EPI_ISL_770597 | 2020-11-25 | Laboratório de Microbiologia Molecular - Universidade FEEVALE | Bioinformatics Laboratory / LNCC | Felipe Benites et al |
| Brazil/RS-00655/2020 | EPI_ISL_770552 | 2020-11-25 | Laboratório de Microbiologia Molecular - Universidade FEEVALE | Bioinformatics Laboratory / LNCC | Felipe Benites et al |
| Brazil/RS-00659/2020 | EPI_ISL_770602 | 2020-11-26 | Laboratório de Microbiologia Molecular - Universidade FEEVALE | Bioinformatics Laboratory / LNCC | Felipe Benites et al |

| | | | | | |
|---|---|---|---|---|---|
| Brazil/RS-00660/2020 | EPI_ISL_770603 | 2020-11-26 | Laboratório de Microbiologia Molecular - Universidade FEEVALE | Bioinformatics Laboratory / LNCC | Felipe Benites et al |
| Brazil/RS-00661/2020 | EPI_ISL_770604 | 2020-11-26 | Laboratório de Microbiologia Molecular - Universidade FEEVALE | Bioinformatics Laboratory / LNCC | Felipe Benites et al |
| Brazil/RS-00662/2020 | EPI_ISL_770605 | 2020-11-26 | Laboratório de Microbiologia Molecular - Universidade FEEVALE | Bioinformatics Laboratory / LNCC | Felipe Benites et al |
| Brazil/RS-00663/2020 | EPI_ISL_770606 | 2020-11-26 | Laboratório de Microbiologia Molecular - Universidade FEEVALE | Bioinformatics Laboratory / LNCC | Felipe Benites et al |
| Brazil/RS-00665/2020 | EPI_ISL_770607 | 2020-11-26 | Laboratório de Microbiologia Molecular - Universidade FEEVALE | Bioinformatics Laboratory / LNCC | Felipe Benites et al |
| Brazil/RS-00676/2020 | EPI_ISL_770617 | 2020-11-26 | Laboratório de Microbiologia Molecular - Universidade FEEVALE | Bioinformatics Laboratory / LNCC | Felipe Benites et al |
| Brazil/SP-870/2020 | EPI_ISL_776766 | 2020-11-26 | Instituto Adolfo Lutz - Regional de Santo Andre | Instituto Adolfo Lutz, Interdiciplinary Procedures Center, Strategic Laboratory | Claudio Tavares Sacchi et al |
| Brazil/SP-923/2020 | EPI_ISL_833161 | 2020-11-26 | Instituto Adolfo Lutz - Central | Instituto Adolfo Lutz, Interdiciplinary Procedures Center, Strategic Laboratory | Claudio Tavares Sacchi et al |
| Brazil/PB-29608/2020 | EPI_ISL_792603 | 2020-11-27 | LACEN-PB | Laboratory of Respiratory Viruses and Measles, Oswaldo Cruz Institute, FIOCRUZ | Paola Resende et al |
| Brazil/RS-00674/2020 | EPI_ISL_770615 | 2020-11-27 | Laboratório de Microbiologia Molecular - Universidade FEEVALE | Bioinformatics Laboratory / LNCC | Felipe Benites et al |
| Brazil/RS-00675/2020 | EPI_ISL_770616 | 2020-11-27 | Laboratório de Microbiologia Molecular - Universidade FEEVALE | Bioinformatics Laboratory / LNCC | Felipe Benites et al |
| Brazil/RS-00679/2020 | EPI_ISL_770618 | 2020-11-27 | Laboratório de Microbiologia Molecular - Universidade FEEVALE | Bioinformatics Laboratory / LNCC | Felipe Benites et al |
| Brazil/RS-00680/2020 | EPI_ISL_770619 | 2020-11-27 | Laboratório de Microbiologia Molecular - Universidade FEEVALE | Bioinformatics Laboratory / LNCC | Felipe Benites et al |
| Brazil/RS-00681/2020 | EPI_ISL_770620 | 2020-11-27 | Laboratório de Microbiologia Molecular - Universidade FEEVALE | Bioinformatics Laboratory / LNCC | Felipe Benites et al |
| Brazil/RS-00682/2020 | EPI_ISL_770554 | 2020-11-27 | Laboratório de Microbiologia Molecular - Universidade FEEVALE | Bioinformatics Laboratory / LNCC | Felipe Benites et al |
| Brazil/RS-00683/2020 | EPI_ISL_770621 | 2020-11-27 | Laboratório de Microbiologia Molecular - Universidade FEEVALE | Bioinformatics Laboratory / LNCC | Felipe Benites et al |
| Brazil/RS-00684/2020 | EPI_ISL_770622 | 2020-11-27 | Laboratório de Microbiologia Molecular - Universidade FEEVALE | Bioinformatics Laboratory / LNCC | Felipe Benites et al |
| Brazil/RS-00686/2020 | EPI_ISL_770624 | 2020-11-27 | Laboratório de Microbiologia Molecular - Universidade FEEVALE | Bioinformatics Laboratory / LNCC | Felipe Benites et al |
| Brazil/RS-00689/2020 | EPI_ISL_770625 | 2020-11-27 | Laboratório de Microbiologia Molecular - Universidade FEEVALE | Bioinformatics Laboratory / LNCC | Felipe Benites et al |
| Brazil/SP-871/2020 | EPI_ISL_776767 | 2020-11-27 | Instituto Adolfo Lutz - Regional de Marilia | Instituto Adolfo Lutz, Interdiciplinary Procedures Center, Strategic Laboratory | Claudio Tavares Sacchi et al |
| Brazil/RS-00692/2020 | EPI_ISL_770628 | 2020-11-28 | Laboratório de Microbiologia Molecular - Universidade FEEVALE | Bioinformatics Laboratory / LNCC | Felipe Benites et al |
| Brazil/SP-924/2020 | EPI_ISL_833162 | 2020-11-28 | Lab LOC - Itapecerica da Serra | Instituto Adolfo Lutz, Interdiciplinary Procedures Center, Strategic Laboratory | Claudio Tavares Sacchi et al |
| Brazil/SP-873/2020 | EPI_ISL_776769 | 2020-11-30 | Instituto Adolfo Lutz - Regional de Santo Andre | Instituto Adolfo Lutz, Interdiciplinary Procedures Center, Strategic Laboratory | Claudio Tavares Sacchi et al |
| Brazil/SP-911/2020 | EPI_ISL_833158 | 2020-11-30 | Instituto Adolfo Lutz - Regional de Santo Andre | Instituto Adolfo Lutz, Interdiciplinary Procedures Center, Strategic Laboratory | Claudio Tavares Sacchi et al |
| Brazil/AM-20142761FC/2020 | EPI_ISL_833131 | 2020-12-01 | Laboratorio de Ecologia de Doencas Transmissiveis na Amazonia, Instituto Leonidas e Maria Deane - Fiocruz Amazonia | Laboratorio de Ecologia de Doencas Transmissiveis na Amazonia, Instituto Leonidas e Maria Deane - Fiocruz Amazonia | Valdinete Nascimento et al |

| | | | | | |
|---|---|---|---|---|---|
| Brazil/SP-872/2020 | EPI_ISL_776768 | 2020-12-01 | Instituto Adolfo Lutz - Regional de Aracatuba | Instituto Adolfo Lutz, Interdiciplinary Procedures Center, Strategic Laboratory | Claudio Tavares Sacchi et al |
| Brazil/AL-30270/2020 | EPI_ISL_792642 | 2020-12-04 | LACEN-AL | Laboratory of Respiratory Viruses and Measles, Oswaldo Cruz Institute, FIOCRUZ | Paola Resende et al |
| Brazil/AM-20842882CA/2020 | EPI_ISL_833137 | 2020-12-04 | Laboratorio de Ecologia de Doencas Transmissiveis na Amazonia, Instituto Leonidas e Maria Deane - Fiocruz Amazonia | Laboratorio de Ecologia de Doencas Transmissiveis na Amazonia, Instituto Leonidas e Maria Deane - Fiocruz Amazonia | Valdinete Nascimento et al |
| Brazil/SP-876/2020 | EPI_ISL_755642 | 2020-12-05 | Instituto Adolfo Lutz - Central | Instituto Adolfo Lutz, Interdiciplinary Procedures Center, Strategic Laboratory | Claudio Tavares Sacchi et al |
| Brazil/SP-883/2020 | EPI_ISL_755649 | 2020-12-06 | Instituto Adolfo Lutz - Regional de Santo Andre | Instituto Adolfo Lutz, Interdiciplinary Procedures Center, Strategic Laboratory | Claudio Tavares Sacchi et al |
| Brazil/SP-877/2020 | EPI_ISL_755643 | 2020-12-07 | Instituto Adolfo Lutz - Central | Instituto Adolfo Lutz, Interdiciplinary Procedures Center, Strategic Laboratory | Claudio Tavares Sacchi et al |
| Brazil/SP-912/2020 | EPI_ISL_836143 | 2020-12-07 | Hospital de Campanha COVID-19 de Mairipora | Instituto Adolfo Lutz, Interdiciplinary Procedures Center, Strategic Laboratory | Claudio Tavares Sacchi et al |
| Brazil/SP-879/2020 | EPI_ISL_755645 | 2020-12-08 | Lab LOC - Itapecerica da Serra | Instituto Adolfo Lutz, Interdiciplinary Procedures Center, Strategic Laboratory | Claudio Tavares Sacchi et al |
| Brazil/SP-915/2020 | EPI_ISL_836977 | 2020-12-08 | Hospital Municiplal Dr. Jose de Carvalho Florence | Instituto Adolfo Lutz, Interdiciplinary Procedures Center, Strategic Laboratory | Claudio Tavares Sacchi et al |
| Brazil/SP-886/2020 | EPI_ISL_755652 | 2020-12-09 | Lab LOC - Itapecerica da Serra | Instituto Adolfo Lutz, Interdiciplinary Procedures Center, Strategic Laboratory | Claudio Tavares Sacchi et al |
| Brazil/SP-918/2020 | EPI_ISL_837054 | 2020-12-09 | UBS Jose Sabino Ferreira | Instituto Adolfo Lutz, Interdiciplinary Procedures Center, Strategic Laboratory | Claudio Tavares Sacchi et al |
| Brazil/AM-20142910VS/2020 | EPI_ISL_833132 | 2020-12-10 | Laboratorio de Ecologia de Doencas Transmissiveis na Amazonia, Instituto Leonidas e Maria Deane - Fiocruz Amazonia | Laboratorio de Ecologia de Doencas Transmissiveis na Amazonia, Instituto Leonidas e Maria Deane - Fiocruz Amazonia | Valdinete Nascimento et al |
| Brazil/SP-885/2020 | EPI_ISL_755651 | 2020-12-10 | Instituto Adolfo Lutz - Central | Instituto Adolfo Lutz, Interdiciplinary Procedures Center, Strategic Laboratory | Claudio Tavares Sacchi et al |
| Brazil/SP-888/2020 | EPI_ISL_755654 | 2020-12-11 | Instituto Adolfo Lutz - Central | Instituto Adolfo Lutz, Interdiciplinary Procedures Center, Strategic Laboratory | Claudio Tavares Sacchi et al |
| Brazil/SP-887/2020 | EPI_ISL_755653 | 2020-12-12 | Instituto Adolfo Lutz - Central | Instituto Adolfo Lutz, Interdiciplinary Procedures Center, Strategic Laboratory | Claudio Tavares Sacchi et al |
| Brazil/AM-L70-CD1719/2020 | EPI_ISL_804827 | 2020-12-15 | DB Diagnosticos do Brasil | Laboratório de Parasitologia Médica - Instituto de Medicina Tropical - Universidade de São Paulo | Nuno Faria et al |
| Brazil/AM-987/2020 | EPI_ISL_833167 | 2020-12-16 | DB Diagnosticos do Brasil | Instituto Adolfo Lutz, Interdiciplinary Procedures Center, Strategic Laboratory | Claudio Tavares Sacchi et al |
| Brazil/AM-L70-CD1716/2020 | EPI_ISL_804815 | 2020-12-16 | DB Diagnosticos do Brasil | Laboratório de Parasitologia Médica - Instituto de Medicina Tropical - Universidade de São Paulo | Nuno Faria et al |
| Brazil/AM-L70-CD1718/2020 | EPI_ISL_804819 | 2020-12-16 | DB Diagnosticos do Brasil | Laboratório de Parasitologia Médica - Instituto de Medicina Tropical - Universidade de São Paulo | Nuno Faria et al |
| Brazil/AM-991/2020 | EPI_ISL_833171 | 2020-12-17 | DB Diagnosticos do Brasil | Instituto Adolfo Lutz, Interdiciplinary Procedures Center, Strategic Laboratory | Claudio Tavares Sacchi et al |
| Brazil/AM-992/2020 | EPI_ISL_833172 | 2020-12-17 | DB Diagnosticos do Brasil | Instituto Adolfo Lutz, Interdiciplinary Procedures Center, Strategic Laboratory | Claudio Tavares Sacchi et al |
| Brazil/AM-997/2020 | EPI_ISL_833176 | 2020-12-17 | DB Diagnosticos do Brasil | Instituto Adolfo Lutz, Interdiciplinary Procedures Center, Strategic Laboratory | Claudio Tavares Sacchi et al |
| Brazil/AM-L70-CD1721/2020 | EPI_ISL_804824 | 2020-12-17 | DB Diagnosticos do Brasil | Laboratório de Parasitologia Médica - Instituto de Medicina Tropical - Universidade de São Paulo | Nuno Faria et al |

| | | | | | |
|---|---|---|---|---|---|
| Brazil/AM-L70-CD1722/2020 | EPI_ISL_804823 | 2020-12-17 | DB Diagnosticos do Brasil | Laboratório de Parasitologia Médica - Instituto de Medicina Tropical - Universidade de São Paulo | Nuno Faria et al |
| Brazil/AM-L70-CD1726/2020 | EPI_ISL_804828 | 2020-12-17 | DB Diagnosticos do Brasil | Laboratório de Parasitologia Médica - Instituto de Medicina Tropical - Universidade de São Paulo | Nuno Faria et al |
| Brazil/AM-20143164MG/2020 | EPI_ISL_833133 | 2020-12-18 | Laboratorio de Ecologia de Doencas Transmissiveis na Amazonia, Instituto Leonidas e Maria Deane - Fiocruz Amazonia | Laboratorio de Ecologia de Doencas Transmissiveis na Amazonia, Instituto Leonidas e Maria Deane - Fiocruz Amazonia | Valdinete Nascimento et al |
| Brazil/AM-993/2020 | EPI_ISL_833173 | 2020-12-18 | DB Diagnosticos do Brasil | Instituto Adolfo Lutz, Interdiciplinary Procedures Center, Strategic Laboratory | Claudio Tavares Sacchi et al |
| Brazil/AM-L70-CD1728/2020 | EPI_ISL_804829 | 2020-12-18 | DB Diagnosticos do Brasil | Laboratório de Parasitologia Médica - Instituto de Medicina Tropical - Universidade de São Paulo | Nuno Faria et al |
| Brazil/AM-20843201ML/2020 | EPI_ISL_833138 | 2020-12-21 | Laboratorio de Ecologia de Doencas Transmissiveis na Amazonia, Instituto Leonidas e Maria Deane - Fiocruz Amazonia | Laboratorio de Ecologia de Doencas Transmissiveis na Amazonia, Instituto Leonidas e Maria Deane - Fiocruz Amazonia | Valdinete Nascimento et al |
| Brazil/AM-L70-CD1733/2020 | EPI_ISL_804821 | 2020-12-21 | DB Diagnosticos do Brasil | Laboratório de Parasitologia Médica - Instituto de Medicina Tropical - Universidade de São Paulo | Nuno Faria et al |
| Brazil/AM-L70-CD1737/2020 | EPI_ISL_804814 | 2020-12-21 | DB Diagnosticos do Brasil | Laboratório de Parasitologia Médica - Instituto de Medicina Tropical - Universidade de São Paulo | Nuno Faria et al |
| Brazil/AM-L70-CD1739/2020 | EPI_ISL_804820 | 2020-12-21 | DB Diagnosticos do Brasil | Laboratório de Parasitologia Médica - Instituto de Medicina Tropical - Universidade de São Paulo | Nuno Faria et al |
| Brazil/AM-20843228ZC/2020 | EPI_ISL_833139 | 2020-12-22 | Laboratorio de Ecologia de Doencas Transmissiveis na Amazonia, Instituto Leonidas e Maria Deane - Fiocruz Amazonia | Laboratorio de Ecologia de Doencas Transmissiveis na Amazonia, Instituto Leonidas e Maria Deane - Fiocruz Amazonia | Valdinete Nascimento et al |
| Brazil/AM-994/2020 | EPI_ISL_833174 | 2020-12-22 | DB Diagnosticos do Brasil | Instituto Adolfo Lutz, Interdiciplinary Procedures Center, Strategic Laboratory | Claudio Tavares Sacchi et al |
| Brazil/AM-996/2020 | EPI_ISL_833175 | 2020-12-22 | DB Diagnosticos do Brasil | Instituto Adolfo Lutz, Interdiciplinary Procedures Center, Strategic Laboratory | Claudio Tavares Sacchi et al |
| Brazil/AM-L71-CD1742/2020 | EPI_ISL_804816 | 2020-12-22 | DB Diagnosticos do Brasil | Laboratório de Parasitologia Médica - Instituto de Medicina Tropical - Universidade de São Paulo | Nuno Faria et al |
| Brazil/AM-20843269RC/2020 | EPI_ISL_833140 | 2020-12-23 | Laboratorio de Ecologia de Doencas Transmissiveis na Amazonia, Instituto Leonidas e Maria Deane - Fiocruz Amazonia | Laboratorio de Ecologia de Doencas Transmissiveis na Amazonia, Instituto Leonidas e Maria Deane - Fiocruz Amazonia | Valdinete Nascimento et al |
| Brazil/AM-988/2020 | EPI_ISL_833168 | 2020-12-23 | DB Diagnosticos do Brasil | Instituto Adolfo Lutz, Interdiciplinary Procedures Center, Strategic Laboratory | Claudio Tavares Sacchi et al |
| Brazil/AM-989/2020 | EPI_ISL_833169 | 2020-12-23 | DB Diagnosticos do Brasil | Instituto Adolfo Lutz, Interdiciplinary Procedures Center, Strategic Laboratory | Claudio Tavares Sacchi et al |
| Brazil/AM-990/2020 | EPI_ISL_833170 | 2020-12-23 | DB Diagnosticos do Brasil | Instituto Adolfo Lutz, Interdiciplinary Procedures Center, Strategic Laboratory | Claudio Tavares Sacchi et al |
| Brazil/AM-20143103JT/2020 | EPI_ISL_833136 | 2020-12-29 | Laboratorio de Ecologia de Doencas Transmissiveis na Amazonia, Instituto Leonidas e Maria Deane - Fiocruz Amazonia | Laboratorio de Ecologia de Doencas Transmissiveis na Amazonia, Instituto Leonidas e Maria Deane - Fiocruz Amazonia | Valdinete Nascimento et al |
| Brazil/AM-20143138FN-R2/2020 | EPI_ISL_811149 | 2020-12-30 | Laboratorio de Ecologia de Doencas Transmissiveis na Amazonia, Instituto Leonidas e Maria Deane - Fiocruz Amazonia | Laboratorio de Ecologia de Doencas Transmissiveis na Amazonia, Instituto Leonidas e Maria Deane - Fiocruz Amazonia | Valdinete Nascimento et al |

All submitters of data may be contacted directly via www.gisaid.org
Shu Y., McCauley, J. (2017) GISAID: from vision to reality EuroSurveillance 22(13) doi:10.2807/1560-7917.ES.2017.22.13.30494 PMCID: PMC5388101
